# SARS-CoV-2 and HSV-1 Induce Amyloid Aggregation in Human CSF Resulting in Drastic Soluble Protein Depletion

**DOI:** 10.1101/2022.09.15.508120

**Authors:** Wanda Christ, Sebastian Kapell, Michal J. Sobkowiak, Georgios Mermelekas, Björn Evertsson, Helena Sork, Osama Saher, Safa Bazaz, Oskar Gustafsson, Eduardo I. Cardenas, Viviana Villa, Roberta Ricciarelli, Johan K. Sandberg, Jonas Bergquist, Andrea Sturchio, Per Svenningsson, Tarja Malm, Alberto J. Espay, Maria Pernemalm, Anders Lindén, Jonas Klingström, Samir El Andaloussi, Kariem Ezzat

## Abstract

The corona virus (SARS-CoV-2) pandemic and the resulting long-term neurological complications in patients, known as long COVID, have renewed the interest in the correlation between viral infections and neurodegenerative brain disorders. While many viruses can reach the central nervous system (CNS) causing acute or chronic infections (such as herpes simplex virus 1, HSV-1), the lack of a clear mechanistic link between viruses and protein aggregation into amyloids, a characteristic of several neurodegenerative diseases, has rendered such a connection elusive. Recently, we showed that viruses can induce aggregation of purified amyloidogenic proteins via the direct physicochemical mechanism of heterogenous nucleation (HEN). In the current study, we show that the incubation of HSV-1 and SARS-CoV-2 with human cerebrospinal fluid (CSF) leads to the amyloid aggregation of several proteins known to be involved in neurodegenerative diseases, such as: APLP1 (amyloid beta precursor like protein 1), ApoE, clusterin, α2-macroglobulin, PGK-1 (phosphoglycerate kinase 1), ceruloplasmin, nucleolin, 14-3-3, transthyretin and vitronectin. Importantly, UV-inactivation of SARS-CoV-2 does not affect its ability to induce amyloid aggregation, as amyloid formation is dependent on viral surface catalysis via HEN and not its ability to replicate. Additionally, viral amyloid induction led to a dramatic drop in the soluble protein concentration in the CSF. Our results show that viruses can physically induce amyloid aggregation of proteins in human CSF and result in soluble protein depletion, and thus providing a potential mechanism that may account for the association between persistent and latent/reactivating brain infections and neurodegenerative diseases.

**Significance Statement:** Viruses have generally been excluded from the etiology of amyloid pathologies based on the assumption that amyloid formation requires a proteinaceous conformational template (a prion) to form. Here we show that neuroinvasive viruses induce amyloid aggregation of a plethora of proteins in human CSF even after UV inactivation. Our work illustrates that viruses can induce amyloid aggregation of endogenous human proteins in their native environment by acting as physical catalysts of amyloid nucleation and phase transition. Demonstrating this direct mechanistic link, which is independent of templating, can help better understand the link between viruses and neurodegenerative disorders, especially in the post-COVID-19 era.

## Introduction

The COVID-19 pandemic is estimated to have caused more than 6 million deaths worldwide in addition to tremendous economic and societal disruptions. While lock-down measures and vaccines helped ameliorate the acute impact of the pandemic, millions of people continue to suffer from post-acute COVID syndrome, or what is commonly known as long COVID (1). Many of the symptoms associated with long COVID are neurological in nature, such as fatigue, frequent headaches, and so-called “brain fog”, which includes difficulties with memory and concentration (2–4). The large and increasing number of people with post-COVID neurological symptoms has renewed the interest in the link between viruses and neurodegenerative brain disorders. There have been similar findings with other post-viral syndromes, including herpes simplex virus 1 (HSV-1) and its link to Alzheimer’s disease (AD) (5) and likewise for influenza virus and Parkinson’s disease (PD) (6). More recently, a strong link has been found between Epstein-Barr virus and multiple sclerosis (7). However, several factors have complicated the understanding of the connection between viruses and neurodegeneration, including:

1. The prevalence of viral infection is usually much higher than the prevalence of neurodegenerative diseases. For example, while nearly two-thirds of the human population have been infected with HSV-1 (8), the prevalence of AD is less than 2% (9).
2. The latent or persistent nature of viral brain infections, which usually involves alternating cycles of latency and activation, together with the long course of neurodegenerative diseases make it difficult to establish accurate temporal causal links.
3. The lack of a mechanism linking viral infections to amyloid aggregates, a hallmark of neurodegenerative disease.

The first two factors relate to the nature of viral infections of the brain and its immune responses to viruses. Since excessive inflammation can lead to irreparable neuronal damage, the brain evolved as an “immune-privileged” organ, where immune responses are tightly balanced and limited in magnitude and duration to prevent tissue damage (10, 11). Thus, while viral access to the brain is generally restricted by physical barriers (skull and meninges) and physiological barriers (the blood-brain-barrier), the viruses that manage to reach the brain benefit from the immune privilege and establish long-term latent (non-replicating) or persistent (low-level replication) infections, which can be reactivated depending on the genetic background and the immune status of the host (12). Furthermore, the non-replicating or low-level-replication of latent or persistent infections makes it very difficult to accurately quantify viral presence in the brain. However, the differential reactivation patterns across individuals may explain the prevalence gap between viral infections and neurodegenerative diseases, where some people are more susceptible than others to the long-term effects of chronic infections. Additionally, more sensitive detection methods are helping to better spatially and temporally correlate viruses with particular neurodegenerative pathologies. For example, RNAscope, a highly sensitive in-situ RNA hybridization method, has recently been utilized to differentiate between latent and lytic transcripts in human brain tissue and in the brains of AD mouse models infected with HSV-1, a result that was not attainable by normal qPCR (13). The same technique was used to show that SARS-CoV-2 infects cortical neurons in the brains of a subgroup of COVID-19 patients and induces AD-like neuropathology, including amyloid aggregation (14). Additionally, the proximity of the appearance of neurological symptoms to SARS-CoV-2 infection and the availability of large datasets from patients are enabling better causal correlations to be made. This has been demonstrated recently in a study of UK biobank participants, which identified significant longitudinal effects of SARS-CoV-2 infection on the brains of infected individuals, such as a reduction in grey matter thickness and global brain volume, compared to controls (15).

The third factor that put into question viral involvement in neurodegenerative diseases was the early finding that brain material inactivated by exposure to ultraviolet (UV) light can induce amyloid aggregation and neurodegeneration when inoculated into naive brains (16, 17). Since UV light inactivates nucleic acids, the agent inducing amyloid protein aggregation was hypothesized to be non-viral in nature, protein-only (prion), since viruses require nucleic acids for propagation (18). Protein-only prions are hypothesized to carry the conformational information required to template the transformation of normal proteins into amyloids (19). However, nanoparticles have been shown to trigger amyloid aggregation in the absence of a protein seed acting as a conformational template (20). Additionally, HSV-1 has been shown to triggers amyloid aggregation in the brain in a mouse model of AD (21) and we have demonstrated earlier that viruses, including HSV-1 and respiratory syncytial virus (RSV), can induce amyloid aggregation of purified amyloidogenic proteins via the physicochemical mechanism of heterogenous nucleation (HEN) (22). Recently, we outlined a theoretical framework that governs this process, illustrating that amyloid formation is a spontaneous folding event that is thermodynamically favorable under certain conditions, which do not rely on conformational templating (23). We have also recently demonstrated that, due to the highly stable and inert nature of amyloids, toxicity in amyloid pathologies might be more dependent on protein sequestration in the insoluble amyloid form (loss-of-function toxicity) rather than direct toxicity from the aggregates (gain-of-function toxicity) (23–25).

In the current study, we show that HSV-1 and UV-inactivated SARS-CoV-2 induce amyloid aggregation of proteins in human cerebrospinal fluid (CSF) ex-vivo. Using mass spectrometry-based proteomics to analyze the virus-induced amyloids, we identify several proteins that have been previously found in plaques from patients with neurodegenerative diseases. Our results indicate that viruses are mechanistically capable of directly triggering amyloid aggregation of proteins in human CSF.

## Results

### HSV-1 and SARS-CoV-2 induce protein aggregation in the CSF

To determine whether viral particles can catalyze amyloid aggregation of proteins in a complex human biofluid at their native concentration, we incubated live HSV-1 virus and UV-inactivated SARS-CoV-2 virus with CSF harvested from healthy individuals. We traced the ability of the viruses to induce amyloid formation using the thioflavin-T (ThT)-assay, where ThT fluorescence is enhanced upon binding to amyloid fibrils (26). Both viruses were able to induce amyloid aggregation of proteins in the CSF, while CSF with no virus added did not display any substantial signal for increasing amyloid aggregation over time (Figure 1. A and B). Additional controls, including non-infected cell medium and virus-only, produced a much lower signal, which is similar to what we reported earlier with purified amyloidogenic proteins (22). The maximum ThT fluorescence enhancement obtained from the viral treated CSF was significantly higher than all the controls (Supplementary Figure 1.). Furthermore, we using 3x-concentrated HSV-1 stock results in a higher amyloid signal (Supplementary Figure 2.), indicating that the interaction is concentration dependent.

**Figure 1.**
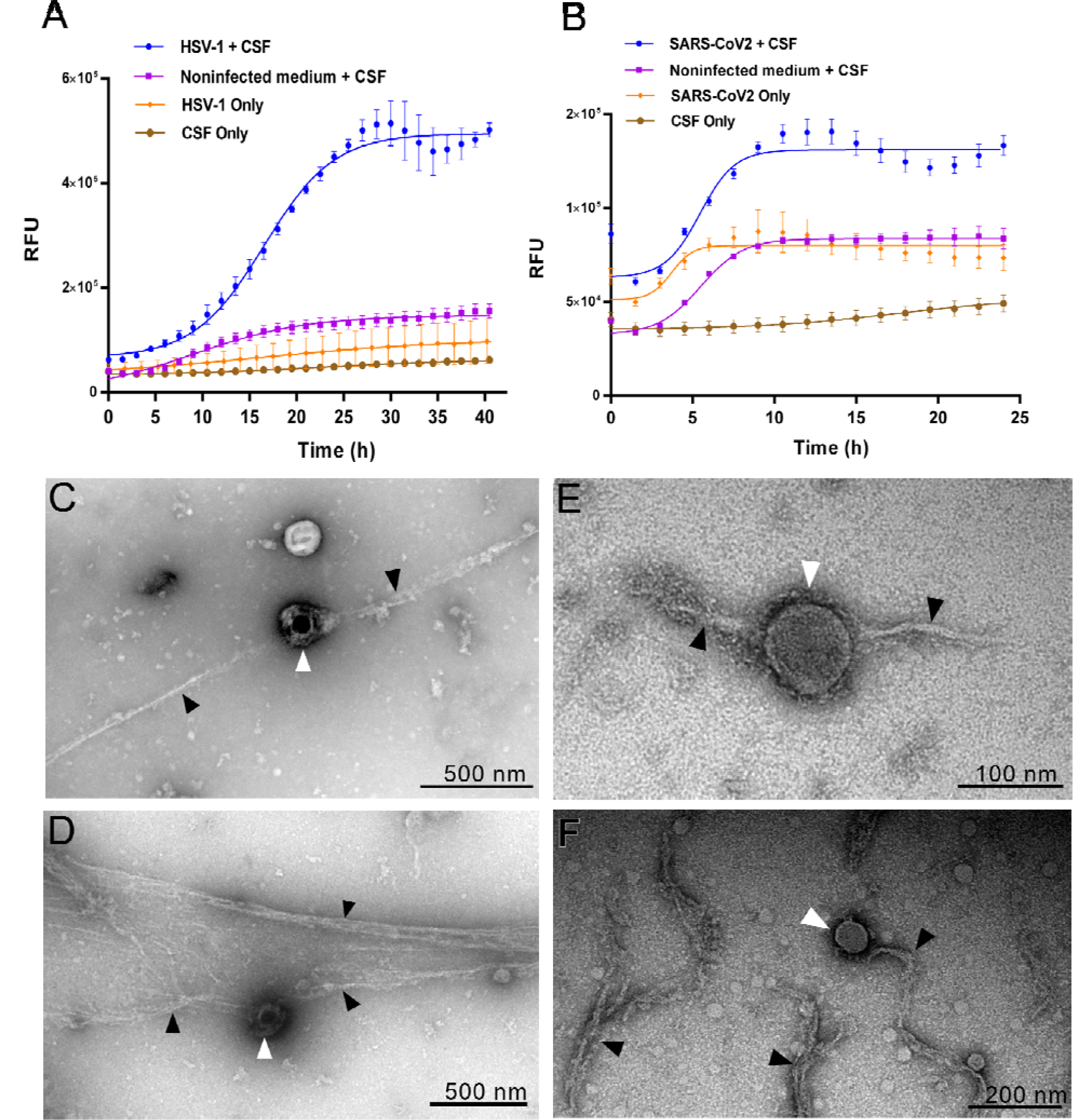
HSV-1 and SARS-CoV-2 induce amyloid aggregation of proteins in human CSF. CSF was incubated with HSV-1(**A**) or UV-inactivated-SARS-CoV-2 (**B**) and ThT solution. Fluorescence was measured at 440 nm excitation an 480 nm emission over 48 h at 37 °C. Means ± SEM of 8 replicates with CSF from two different individuals are shown. RFU = relative fluorescent unit. Negatively stained TEM images of HSV-1 (**C, D**) or UV-inactivated-SARS-CoV-2 (**E, F**) HSV-1 incubated with CSF for 48 h at 37 °C. White arrows indicate viral particles and black arrows indicate amyloid fibrillar structures.

**Figure 2.**
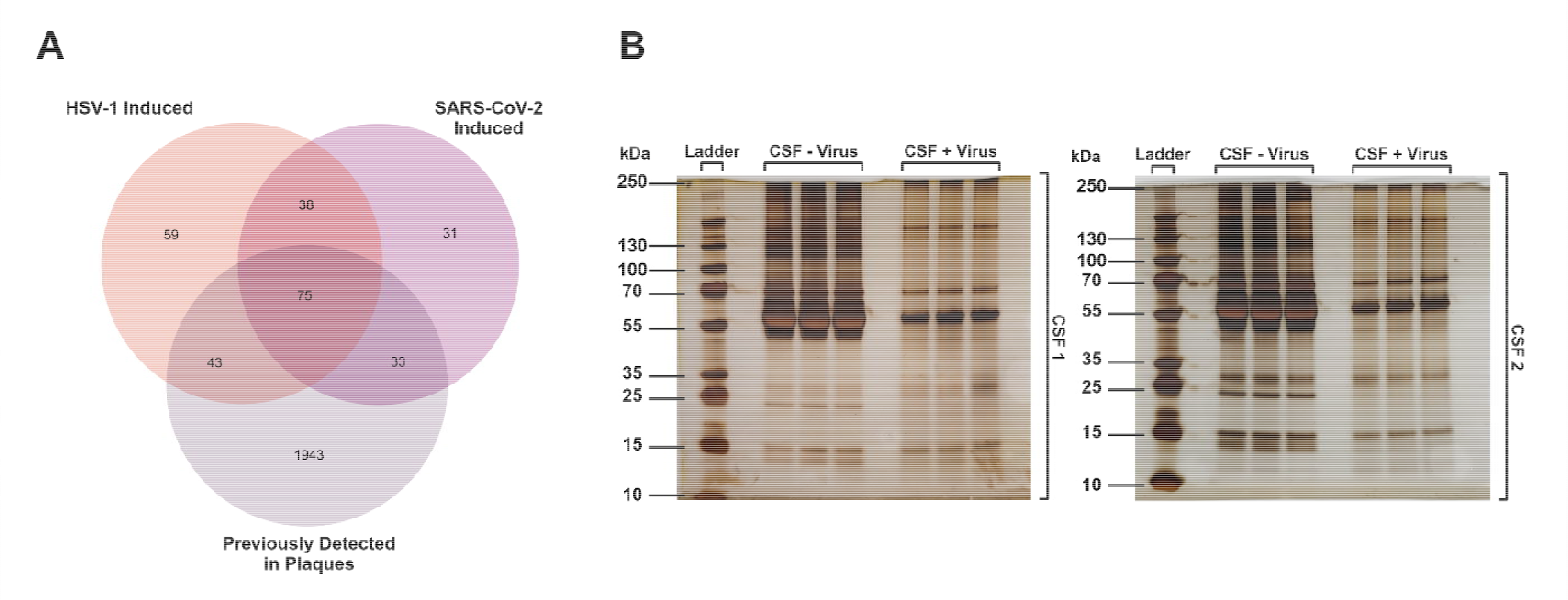
Viral particles induce the aggregation of overlapping sets of proteins and result in soluble protein depletion in the CSF A. Venn Diagram showing the overlap between the proteins detected in the HSV-1-induced amyloid fraction, the SARS-CoV-2-induced amyloid fraction in the amyloids fractions proteins detected before in amyloid plaques in AD patients (from Drummond E. et al., Acta Neuropathol 2017 (31)). **B**. CSF from two different individuals (CSF 1 and CSF 2) was incubated with UV-inactivated HSV-1 for 72 h at 37 °C. After removing the amyloid fraction by centrifugation, the soluble fraction was run on a polyacrylamide gel (3 replicates/CSF) and stained with silver staining.

In addition, we used transmission electron microscopy (TEM) to visualize amyloid formation induced by the viruses in the CSF. We found multiple fibrillar amyloid structures interacting at the surface of HSV-1 (Figure 1. C & D) and SARS-CoV-2 (Figure 1. E & F), suggestive of surface-mediated catalytic nucleation (HEN) events. We also include all the raw electron microcopy pictures in Supplementary Figure 3 for UV-inactivated SARS-CoV-2 + CSF & Supplementary Figure 4 for HSV-1 + CSF.

### Proteomic analysis of the viral-induced amyloid aggregates

To characterize the proteins present in the amyloid fractions, we collected and purified the amyloid aggregates induced by the viruses. We applied a protocol that involved using 4% SDS to remove the associated non-amyloid proteins. While some amyloid enrichment protocols use another detergent (sarkosyl), which is less stringent than SDS in order to preserve less robust aggregates that cannot withstand SDS treatment (27), we used a high concentration of SDS to ensure that have the highest possible purity in our amyloid fraction (28, 29). SDS treatment was followed by washing, centrifugation, then solubilization of the amyloid fraction in 99% formic acid, which is among the few reagents that can solubilize the highly stable amyloid structures (30). In the proteomic analysis, we found 279 proteins enriched in the virus-induced amyloid fractions, where about 40% (n= 113) of the enriched proteins were shared in the amyloid fractions induced by both viruses, while about 37% (n= 102) were unique for the HSV-1-induced amyloid fraction, and 23% (n= 64) unique for the SARS-CoV-2-induced amyloid fraction (Figure 2A & Supplementary Table 1.). The large overlap between the sets of proteins enriched by the two viruses indicates that certain proteins in the CSF are more vulnerable to aggregation catalysts. Previous proteomic studies have shown that a multitude of proteins can be found in amyloid plaques harvested from patients (29, 31). We found 151 proteins (Figure 2A & Supplementary table 2) previously reported in plaques harvested from AD patients (31), including established components such as APLP1 (amyloid beta precursor like protein 1), ApoE, clusterin and α2-macroglobulin. Moreover, we identified proteins that have been linked t PD (32) such as ceruloplasmin, nucleolin, 14-3-3 and PGK-1 (phosphoglycerate kinase 1), and proteins related to other amyloid pathologies including transthyretin (TTR) and vitronectin (33). These results demonstrate that viral particles can efficiently catalyze the amyloid aggregation of a plethora of proteins in human CSF.

Furthermore, to demonstrate the effect of viral-induced amyloid aggregation on the soluble protein fraction in CSF, we incubated the CSF with UV-inactivated HSV-1 viral particles to induce amyloid aggregation, and then removed the formed amyloids using centrifugation. Upon staining for proteins in the soluble fraction we found that the viral-incubated CSF had a dramatic reduction in the concentration of a large percentage of soluble proteins compared to the control incubated with non-infected cell medium (Figure 2B). These results clearly demonstrate that viral-induced amyloid aggregation depletes the CSF of a large fraction of its soluble proteome, a potential loss-of-function mechanism of pathogenesis.

## Discussion

While viruses have long been implicated in neurodegenerative diseases, it has been difficult to establish a mechanistic link. However, the large number of patients developing post-COVID neurological symptoms and the extensive data available from studying COVID-19 patients are starting to shed light on the possible causative role of viruses in chronic neurological disorders. A recent longitudinal study of 785 UK biobank participants demonstrated that SARS-CoV-2 infection led to a significant reduction in grey matter thickness and global brain volume, in addition to greater cognitive decline in infected individuals compared to non-infected controls (15). These longitudinal results corroborated previous observations about the detrimental effects of SARS-CoV-2 infection on the brain and cognition of infected individuals (4, 34). Furthermore, in older adults (age ≥65 years), COVID-19 patients were shown to have significantly higher risk for a new diagnosis of AD within 360 days after the initial COVID-19 diagnosis, in comparison with age-matched controls (35). Moreover, results from recent studies in animal models and in brain tissue of COVID19-patients suggest that SARS-CoV-2 infection of the brain is associated with amyloid protein aggregation (14, 36). In addition, several SARS-CoV-2 proteins have been shown to have amyloidogenic potential (37, 38). Similar findings have been previously demonstrated for HSV-1, where HSV-1 reactivation was shown to increase AD risk (39), and HSV-1 infection was shown to induce amyloid aggregation in animal models (22, 40, 41), including the induction of amyloid deposition in the 5XFAD mouse model of AD after stereotactic injection of the virus into the hippocampal region (21). Thus, in parallel with the advancements in etiologically connecting viruses to neurodegenerative diseases, it is also important to uncover the pathophysiological mechanisms linking the two.

We have previously shown that viruses can directly induce amyloid aggregation of purified amyloidogenic proteins such as amyloid-β peptide 1-42 (Aβ42) and amylin via HEN by acting as catalytic surfaces for nucleation (22). Such surface-catalyzed nucleation has also been previously demonstrated with a variety of nanoparticles and different proteins, which illustrates the generality of this mechanism (20, 42–44). In a recent review, we outlined the physicochemical and thermodynamic basis of this interaction in light of the classical nucleation theory, where the presence of viral or nanoparticle surfaces lowers the energy barrier for the phase transition of proteins from the soluble form into the solid amyloid form (23). This phase transition is spontaneous, and any protein sequence possesses the information necessary to adopt the amyloid conformation (cross-β), with no requirement for a protein seed/prion to act as a conformational template. In the current study, we show that HSV-1 and UV-inactivated SARS-CoV-2 catalyze amyloid aggregation of a multitude of proteins in their native environment and at their physiologic concentration in human CSF. This result further supports the notion that protein aggregation is a function of the recipient environment, where a subset of proteins are more susceptible to aggregation triggers, most probably due their high concentration (supersaturation) and their ability to interact with nucleating surfaces (45–47). The molecular proximity at higher concentrations can make the generic *inter*molecular interactions necessary for amyloid formation more favorable than the specific *intra*molecular interactions required for native folding, while surfaces act as nucleation sites that catalyze phase transition into amyloids (23). In this physical sense, UV-inactivation does not prevent SARS-CoV-2 viral particles from catalyzing amyloid nucleation on their surfaces. Indeed, in our current study, both live HSV-1 and UV-inactivated SARS-CoV-2 induced amyloid aggregation of a highly overlapping set of proteins. Thus, the resistance of infectious amyloidogenic agents to UV inactivation, which was demonstrated early on (16, 17) and was taken as evidence of the non-viral nature of such agents (18, 19), should be reconsidered. UV-inactivation, while it deactivates viral nucleic acids, does not affect the ability of viruses or other membranous structures to act as catalytic surfaces that induce amyloid aggregation. Importantly, the ability of viruses to invade the CNS and replicate, together with their ability to catalyze amyloid nucleation via HEN, make them more likely causes of amyloid aggregation in the brain compared to seed/prion transmission.

Furthermore, we applied mass spectrometry-based proteomics to identify the proteins that were enriched in the amyloid fraction triggered by the two viruses compared to the normal CSF proteome. We found several proteins that were induced to form amyloids by HSV-1 and SARS-CoV-2, many of which are known to be involved in neurodegenerative diseases and have been reported before to be present in amyloid plaques harvested from patients (Supplementary Tables 1 and 2). For example, APLP1, which is highly homologous to APP (amyloid precursor protein), is present in the plaques in the subiculum and entorhinal cortex in AD (48, 49). ApoE is very commonly found in amyloid plaques, not only in AD, but also in kuru and Creutzfeldt-Jakob disease (50, 51). Furthermore, ApoE is one of the major genetic risk factors for AD and has been shown to interact with Aβ in a variety of ways (52). The same has been demonstrated for clusterin and α2-macroglobulin, which were shown to be associated with amyloid plaques in AD (53, 54). We also identified other proteins that are often related to PD. For example, PGK1, whose deficiency is associated with young-onset parkinsonism (55), and ceruloplasmin and nucleolin, which have been shown to be lower in patients with PD compared to controls (32, 56). Interestingly, 14-3-3 proteins, were also identified in the viral induced amyloid fractions. The 14-3-3 protein family is highly abundant in the brain and has been found to accumulate within Lewy bodies in PD and in plaques and tangles in AD (57–61). They have also been shown to interact with SARS-CoV-2 (62). Moreover, we identified proteins commonly associated with systemic amyloidosis such as transthyretin (TTR) and vitronectin (33, 63). However, both have also been detected in CNS amyloids (64, 65). Other proteins identified in the viral-induced amyloid fractions include immunological components (e.g. IGHG1, IGHG2, complement C3), ribosomal proteins (e.g. ribosomal protein L27a, ribosomal protein L3, ribosomal protein L34), extracellular matrix proteins (e.g. collagen type XVIII alpha 1 chain, keratin 9, fibronectin 1) cytoskeleton proteins (e.g. actin-related protein 2, microtubule-associated protein RP/EB family member 3), heatshock proteins (e.g. HSP 60, 70 and 90) and proteasome proteins (e.g. proteasome 26S subunit alpha 1, proteasome activator subunit 1). While many of these proteins have been detected before in patient-derived plaques (31), further studies are required to address their contribution to a particular disease.

To demonstrate the effect of virus-induced amyloid aggregation on the soluble protein fraction of the CSF, we removed the amyloid fraction that was formed after viral incubation and then ran the soluble fraction on a gel in comparison to CSF treated with non-infected cell medium. We found that the viral-treated CSF was depleted of a large proportion of its soluble proteins compared to the control CSF, indicating that a large fraction of the soluble CSF proteome ended up in amyloids (Figure 2B). Our results suggest that catalyzing protein aggregation and subsequent depletion and loss-of-function might be one mechanism by which viruses cause neurodegeneration. We have recently demonstrated that protein depletion and related loss-of-function is more pathologically important than plaque burden in AD, where high levels of soluble Aβ42 in the CSF were associated with preserved cognition even in individuals with high amyloid plaque burden (24). Depletion of soluble Aβ42 in the CSF, which is a recognized feature of many neurodegenerative disorders, has been demonstrated in patients with post-COVID neurological symptoms (66) and in post infectious neurological sequelae involving other viruses including HSV, varicella zoster virus (VZV) (67), enteroviruses (68) and HIV (69). This suggests that protein depletion post-infection might contribute to the neurological symptoms. Further studies examining the levels of CSF proteins in patients suffering neurological symptoms post infection could reveal additional information on this potential pathophysiological mechanism.

In conclusion, we have demonstrated that HSV-1 and SARS-CoV-2 induce aggregation of a multitude of proteins in human CSF. We have also demonstrated that UV-inactivation does not destroy the ability of viral particles to act as catalytic surfaces for amyloid nucleation. Hence, the role of viruses as causative agents of protein aggregation in neurodegeneration needs to be reevaluated in light of the important role they can play in this process without the need for a proteinaceous conformational template. Understanding the mechanisms by which viruses may cause neurological disorders is especially important due to the SARS-CoV-2 pandemic which has led to a large number of people suffering from long-COVID neurological symptoms post infection. The availability of large and accurate databases of patients together with more sensitive methods to detect viruses in the brain are making the link between virus infection and neurodegenerative brain disorders more robust (70). Taken together with other mechanisms that might lead to viral-induced brain damage (inflammation, autoimmunity, blood-clots) (71), viral-induced protein aggregation via HEN can contribute to neuronal pathology by catalyzing the aggregation and depletion of important neuronal proteins.

## Materials and Methods

### Viral production

For HSV-1, strain F HSV-1 stocks were prepared by infecting African green monkey kidney (vero) cells at 80– 90% confluency in serum-free VP-SFM (Gibco). The virus was harvested 2 d after infection. Cells were subjected to two freeze-thaw cycles and spun at 20,000 g for 10 min to remove cell debris. Clarified supernatant was aliquoted and stored at –80°C until use. Non-infected cell medium was prepared with the same procedure without viral infection. Plaque assay was used to determine viral titers. 10-fold dilutions of virus were added onto vero cells for 1 h at 37°C, then inoculum was removed, and fresh medium containing 0.5% carboxymethyl cellulose (Sigma Aldrich) was added. Cells were fixed and stained 2d later with a 0.1% crystal violet solution and the number of plaques was counted. For SARS-CoV-2 stock production, Vero E6 cells were infected with the SARS-CoV-2 wild type (isolate SARS-CoV-2/human/SWE/01/2020; Genbank accession: MT093571) in VP-SFM (Gibco). The virus was harvested at day three, four and five post-infection and centrifuged at 1000 g for 6 min to remove cells debris. The clarified supernatant was further centrifuged at 45000 g for 4h. After centrifugation, the supernatant was removed, and the virus pellet was resuspended in fresh VP-SFM. Viral titers were determined via end-point-dilution assay. For UV-inactivation of SARS-CoV-2, the virus stock was incubated under UV-light for 3 × 1.5 min, and complete inactivation was confirmed by treating Vero cells with the inactivated stock and observing no cytopathic effects (CPE) 5 days post-treatment. For UV-inactivation of HSV-1, the virus stock was incubated under UV-light for 10 min, and complete inactivation was confirmed by treating Vero cells with the inactivated stock and observing no cytopathic effects (CPE) 3 days post-treatment.

### CSF ethical permit and harvesting

CSF samples were collected at Karolinska University Hospital Huddinge. The collection and test were approved by the Regional Ethical Review Board in Stockholm (Diary number: 2020-03471 and 2009/2107-31-2). All CSF samples were pre-cleared by 400□× □g for 10□min and subsequent 2,000□× □g centrifugation for 10□min. After that aliquoted to a minimal volume of 500 µl and then frozen to –80°C, within 2 hours from sampling.

### CSF amyloid aggregation

ThT (Sigma Aldrich) was prepared at 4 mM in MQ water. For the assay with HSV-1, 50µl of CSF were incubated with 150µl of 4 mM ThT solution and 100 µl of HSV-1 (4.2 × 10^8^ PFU/ml), or non-infected supernatant. For the assay with UV-inactivated SARS-CoV-2, 25µl of CSF were incubated with 150µl of 4 mM ThT solution and 25µl of UV-inactivated SARS-CoV-2 (4.6 × 10^7^ PFU/ml), or non-infected supernatant. UV-inactivated SARS-CoV-2 was used instead of live SARS-CoV-2 for biosafety reasons since live SARS-CoV-2 was not safe to incubate in the spectrophotometer outside the BSL-3 laboratory. Controls included CSF only without virus and virus only without CSF in addition to 150µl of 4 mM ThT solution, and in both cases the missing volume was substituted with MQ water. ThT fluorescence was measured at 440 nm excitation and 480 nm emission in a black clear-bottom 96-well plates (Corning, USA) at 440 nm excitation and 480 nm emission at 10-15 min. intervals (from bottom with periodic shaking) over 24-48 h on SpectraMax i3 microplate reader (Molecular Devices, USA). Curves were fitted using GraphPad Prism software.

### Electron Microscopy

For TEM, equal volumes (50 µl) of viruses (HSV-1 (4.2 × 10^8^ PFU/ml) or UV-inactivated SARS-CoV-2 (4.6 × 10^7^ PFU/ml) and CSF were incubated at 37 °C for 48 h. Samples were then applied to Formvar/carbon coated 200 mesh nickel grids (Agar Scientific, UK), then negatively stained using an aqueous solution of uranyl acetate (1%) and visualized.

### Amyloid purification

Equal volumes (100 µl) of virus (HSV-1 (4.2 × 10^8^ PFU/ml) or UV-inactivated SARS-CoV-2 (4.6 × 10^7^ PFU/ml) and CSF were incubated at 37 °C for 48 h. Amyloid aggregates were then collected by spinning at 20 000 g for 15 min. at room temperature. The pellet was then washed 2x by vortexing in 100 µl lysis buffer (4% SDS, 25 mM HEPES pH 7.6, 1 mM DTT) for 2 min. followed by 15 min, 20 000 g centrifugation at room temperature. The pellet was then dissolved in 99% formic acid by vortexing for 2 min and sonication for 5 minutes. Formic acid was then removed by speedvacing at room temperature for 30 min. and the proteins were resuspended in 50 µl PBS for proteomic analysis.

### Sample preparation for proteomics

The samples collected were lysed by 4 % SDS lysis buffer and prepared for analysis with mass spectrometry using a modified SP3 clean up and digestion protocol. In brief, the sample was alkylated with 4 mM hloroacetamide. 20 µl of Sera□Mag SP3 bead mix were added to the protein sample plus 100% Acetonitrile, reaching a final concentration of 70 %. The mix was incubated at room temperature and under rotation for 18 min. The mix was then placed on a magnetic rack and the supernatant was discarded. This was followed by 2 washes with 70 % ethanol and 1 wash with 100 % acetonitrile. The mixture of beads and proteins was then reconstituted in 100 µl LysC buffer (50 mM HEPES pH 7.6, 0.5 M Urea, and 1:50 enzyme (LysC): protein ratio) and incubated overnight. At the last step, trypsin was added (1:50 enzyme: protein ratio) in 100 µl 50 mM HEPES pH 7.6 and incubated overnight followed by SP3 peptide clean up. The peptide mix was then suspended in 15 μl LC mobile phase A and 5 μl were injected into the LC-MS/MS system. These methods were similar to what was published earlier in (72, 73).

### Mass spectrometry

LC-ESI-MS/MS, Q-Exactive Online LC-MS was performed on a Dionex UltiMate™ 3000 RSLCnano System, which was coupled to a Q-Exactive mass spectrometer (Thermo Scientific). 5 uL were injected from each sample. A C18 guard desalting column was used for samples entapment (Acclaim PepMap 100, 75µm x 2 cm, nanoViper, C18, 5 µm, 100 Å), and the samples were thereafter separated on a 50 cm C18 column (Easy spray PepMap RSLC, C18, 2 µm, 100Å, 75 µmx15cm). The composition of the nano capillary solvent A was: 5%DMSO, 95% water, 0.1% formic acid; and solvent B was 5% DMSO, 5% water, 0.1% formic acid, 95% acetonitrile. The curved gradient went from 6%B up to 43%B in 180 min at a constant flow of 0.25 μl min^-1^, followed by an increase to 100%B in 5 min. FTMS master scans with mass range 300-1500 m/z (and 60,000 resolution) were followed by data-dependent MS/MS (30000 resolution) on the top 5 ions utilizing higher-energy collision dissociation (HCD) where collision energy was normalized at 30%. A a 2m/z window was used for isolation of precursors. The targets for automatic gain control (AGC) were 1e6 for MS1 and 1e5 for MS2. Maximum injection times were 100ms for both MS1 and MS2 and the whole duty cycle lasted approximately 2.5s. Precursors with unassigned charge state or charge state 1 were excluded and dynamic exclusion was used with 60s duration. The underfill ratio was 1%. These methods were similar to what was published earlier in (72, 73).

### Bioinformatics

MsConvert from the ProteoWizard tool suite (74) was used to convert Orbitrap raw MS/MS files to mzML format using and processed with a pipeline built using Nextflow (75) with MSGF+ (76) as a peptide identification search engine, Percolator for target-decoy scoring (77) and Hardklor/Kronik (78) for quantifying MS1 peptide features. For the MSGF+ settings, precursor mass tolerance was 10 ppm, fully-tryptic peptides, maximum length for peptides was 50 amino acids and the maximum charge was 6. The fixed modification was carbamidomethylation on cysteine residues and the variable modification was oxidation on methionine residues. PSMs found at 1% false discovery rate (FDR) were used to infer protein identities. Searches were performed against the human subset of Ensembl 102 and custom-built peptide database. 6 replicates with CSF from two different individuals were analyzed (separate, not pooled). Output from pipeline was filtered, removing protein identifications present in less that 4/6 replicates per sample group any missing values after filtering was imputed using the MissForest method (79). Protein intensity values were then normalized using the cyclic-loess method with the NormalizerDE package (80), differentially expression analysis conducted using LIMMA (81). These methods were similar to what was published earlier in (72, 73).

### PAGE electrophoresis and silver staining

Equal volumes (15 µl) of HSV-1 (2.1 × 10^8^ PFU/ml) or uninfected serum-free cell medium were incubated with CSF (in triplicates with CSF from two different individuals) at 37 °C for 72 h. After incubation, samples were centrifuged at a speed of 20 000 g for 15 minutes, and 15 μl of the supernatant was mixed with NuPAGE LDS Sample Buffer (4x). Then, the samples were Incubated at 80°C for 10 minutes. Samples were then loaded in NuPAGE™ 4-12% Bis-Tris Protein Gels and run for 1.5h at 120V in 1x MES running buffer. After the proteins were electrophoresed in the mini polyacrylamide gel, the gel was silver-stained using the ProteoSilver Silver Stain Kit (Sigma-Aldrich) according to the manufacturer’s instructions. The images of the gels were then taken with a handhold camera with a 2x zooming.

## Supporting information

Reviewer comments & responses 1st round

Reviewer comments & responses 2d round

## Acknowledgements

Support by NBIS (National Bioinformatics Infrastructure Sweden) is gratefully acknowledged.

## Notes

### Competing Interest Statement

Competing interests statement: KE, AS, AJE, SA, and TM are cofounders of REGAIN Therapeutics and co-inventors of some of the company's patents.

### Summary of Updates

This version is updated based on two rounds of review in PNAS. The most significant change is the new Figure 2. where we show that HSV-1 induction of amyloid aggregation in human CSF leads to drastic loss of soluble proteins from the CSF linking viral catalyzed protein aggregation with loss of function mechanism of toxicity. In addition, we attach with this version the two rounds of reviewer comments and our responses to them. Moreover, we added new supplementary figures including all the raw electron microscopy pictures demonstrating viral-induced amyloid fibrils. Furthermore, we corrected some errors found in the earlier version.

