## Supplementary Figures 1 & 2 for "SARS-CoV-2 and HSV-1 Induce Amyloid Aggregation in Human CSF Resulting in Drastic Soluble Protein Depletion"

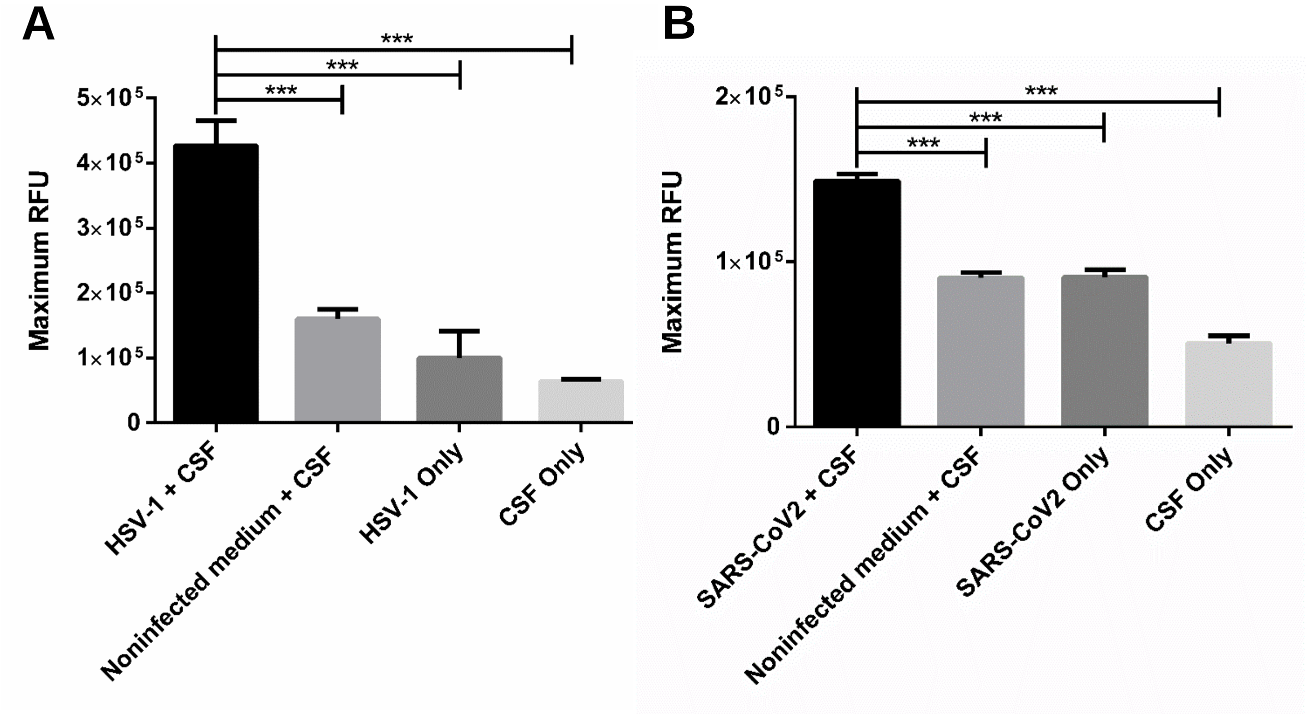


**Supplementary Figure 1. Significant induction of amyloid formation in human CSF upon viral incubation.** Replot of the data from Figure 1. to demonstrate maximum ThT fluorescence enhancement in CSF after incubation with (**A**) HSV-1 or (**B**) UV-inactivated SARS-CoV-2 in comparison to controls. Means ± SEM of 8 replicates with CSF from two different individuals are shown. Significant differences were assessed using one-way ANOVA with Tukey multiple comparisons test. ***$P < 0.001$. RFU = relative fluorescent unit.

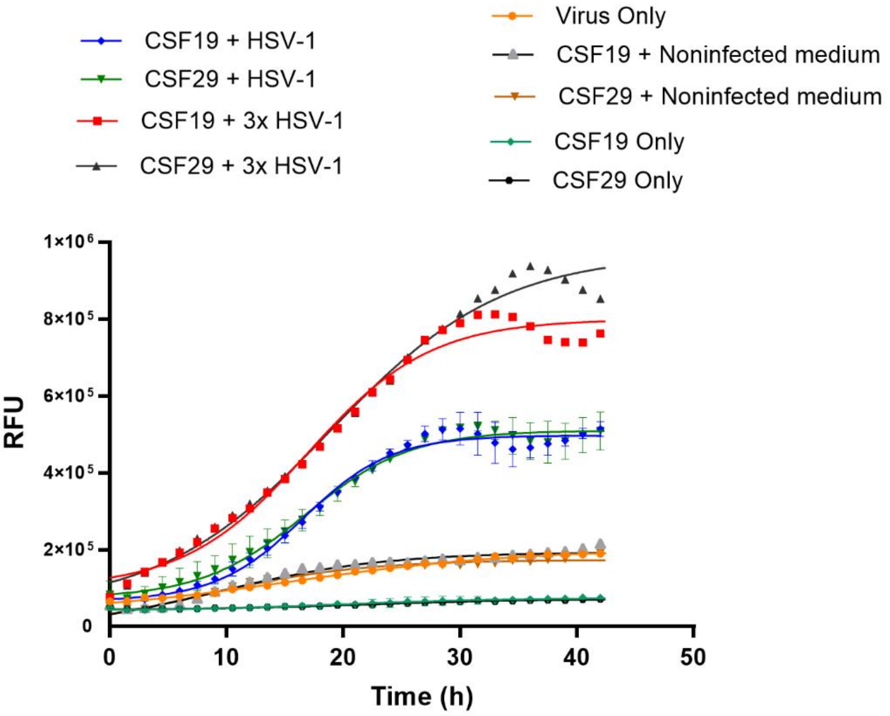


**Supplementary Figure 2. HSV-1 induce amyloid aggregation of proteins in human CSF in a concentration dependent manner.** CSF was incubated with HSV-1 (4.2 x $10^8$ PFU/ml ) or 3x concentrated (1.26 x $10^9$ PFU/ml) and ThT solution. Fluorescence was measured at 440 nm excitation and 480 nm emission over 48 h at 37 °C. Means ± SEM of duplicates with CSF from two different individuals (CSF19 and CSF29) are shown. RFU = relative fluorescent unit.
