## Supplementary Figure 3 for "SARS-CoV-2 and HSV-1 Induce Amyloid Aggregation in Human CSF Resulting in Drastic Soluble Protein Depletion"

**Supplementary Figure 3. Amyloid fibrils formed upon SARS-CoV-2 incubation with human CSF.** Negatively stained TEM images of UV-inactivated-SARS-CoV-2 incubated with CSF for 48 h at 37 °C. Raw pictures.

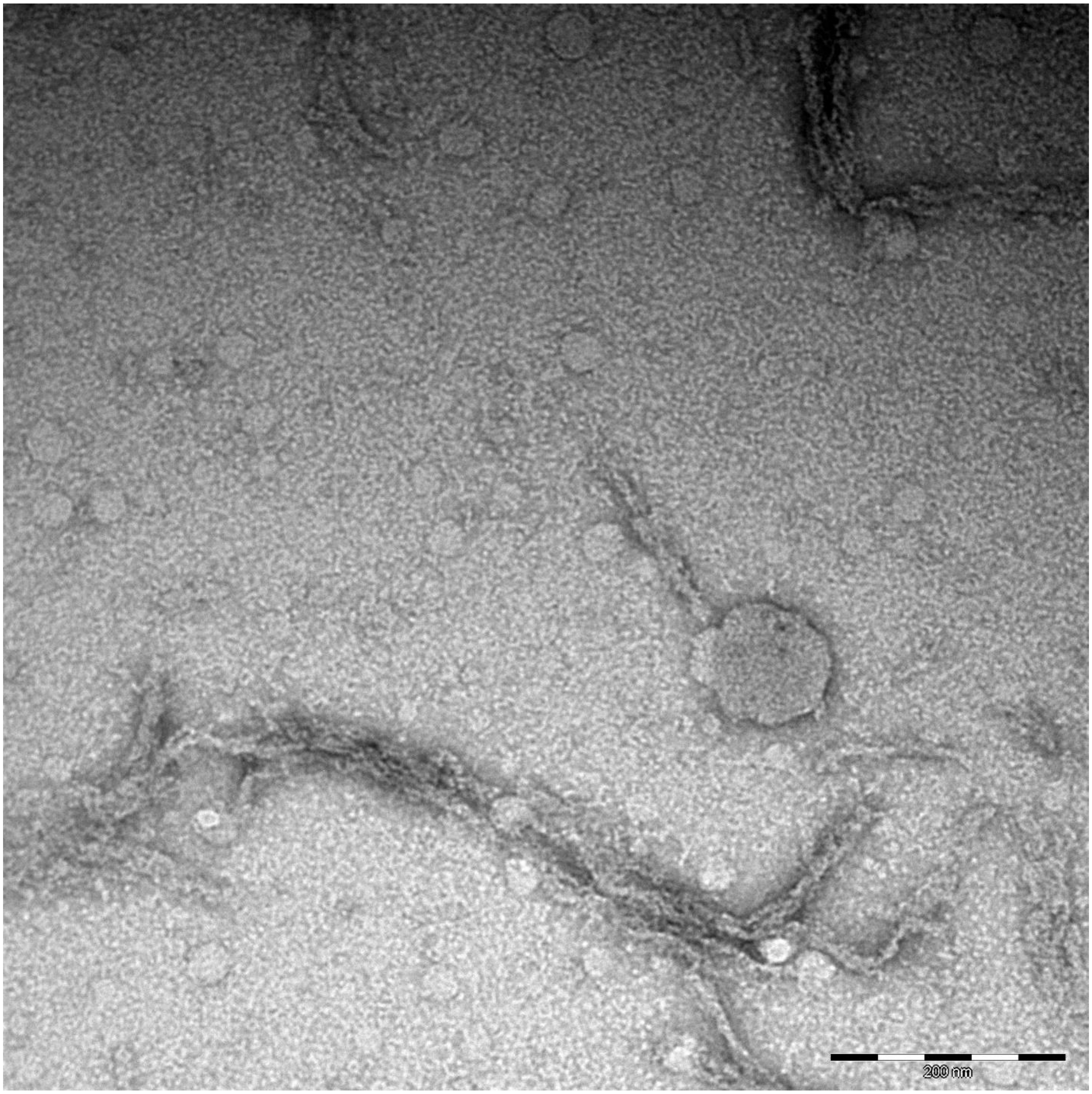
200 nm

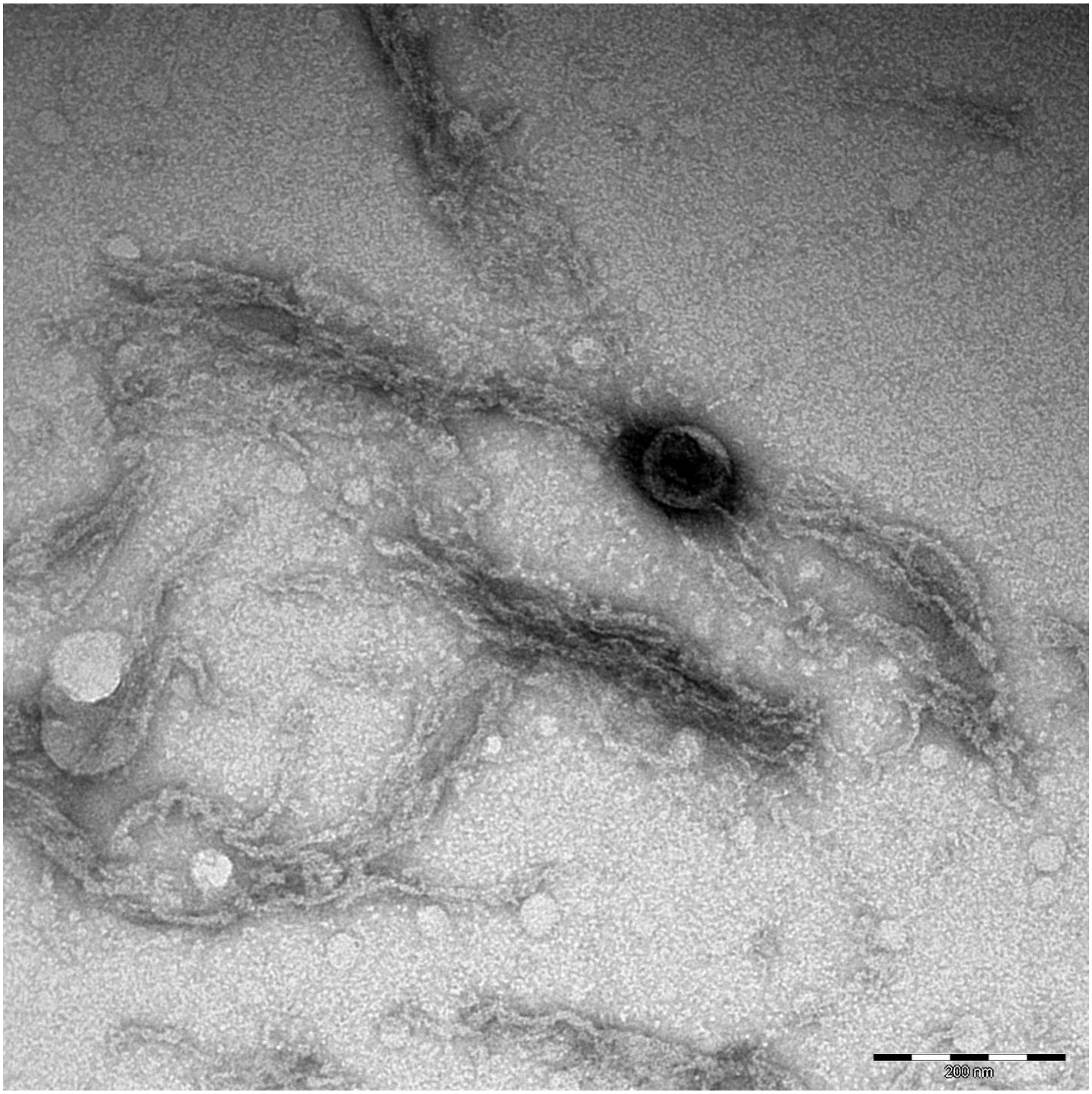
200 nm

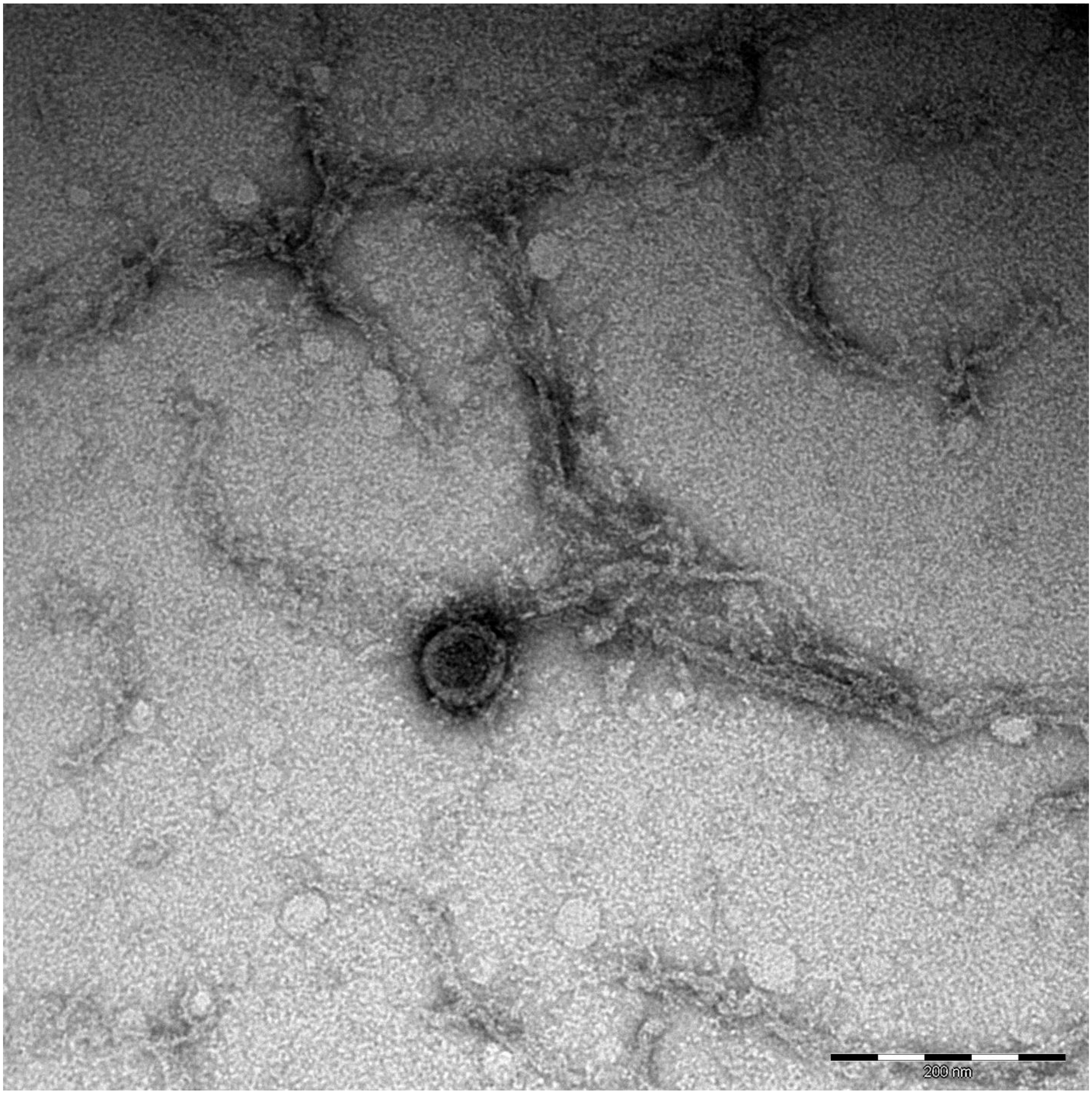
200 nm

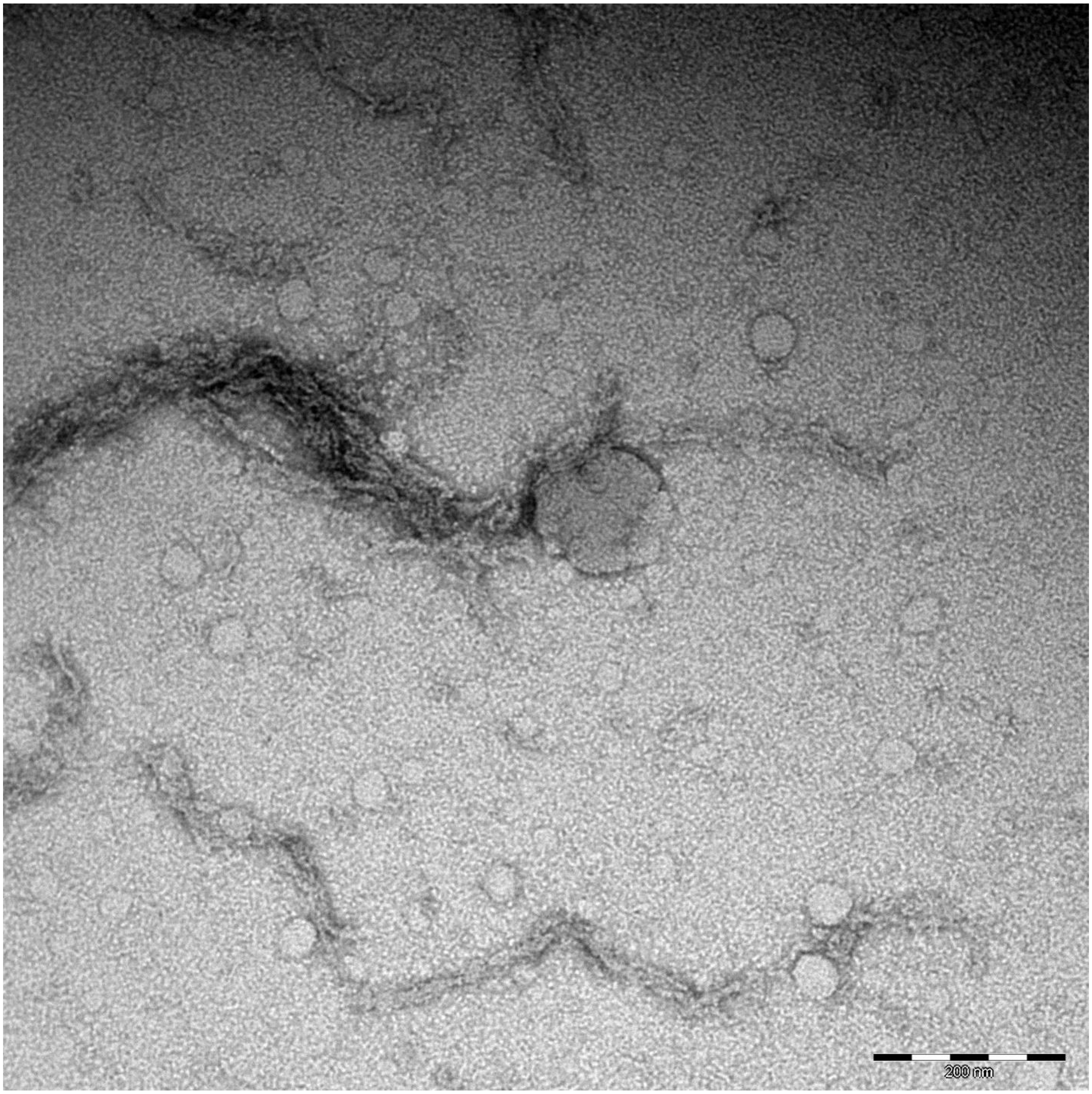
200 nm

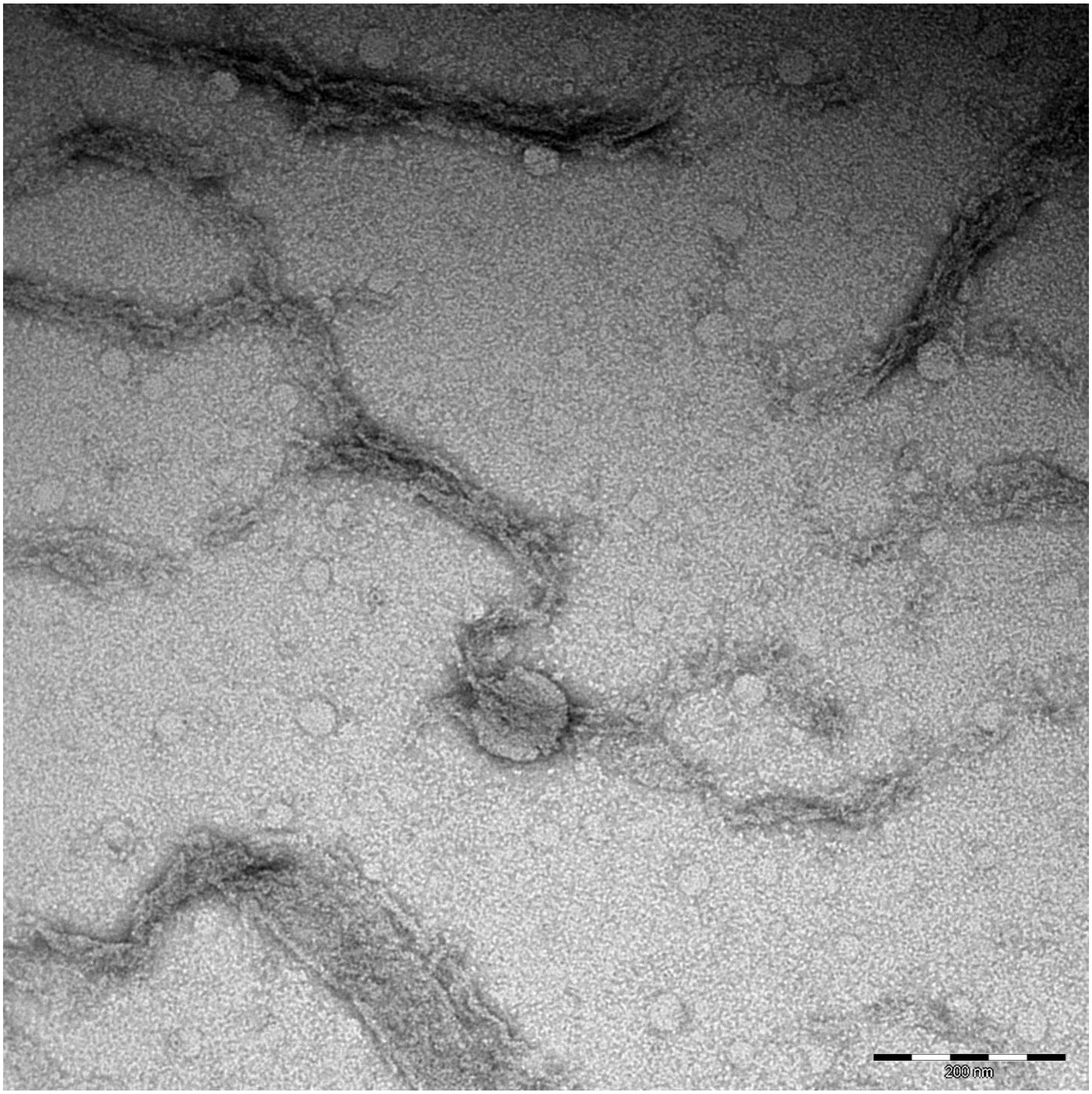
200 nm

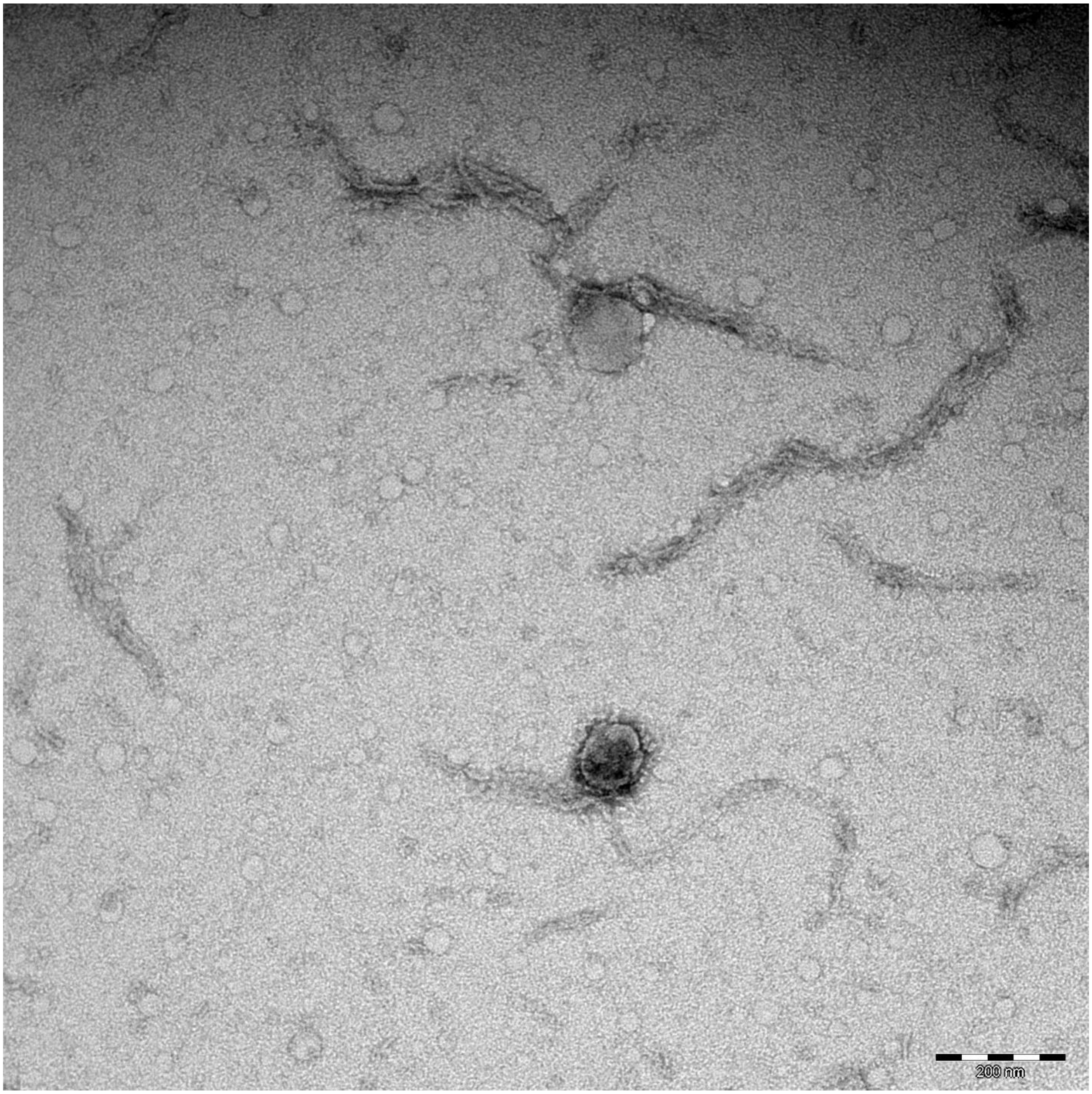
200 nm

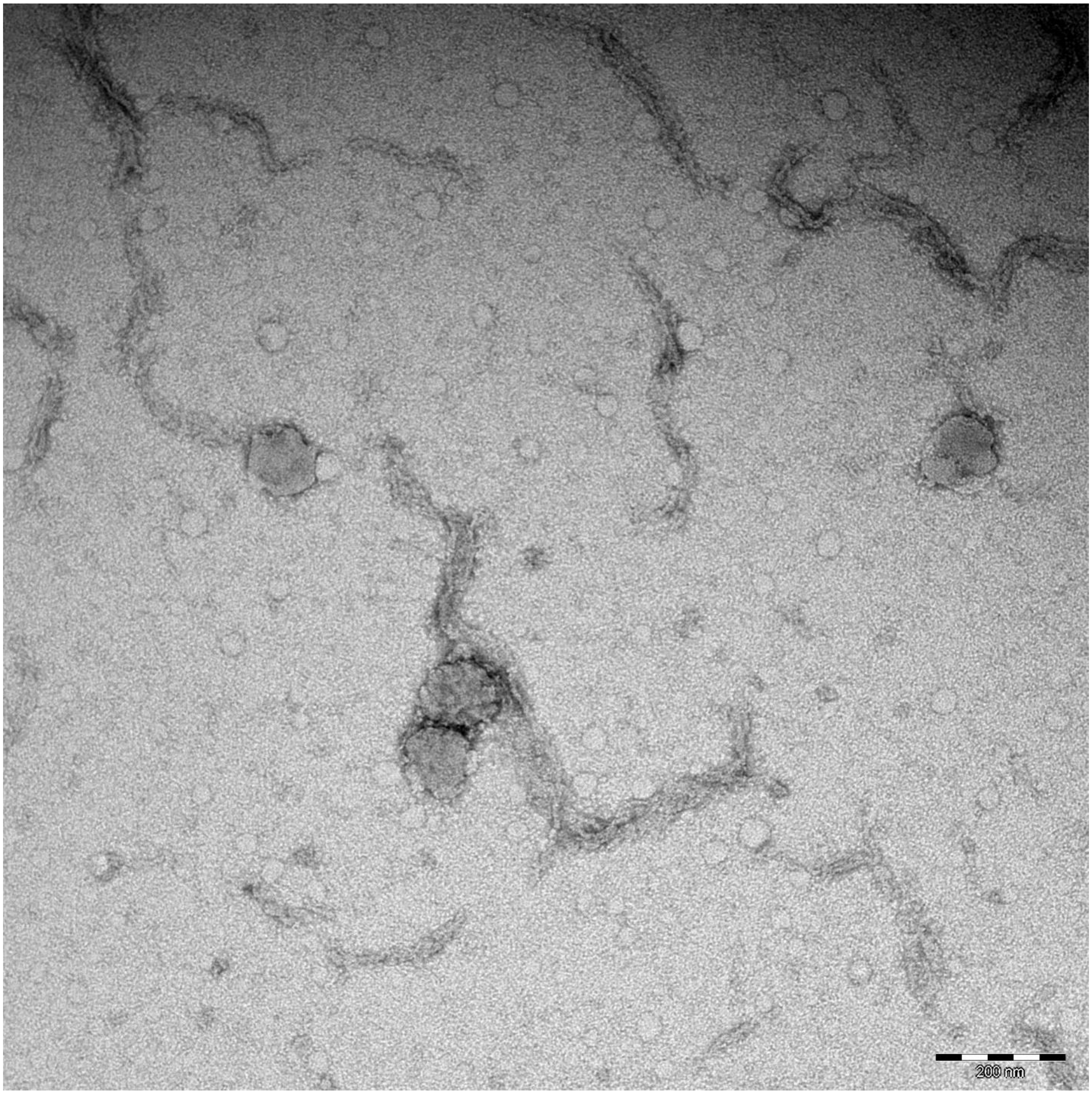
200 nm

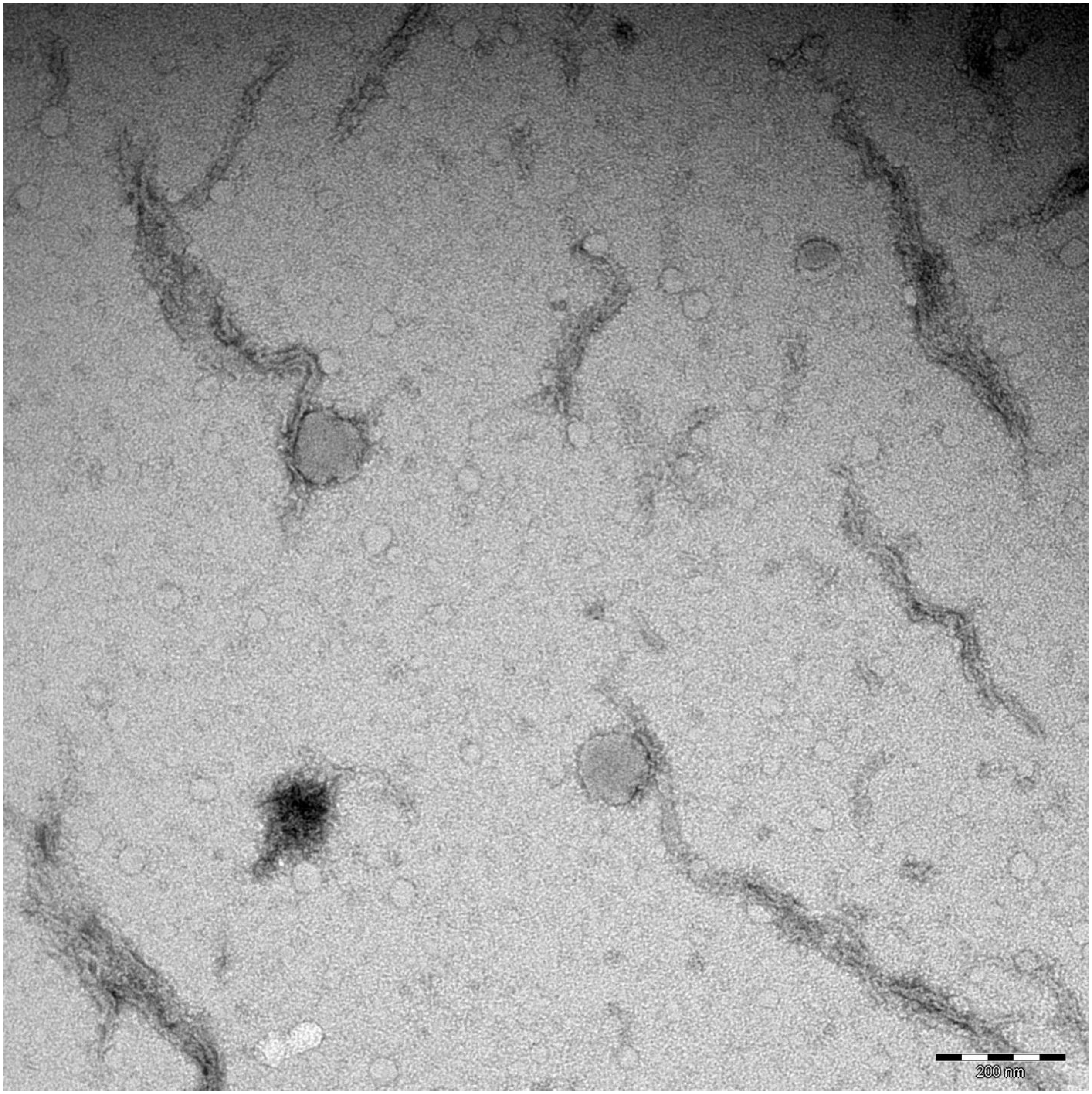
200 nm

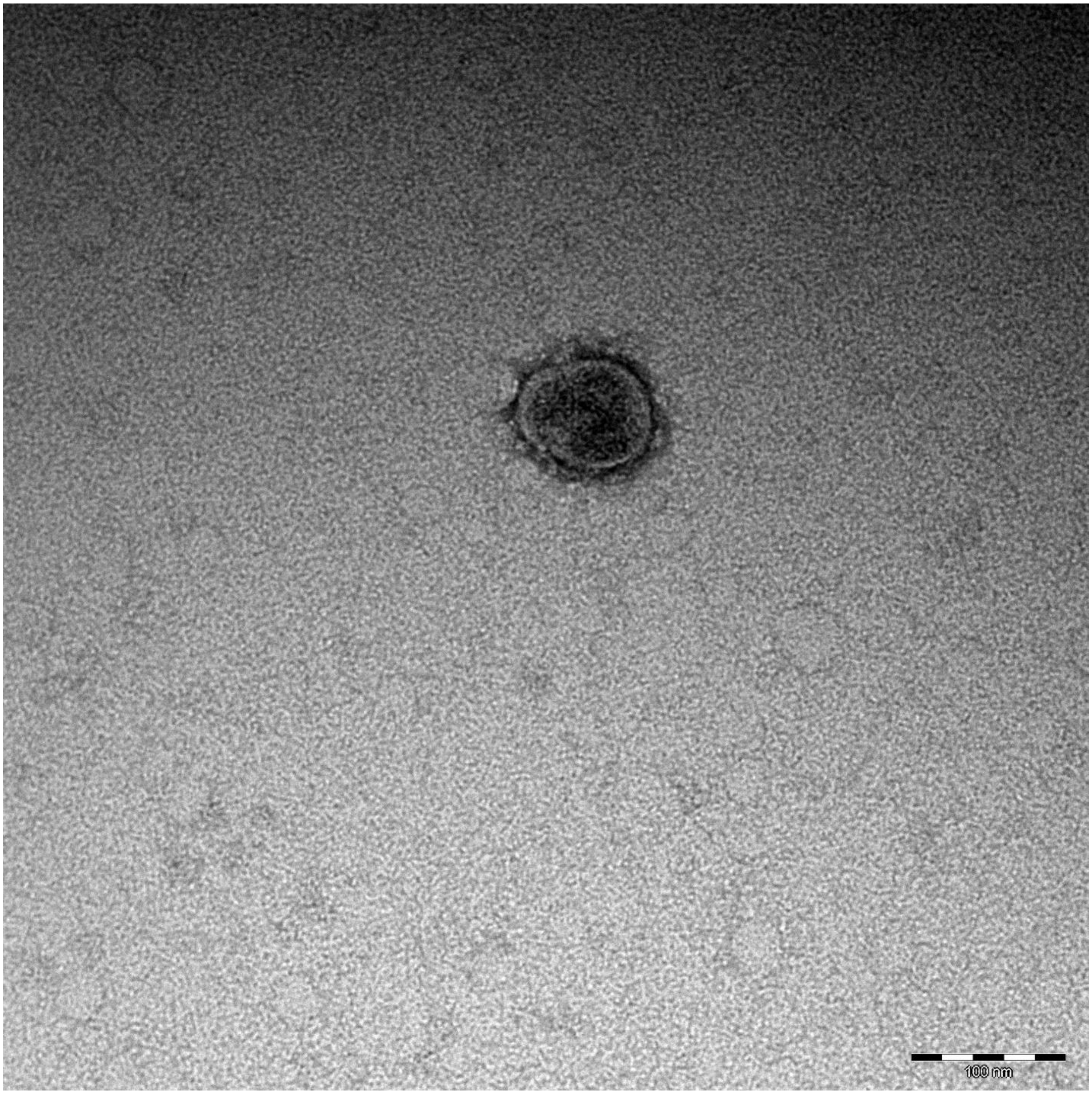
100 nm

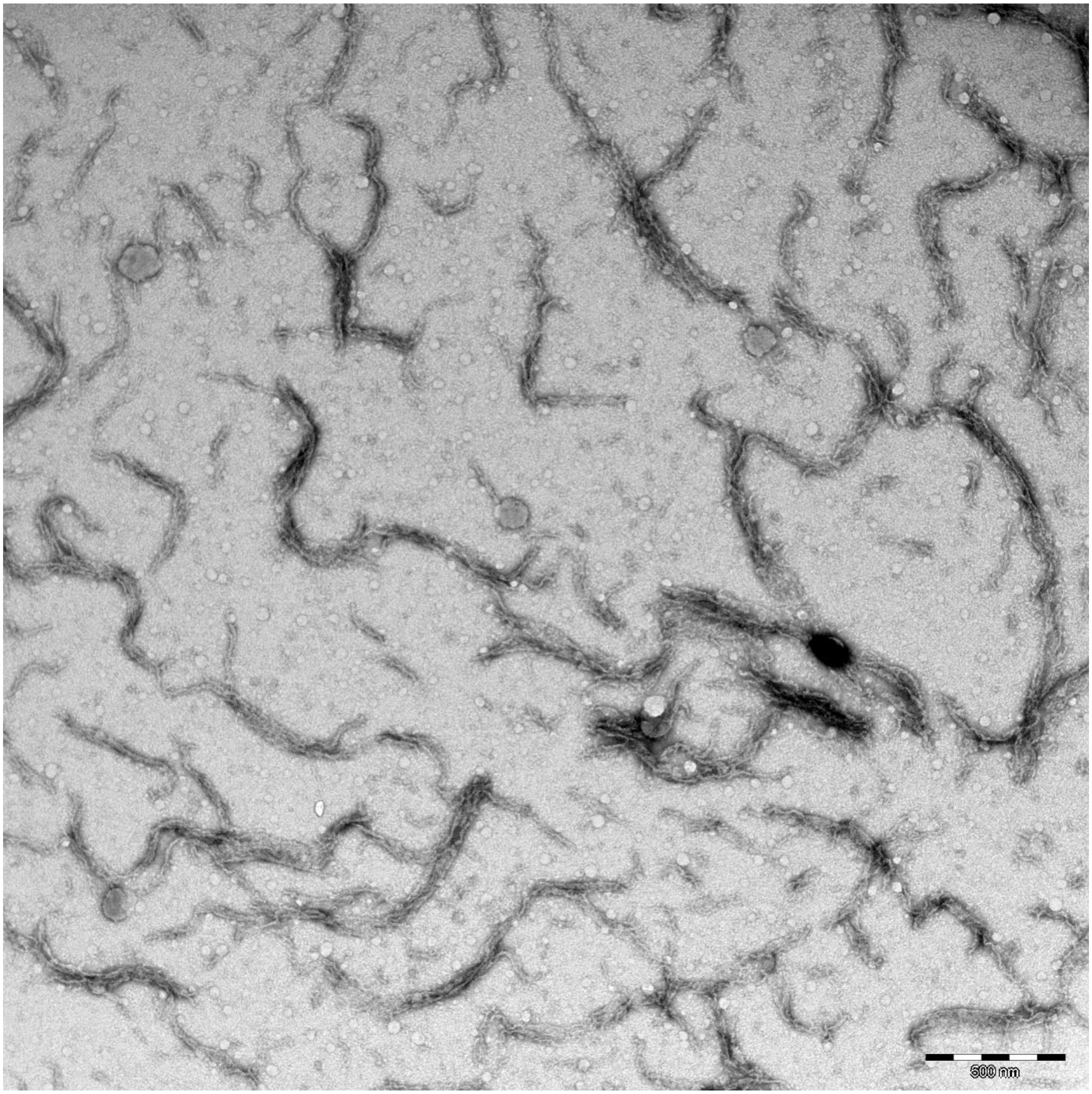
500 nm

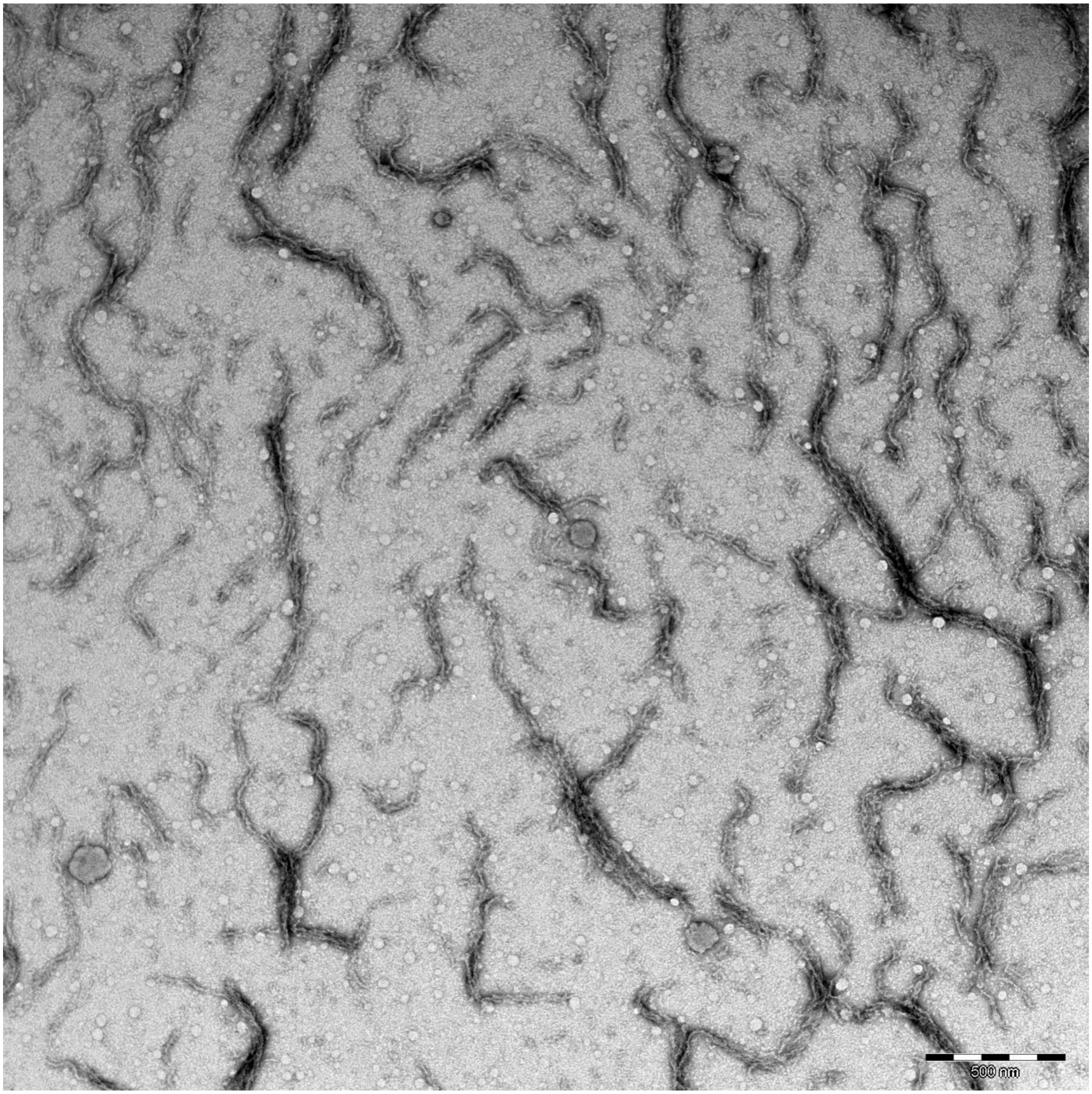
500 nm

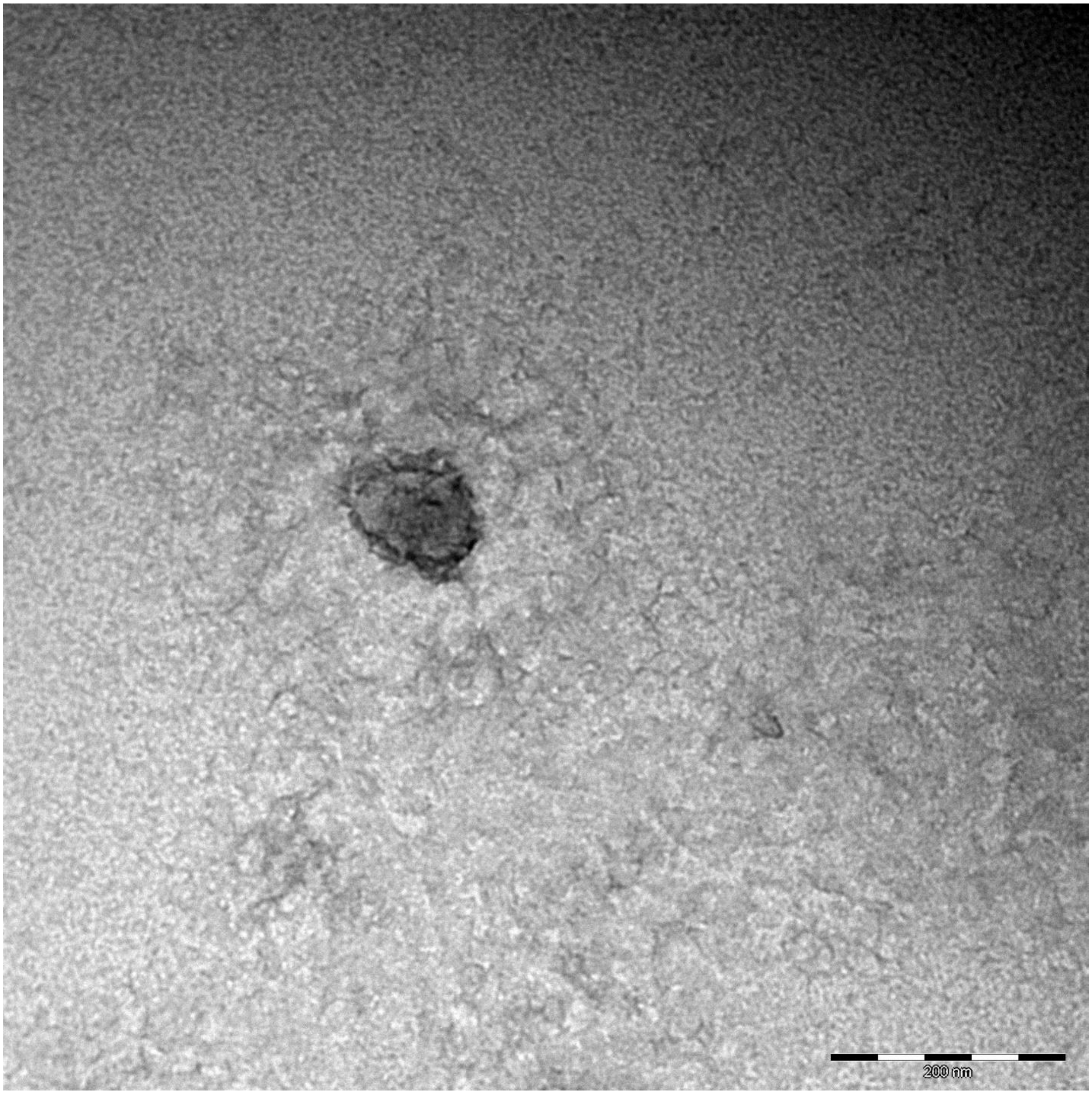
200 nm

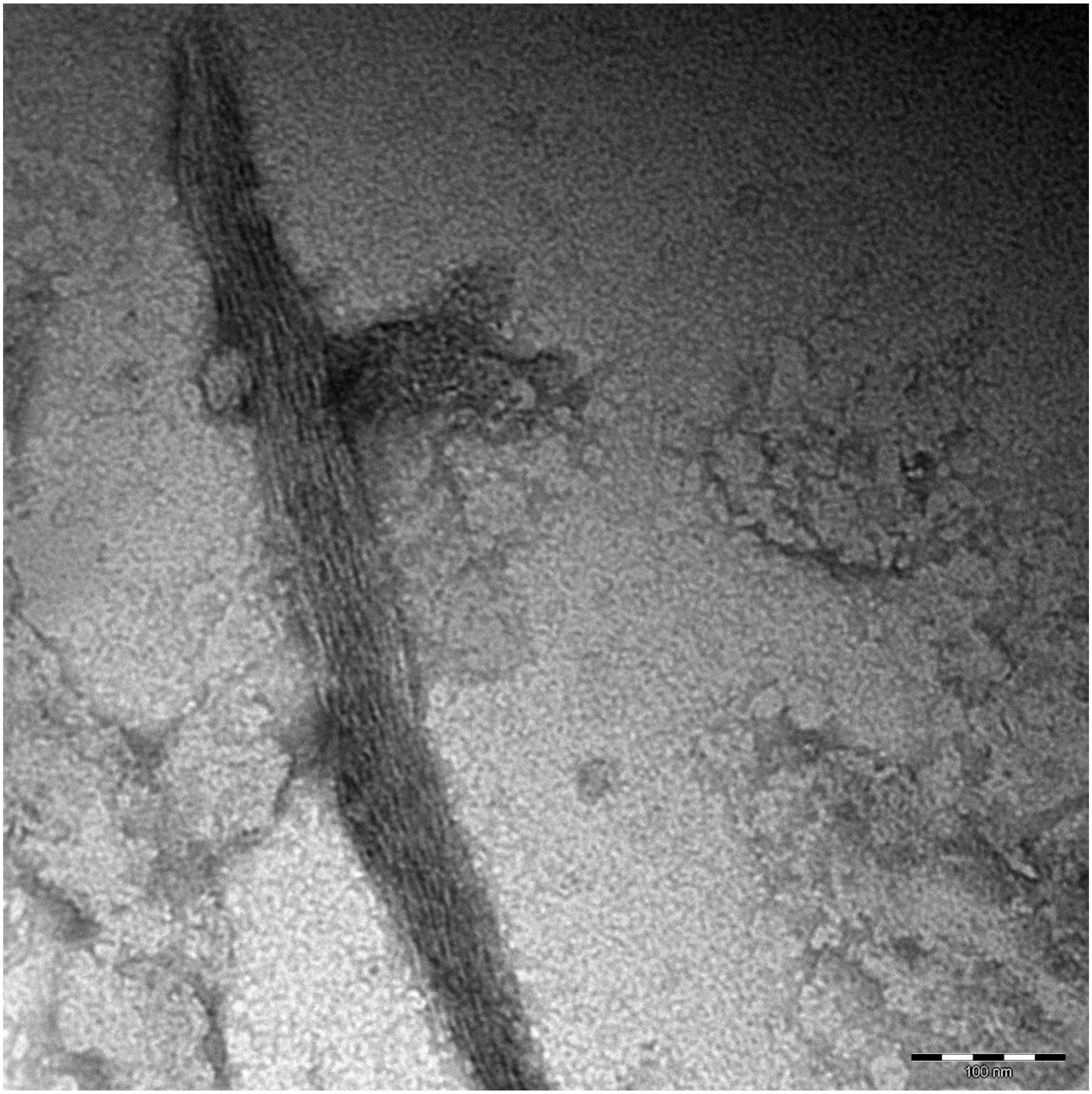
100 nm

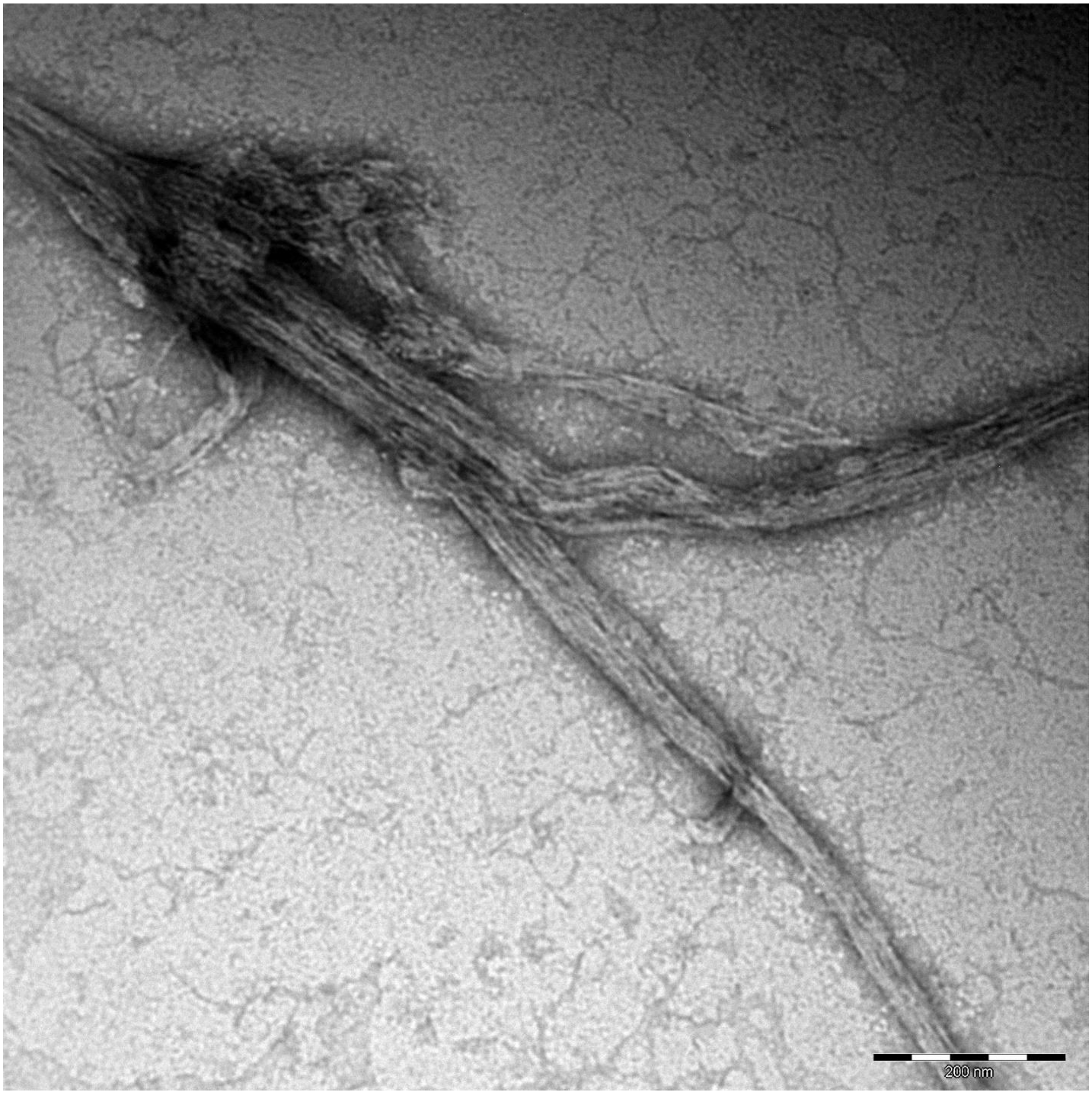
200 nm

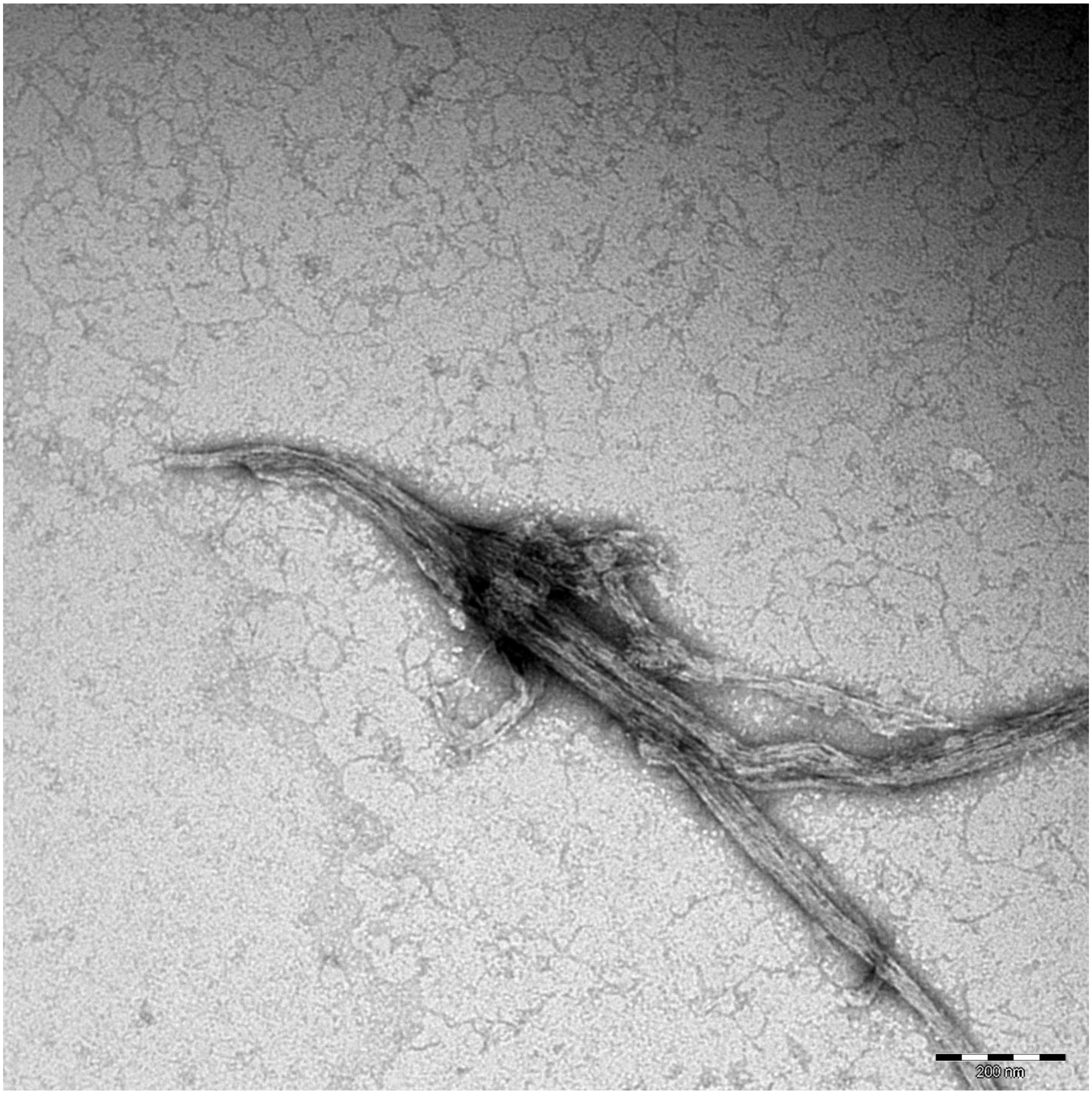
200 nm

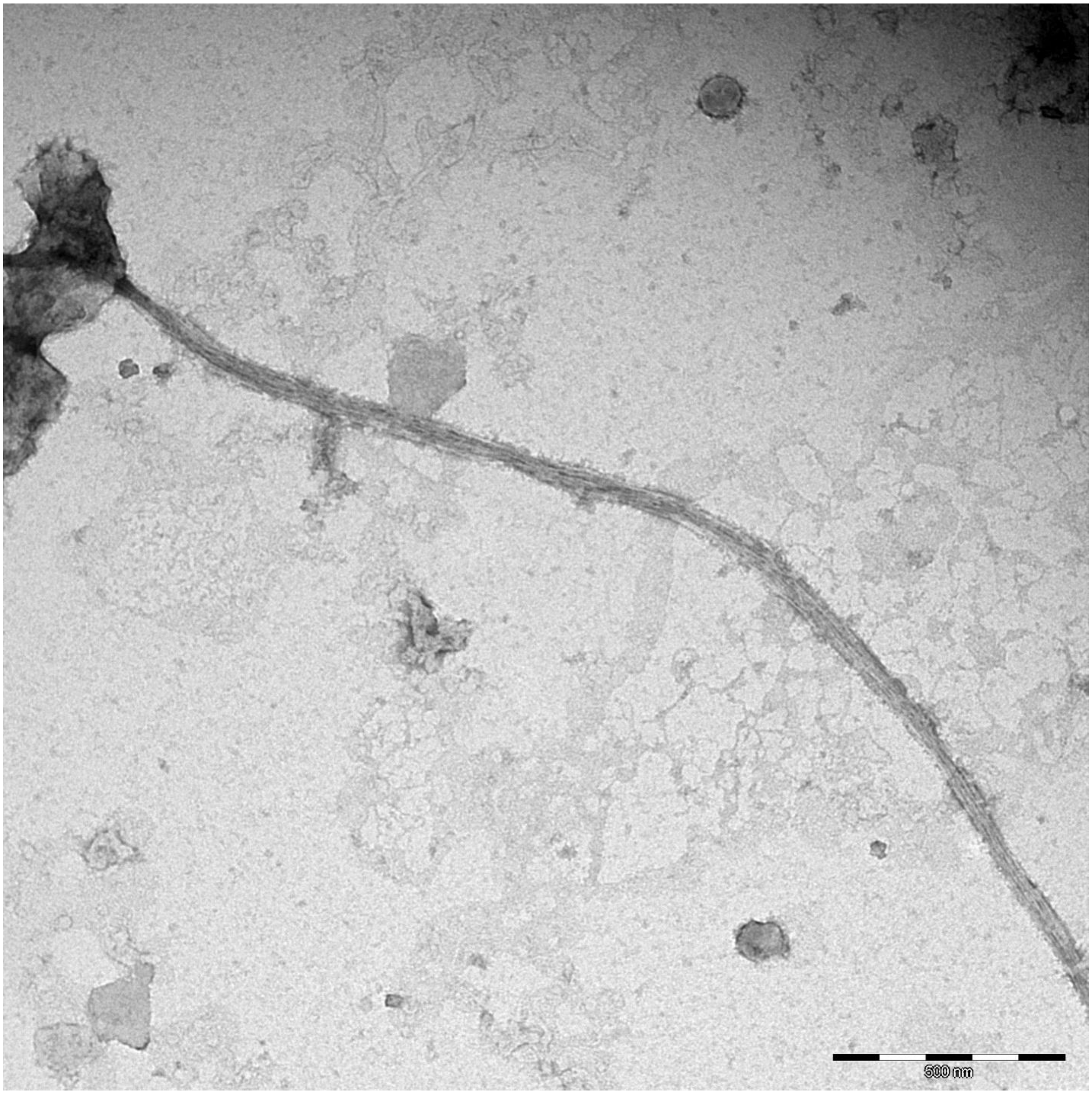
500 nm

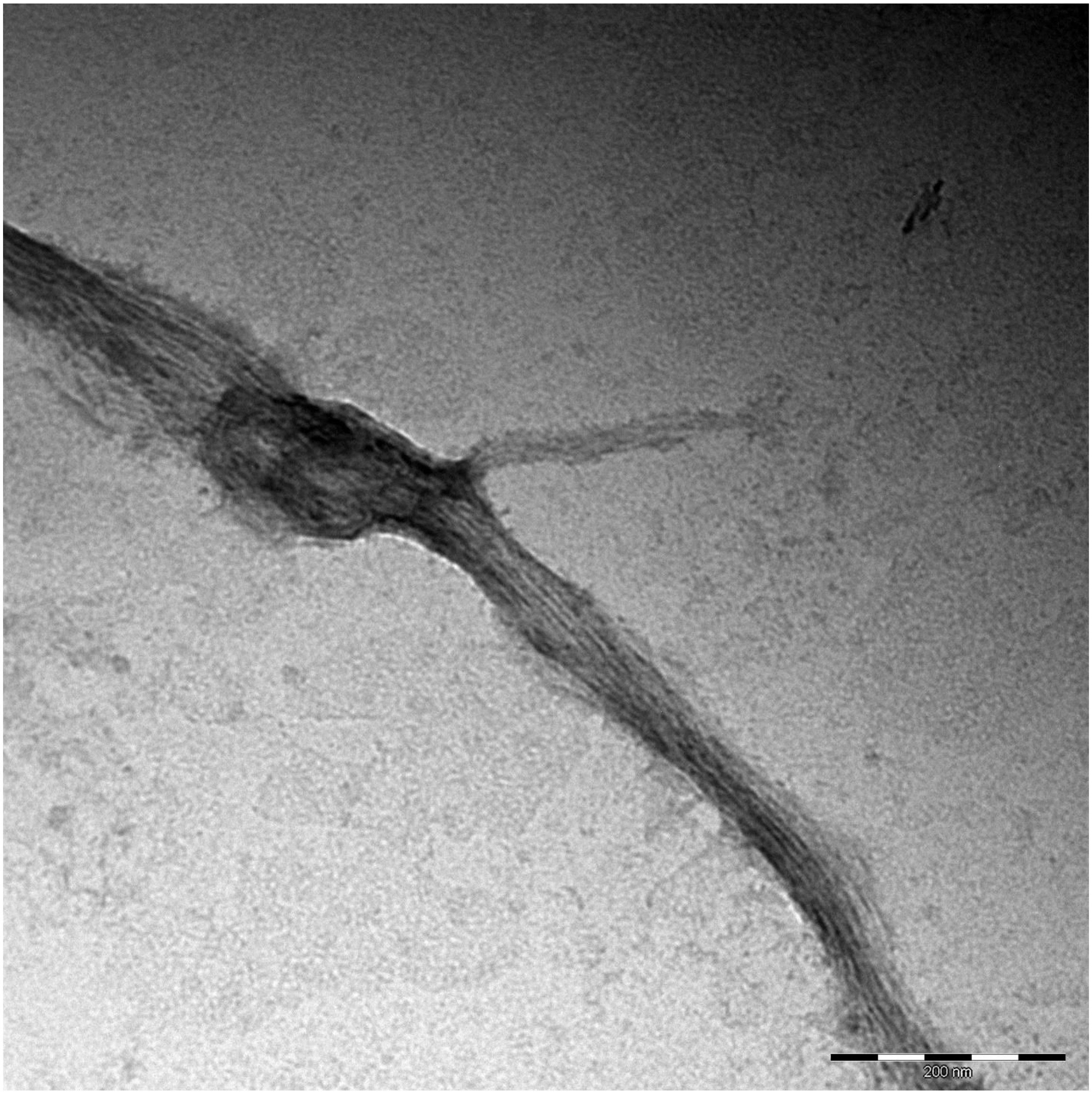
200 nm

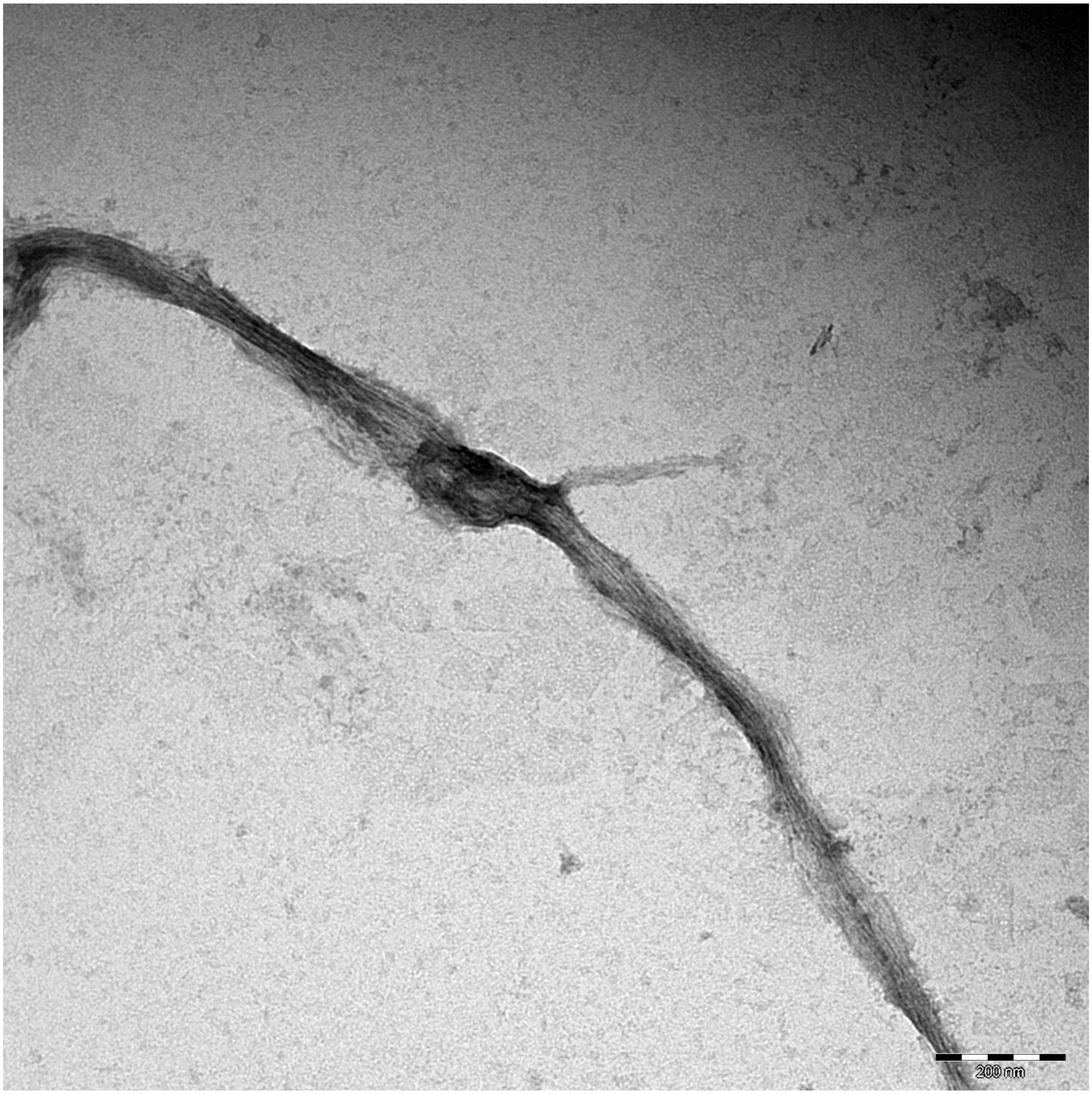
200 nm

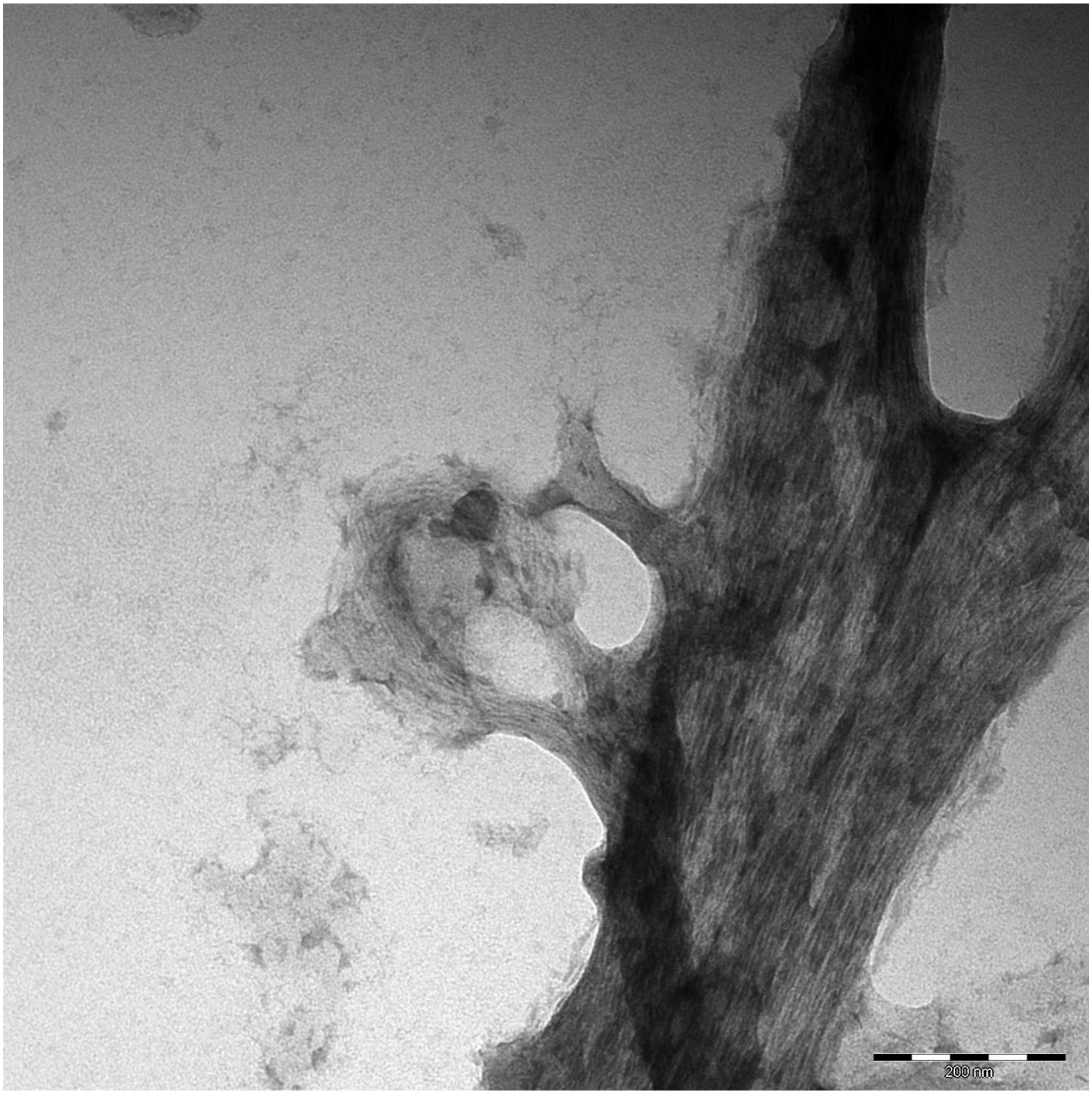
200 nm

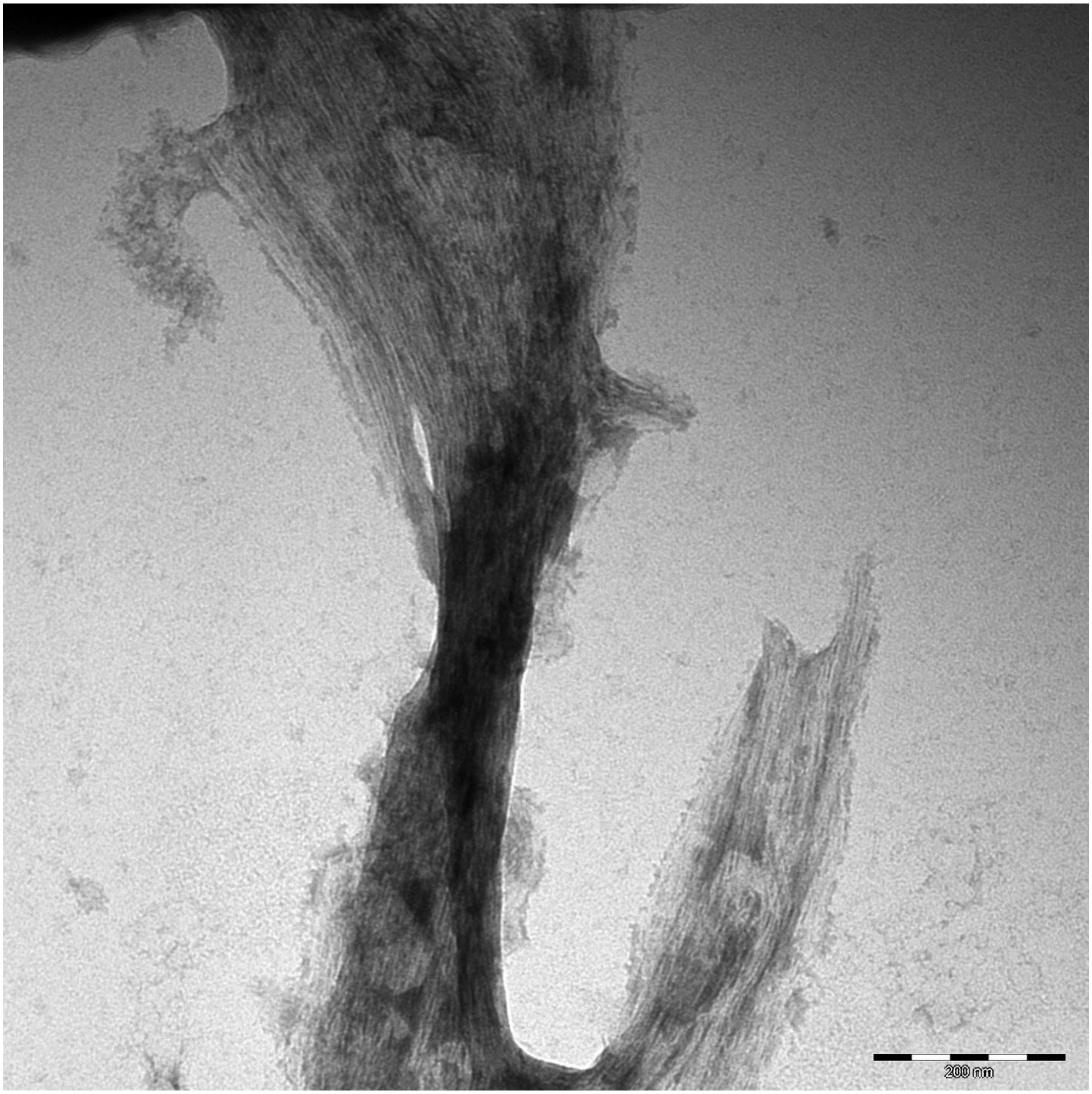
200 nm

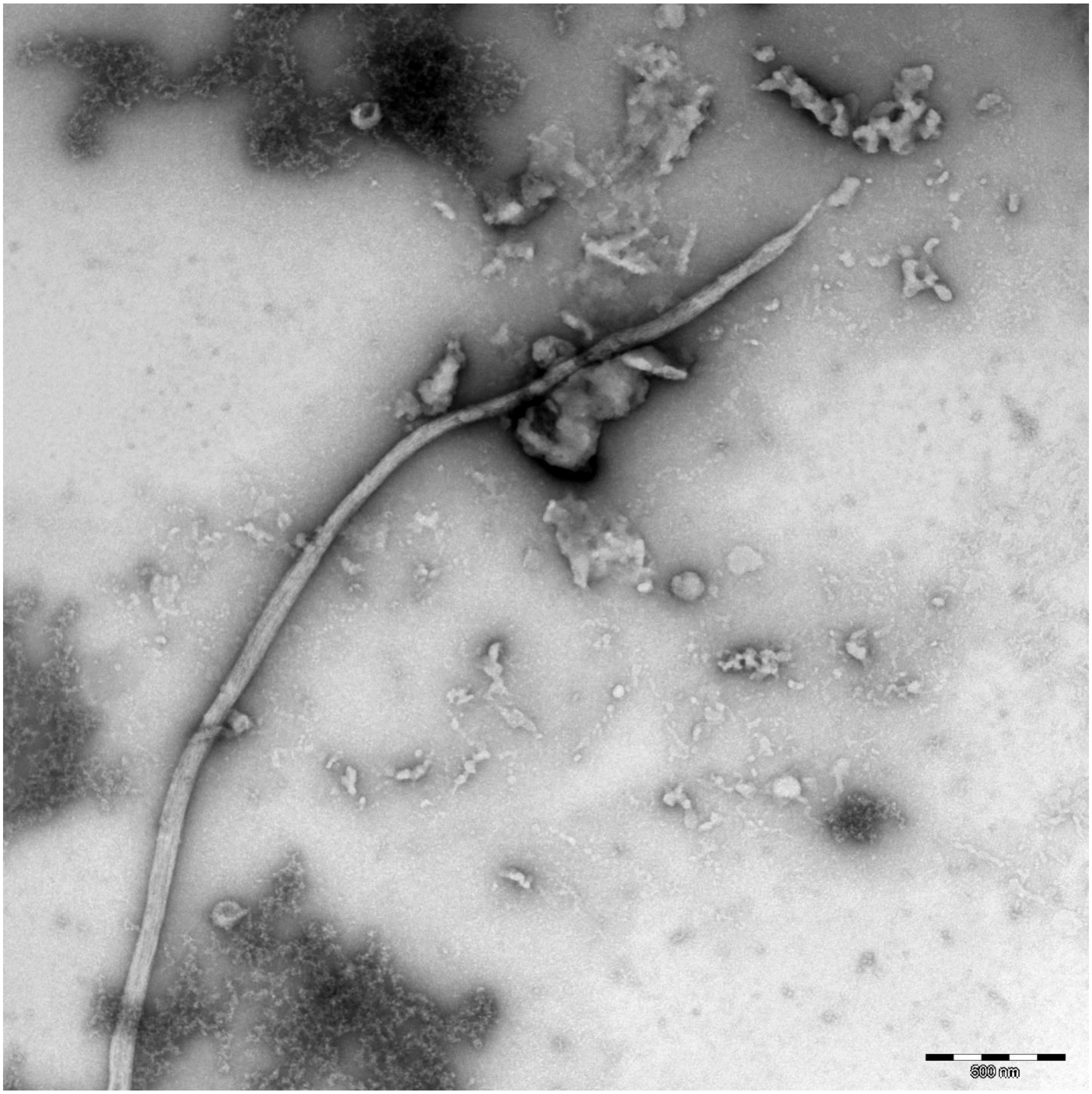
500 nm

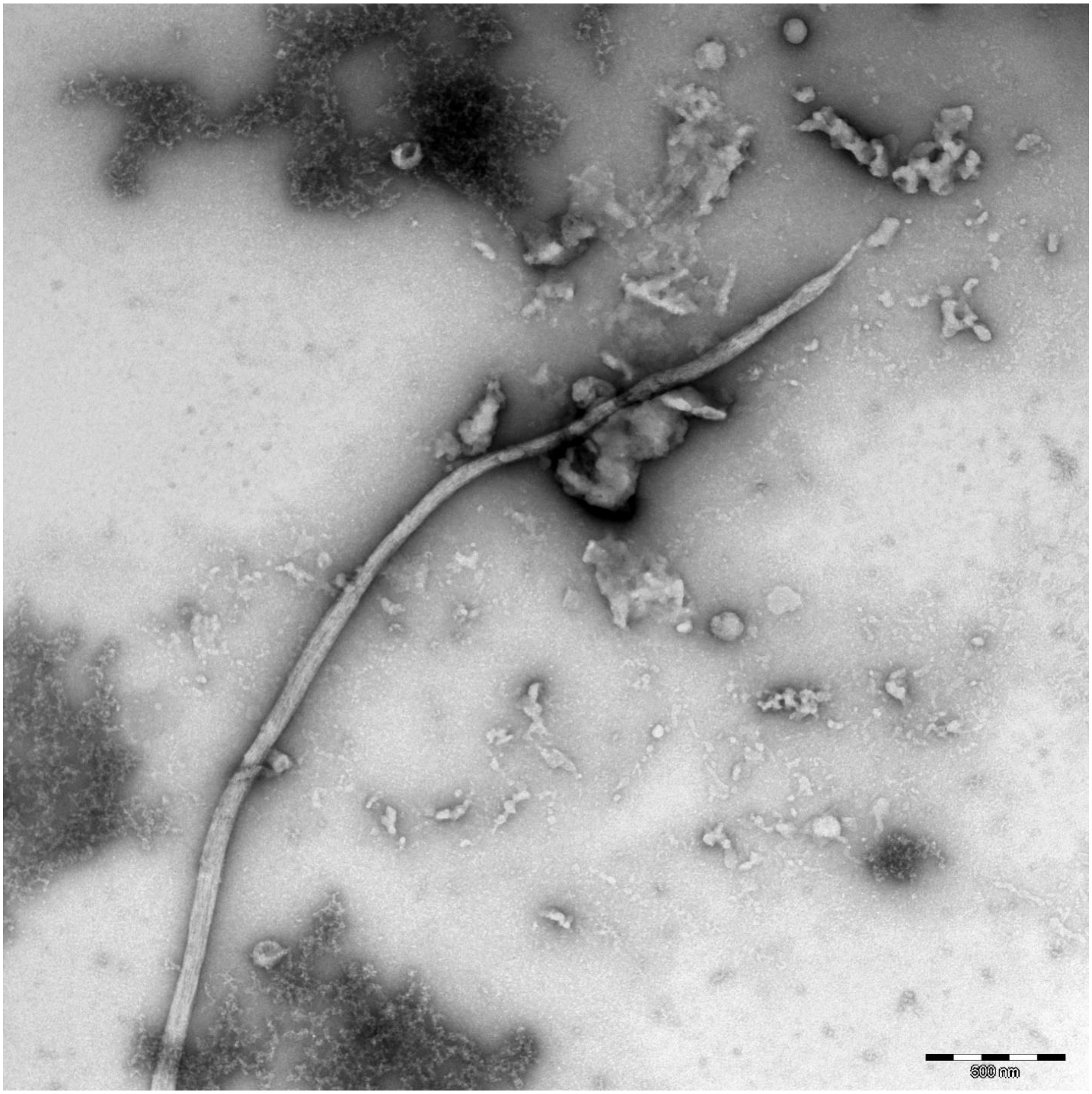
500 nm

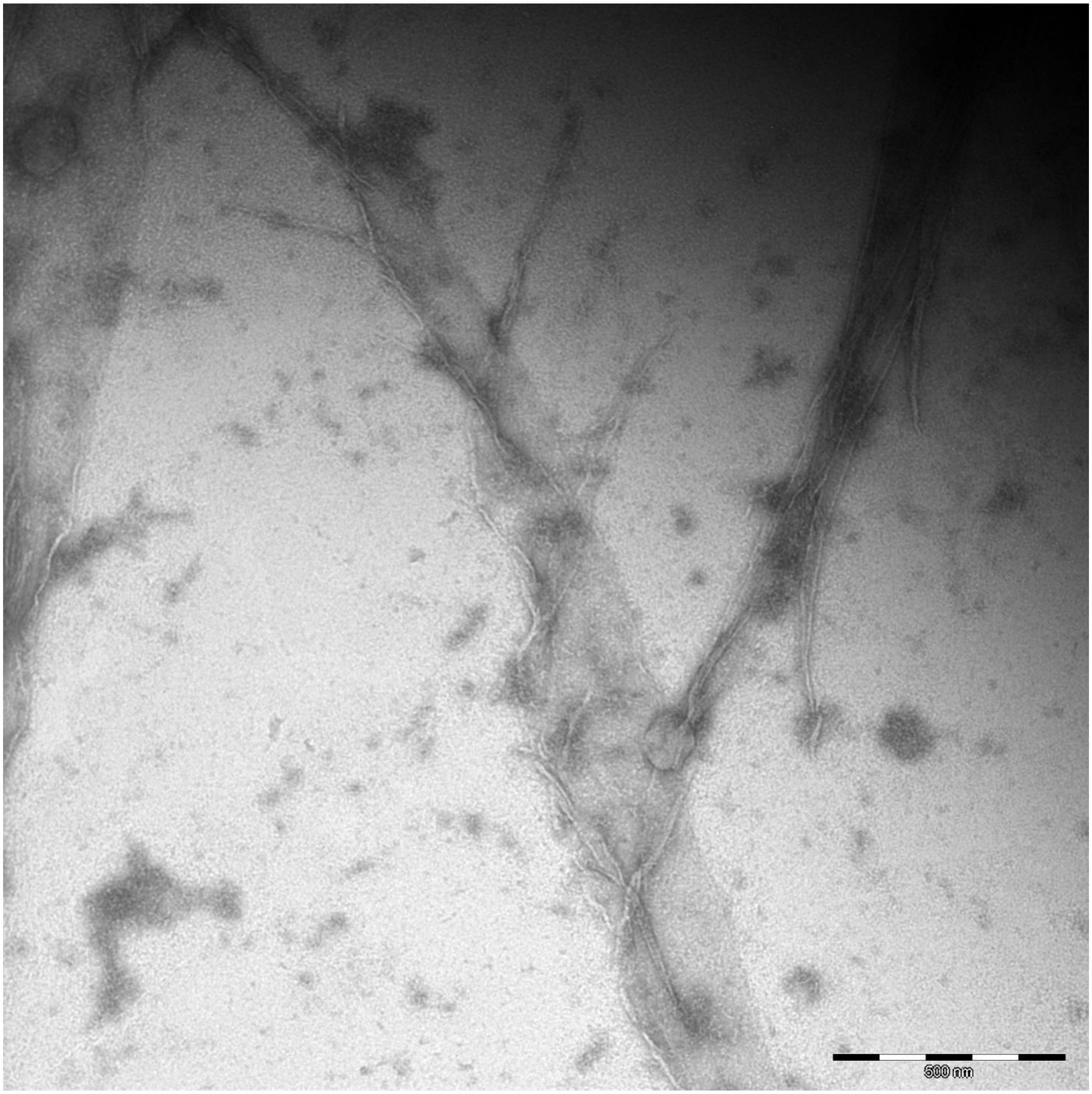
500 nm

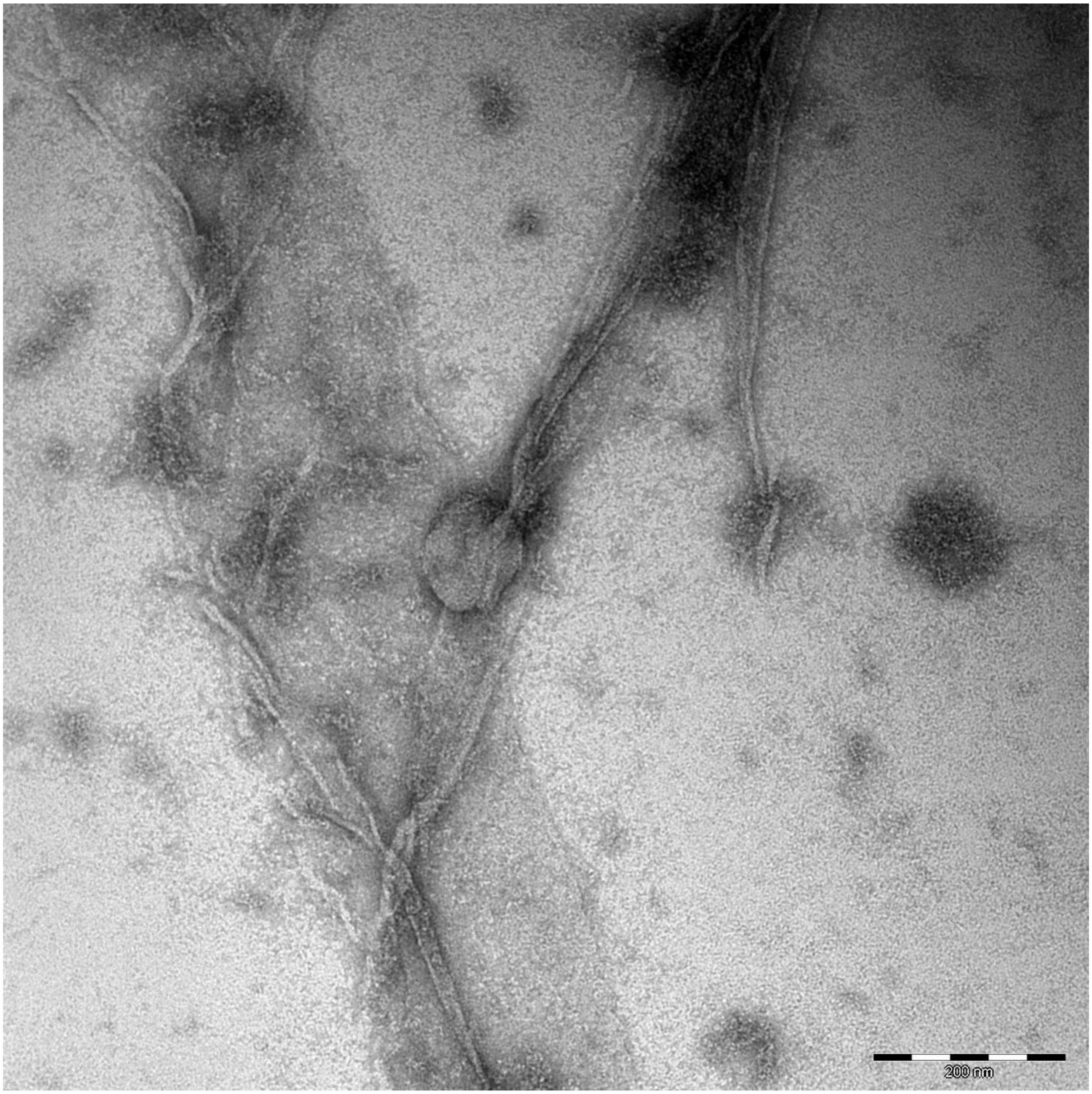
200 nm

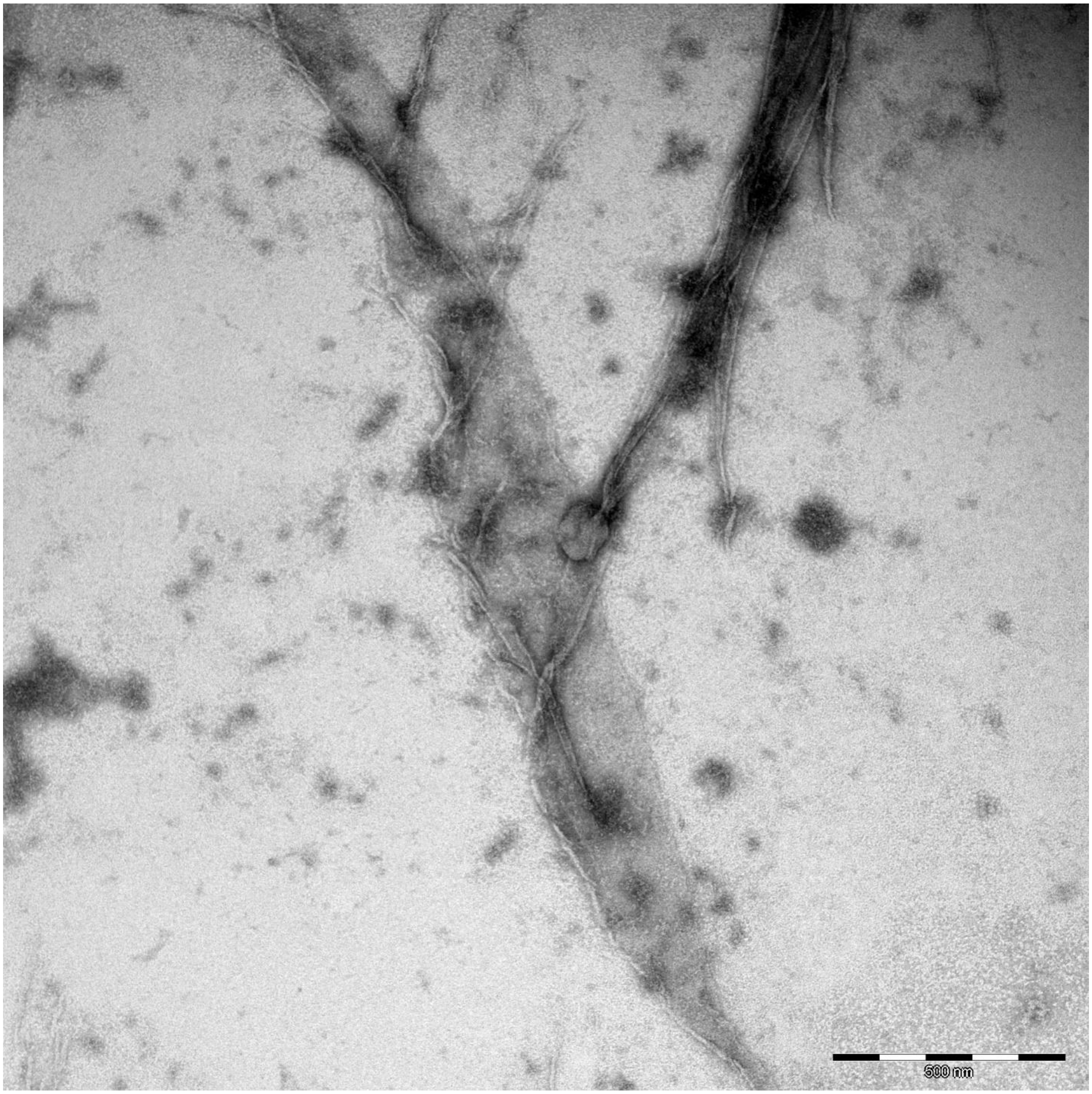
500 nm

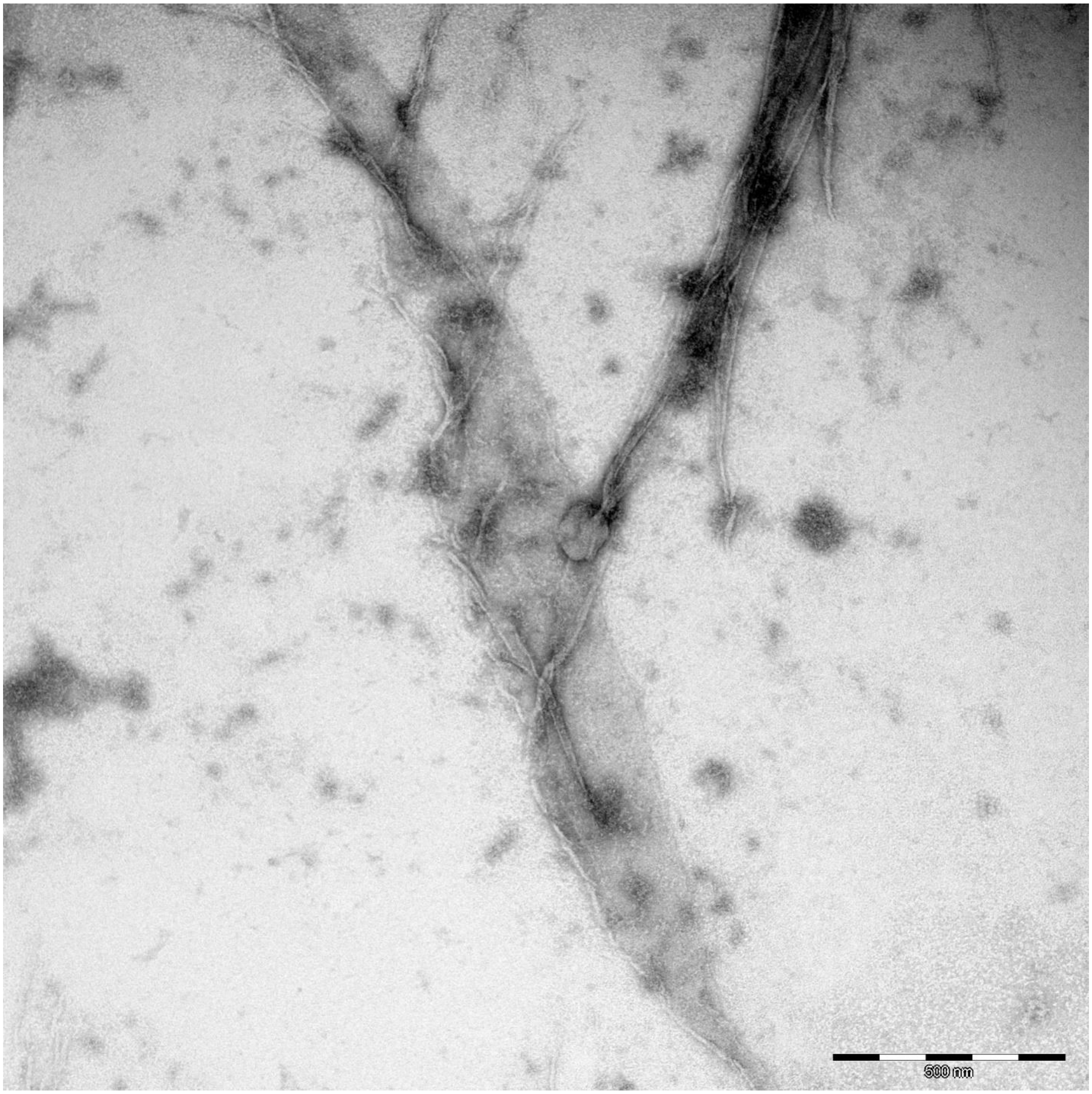
500 nm

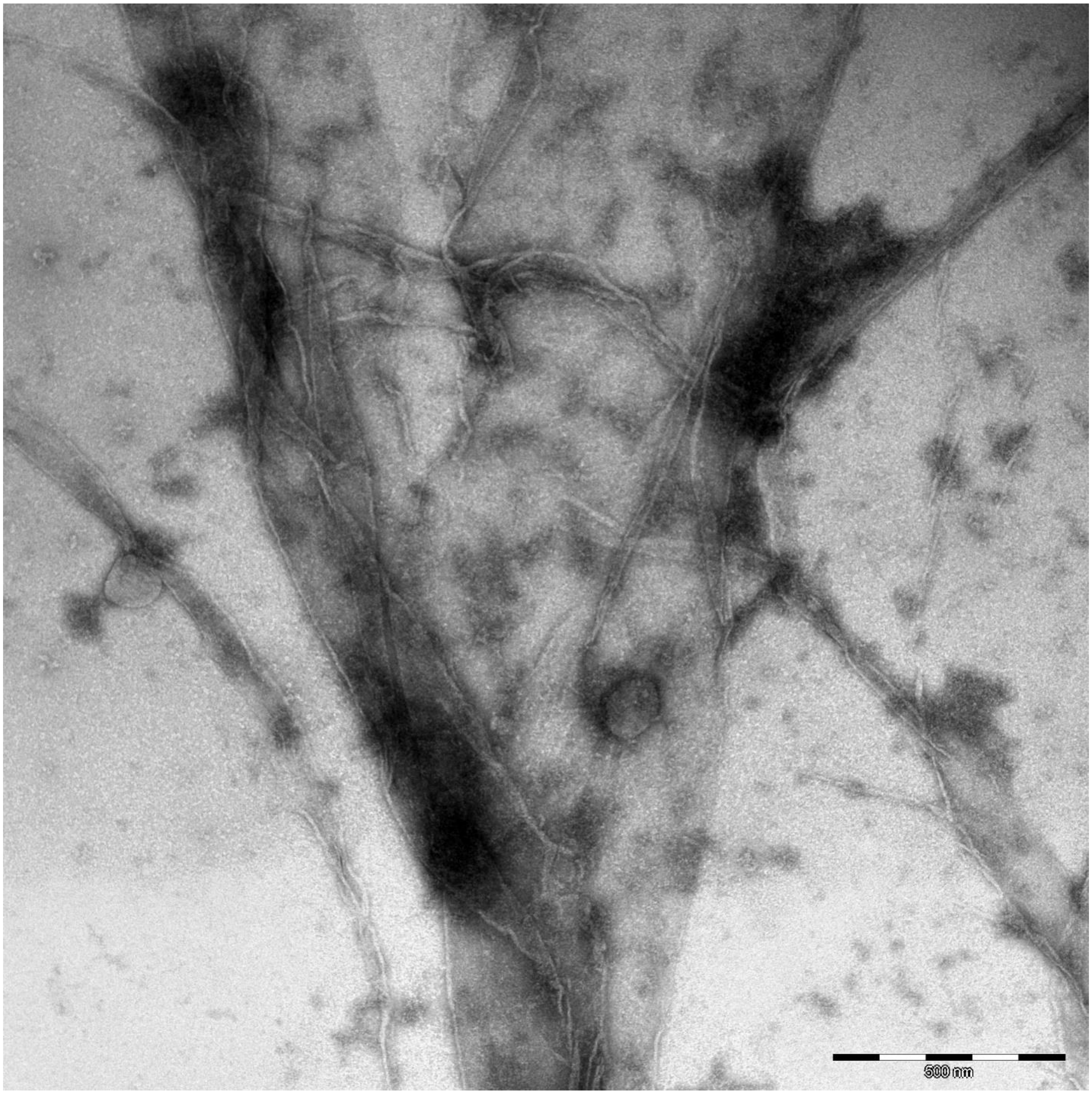
500 nm

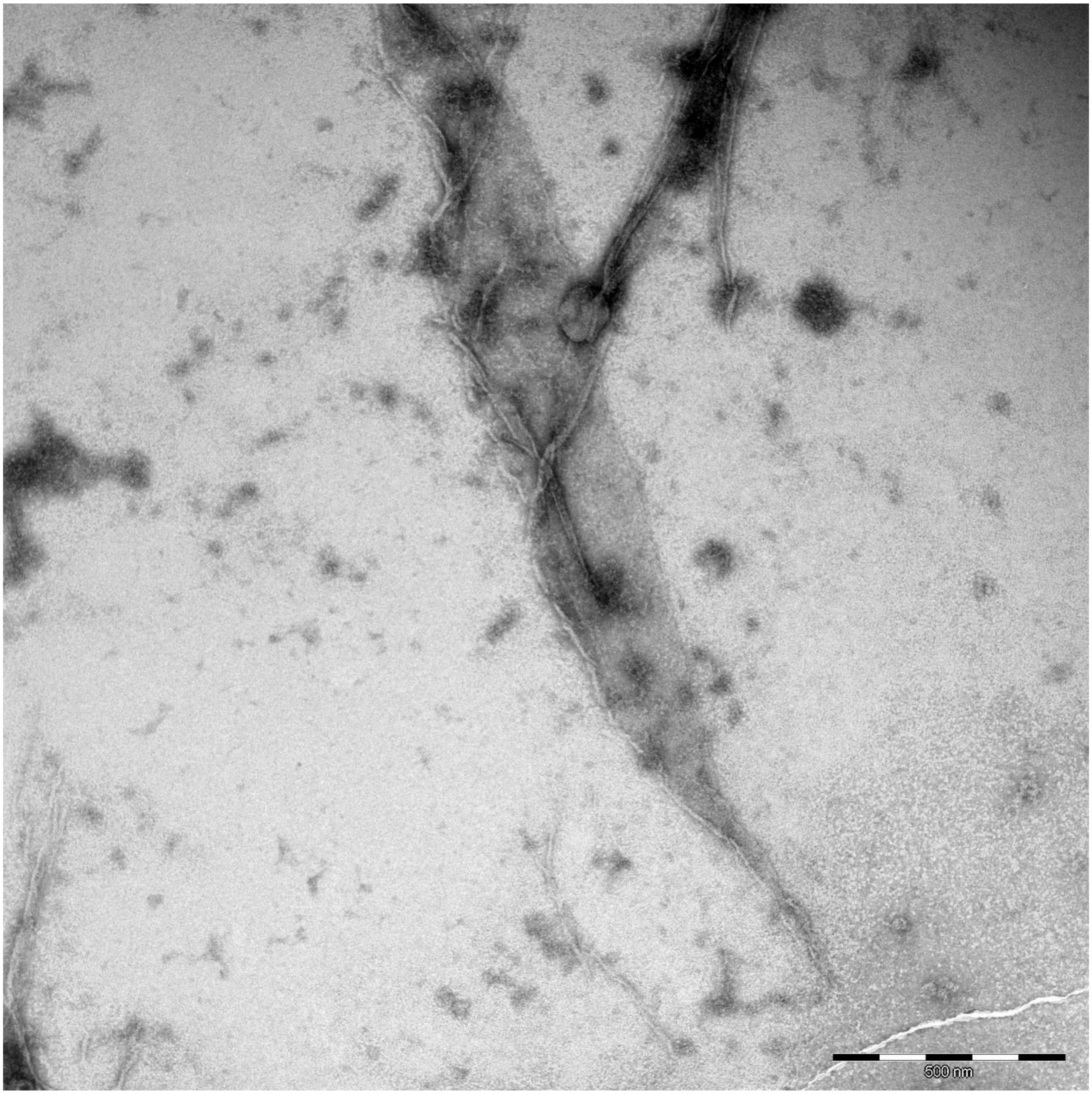
500 nm

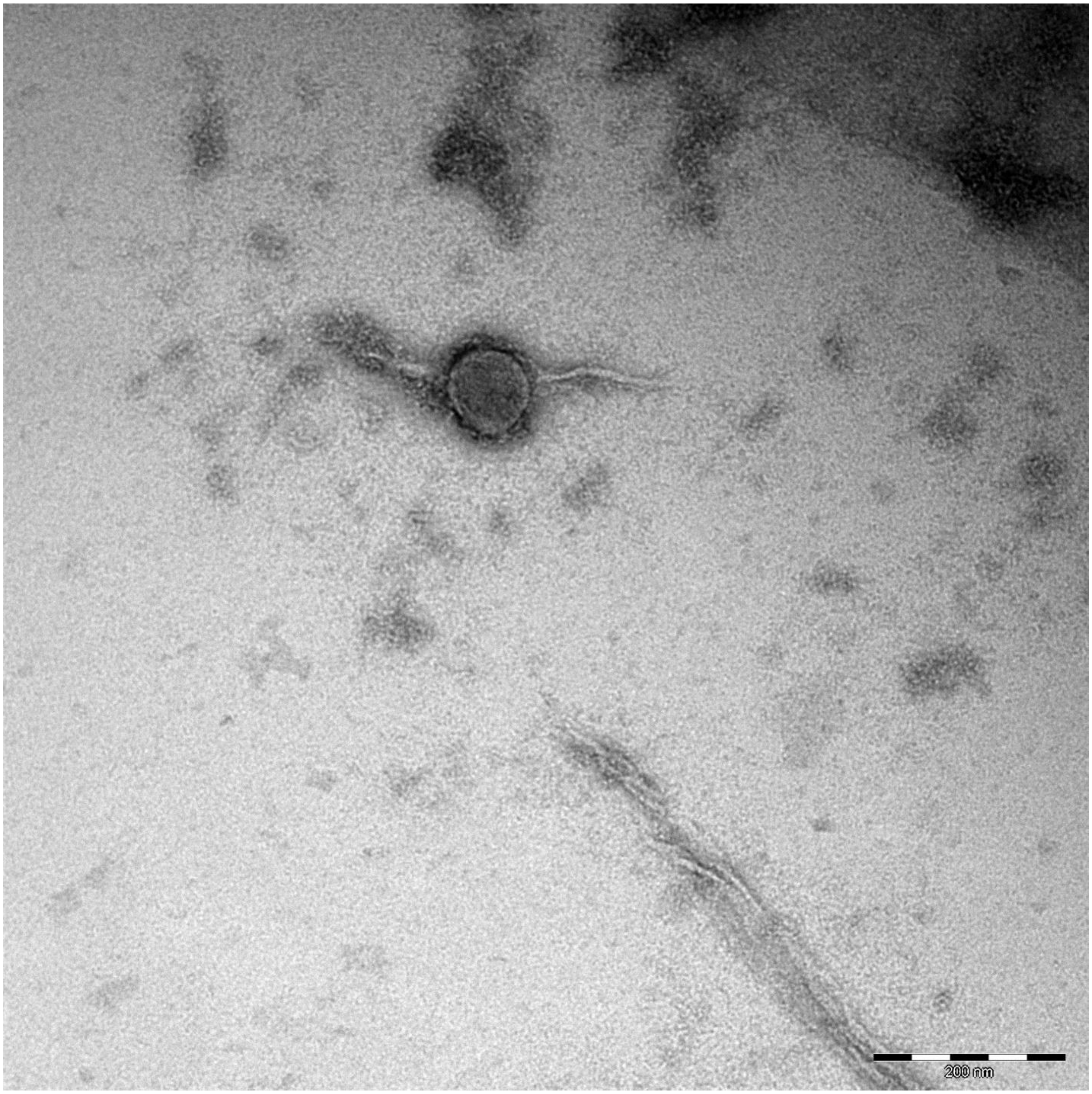
200 nm

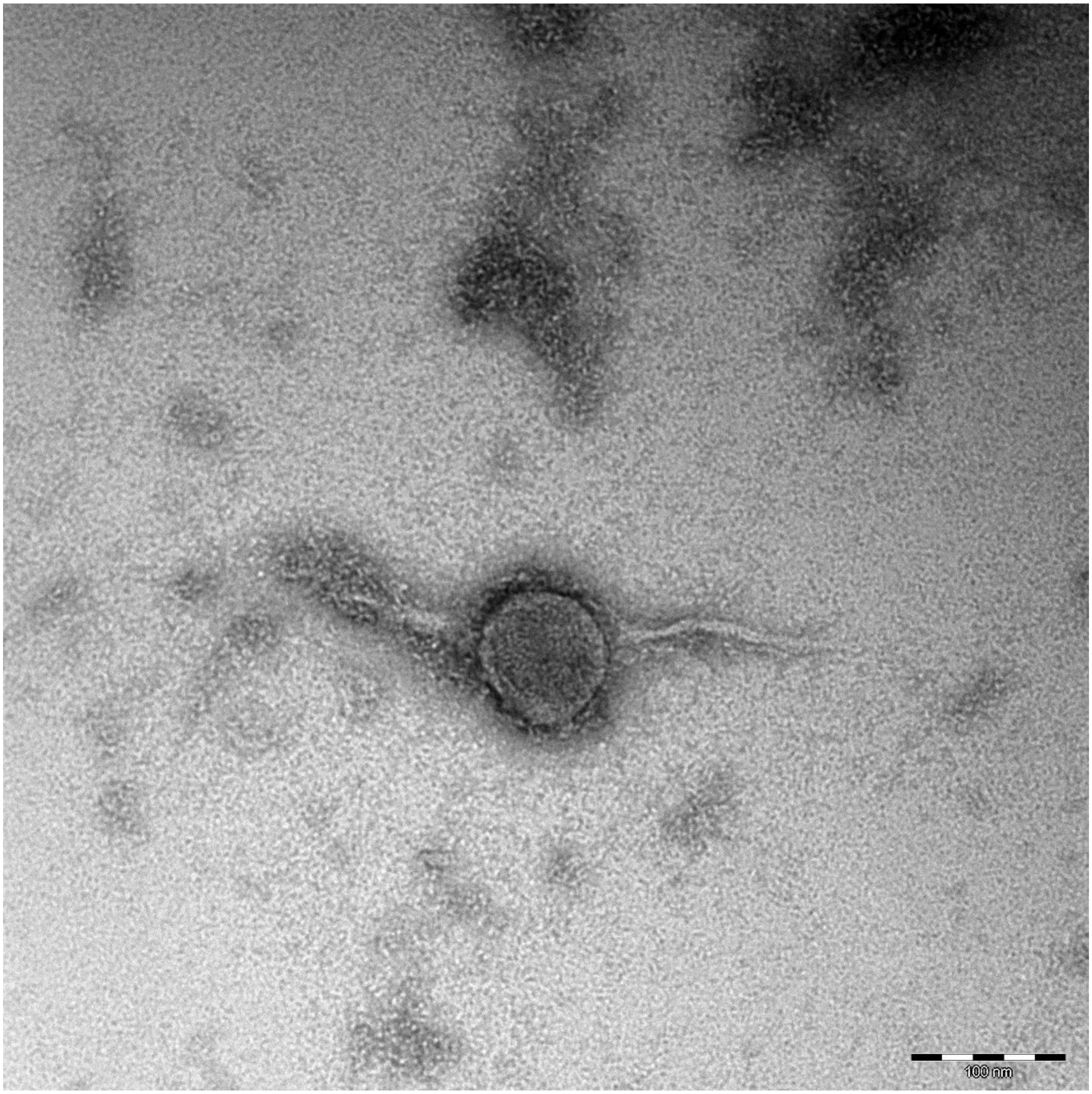
100 nm

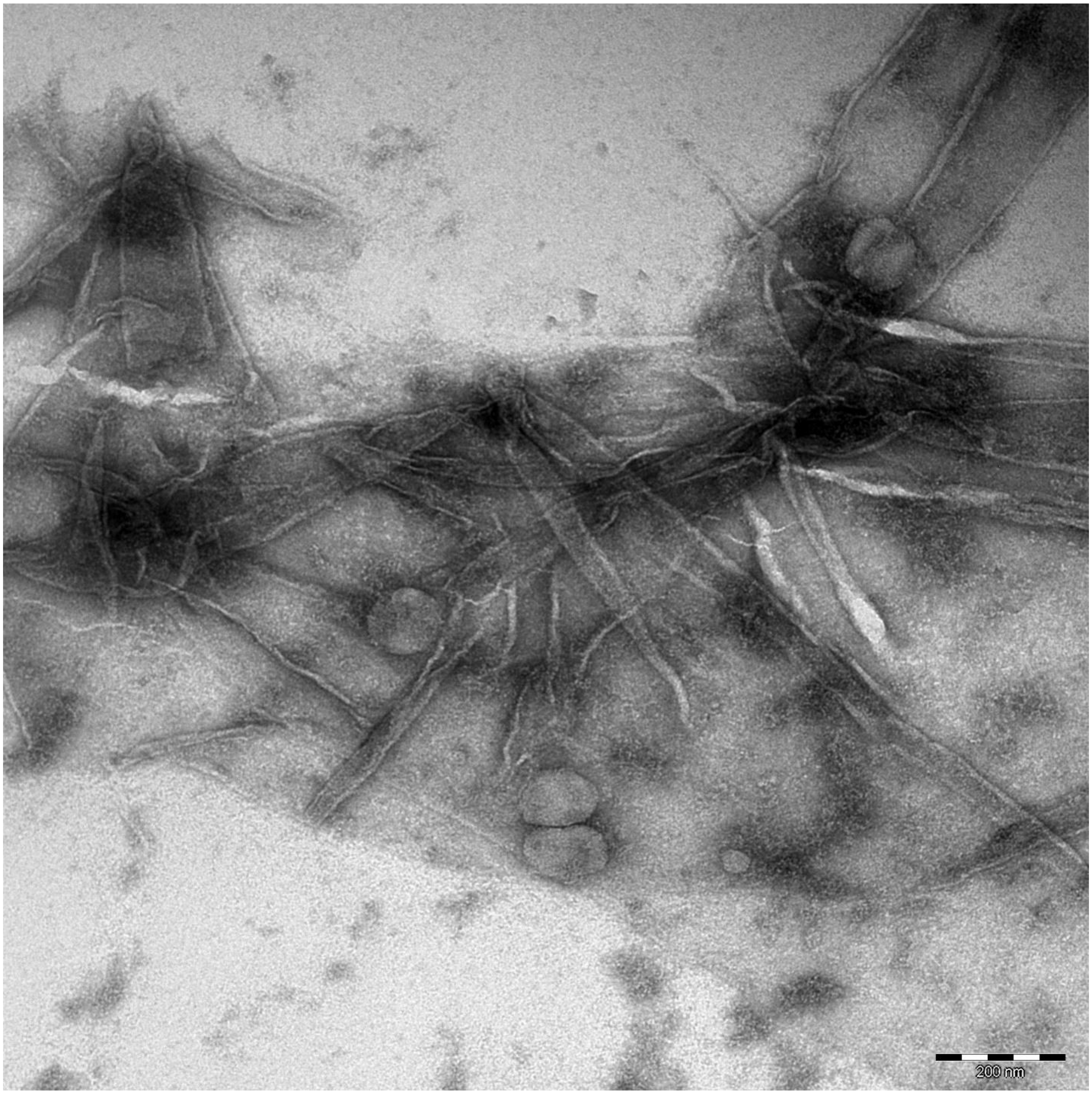
200 nm

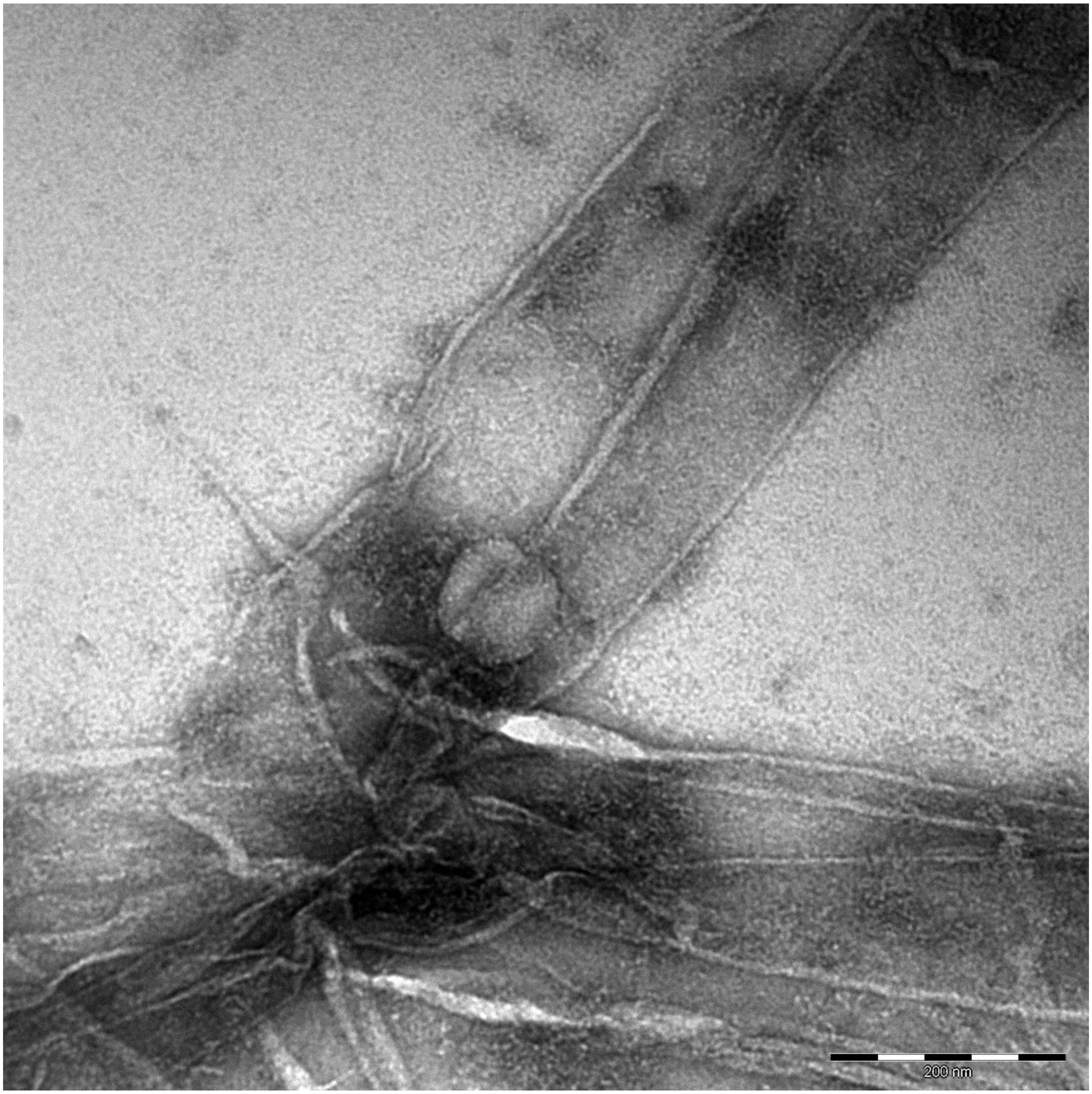
200 nm

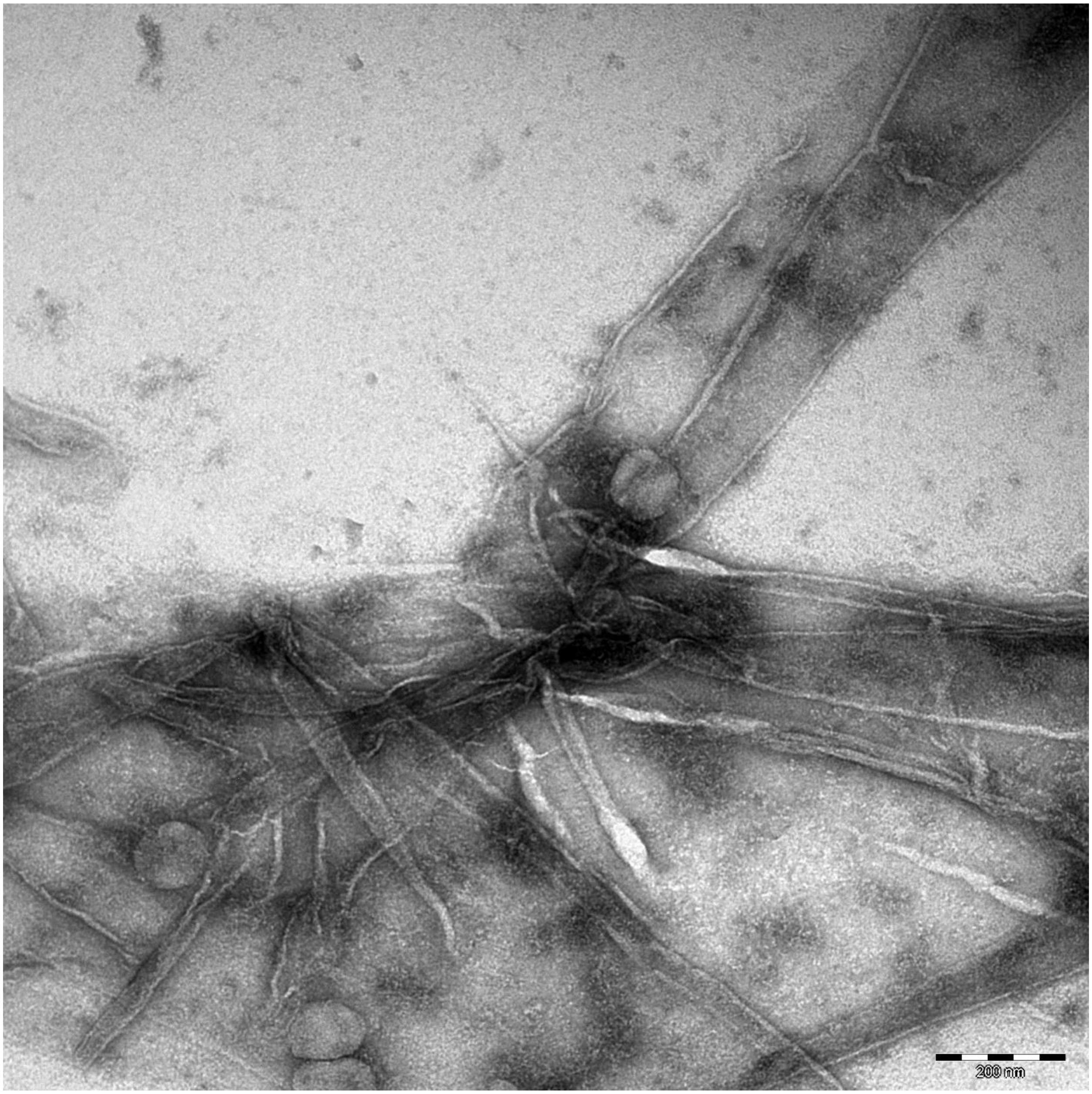
200 nm

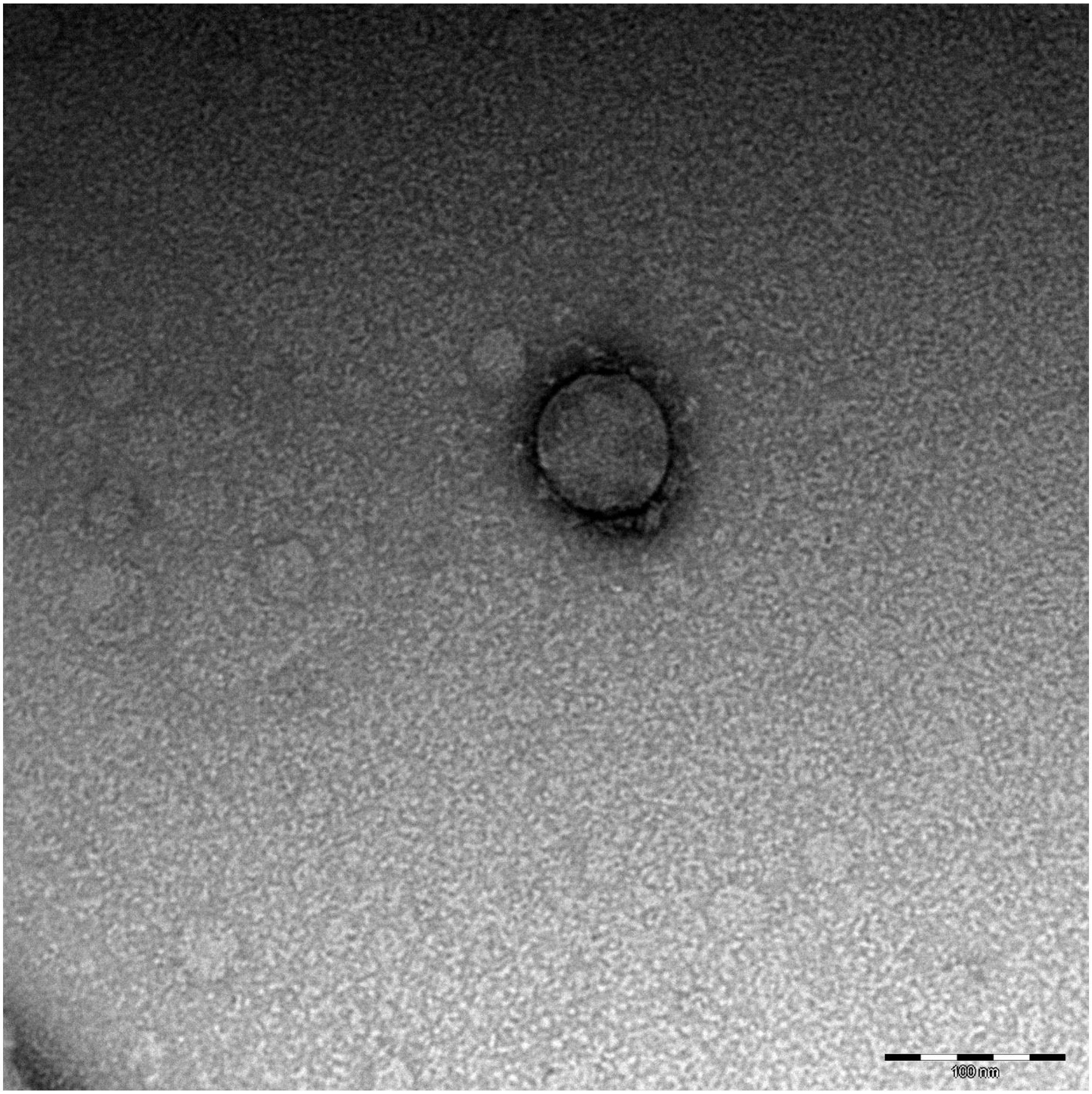
100 nm

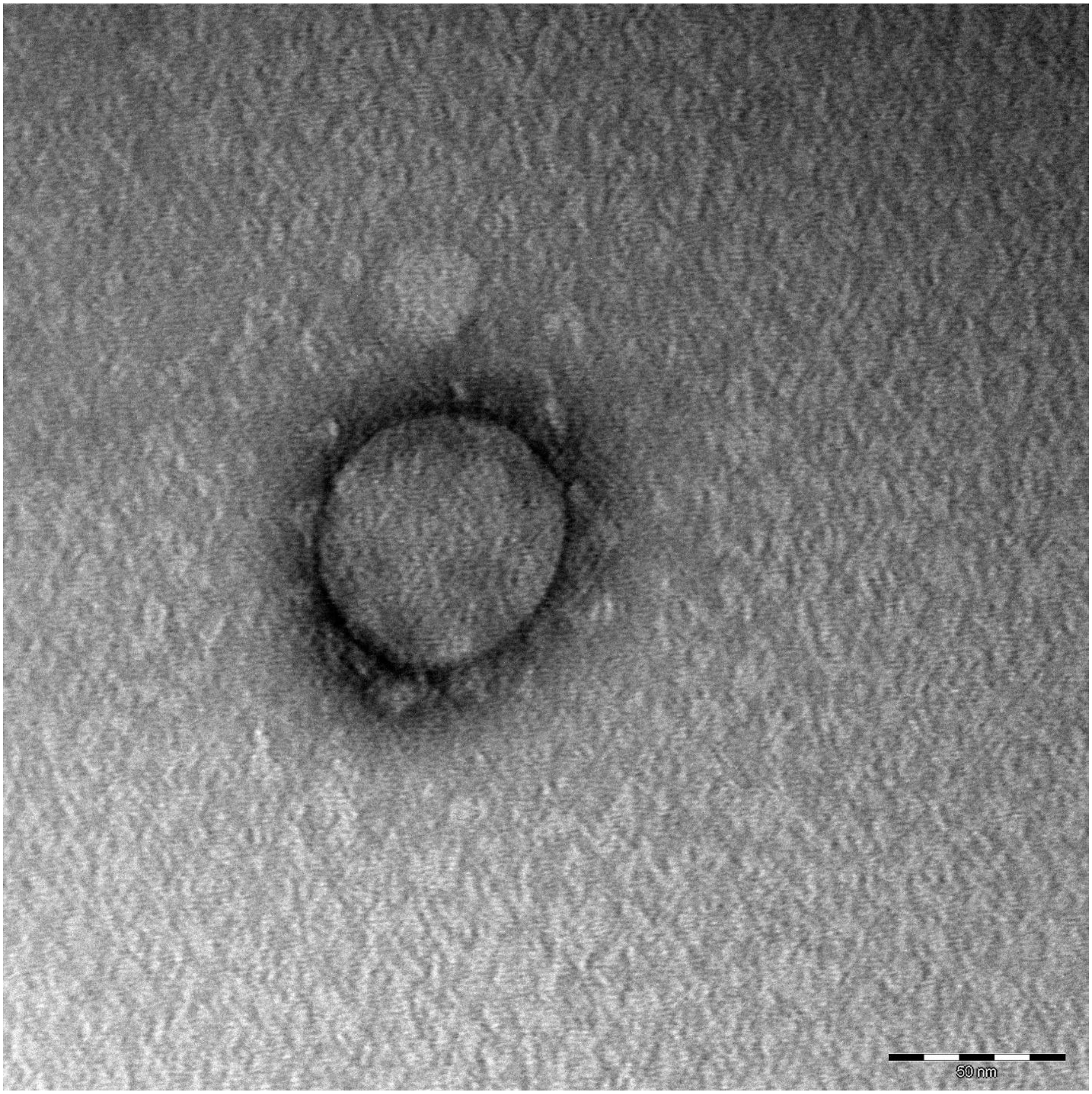
50 nm

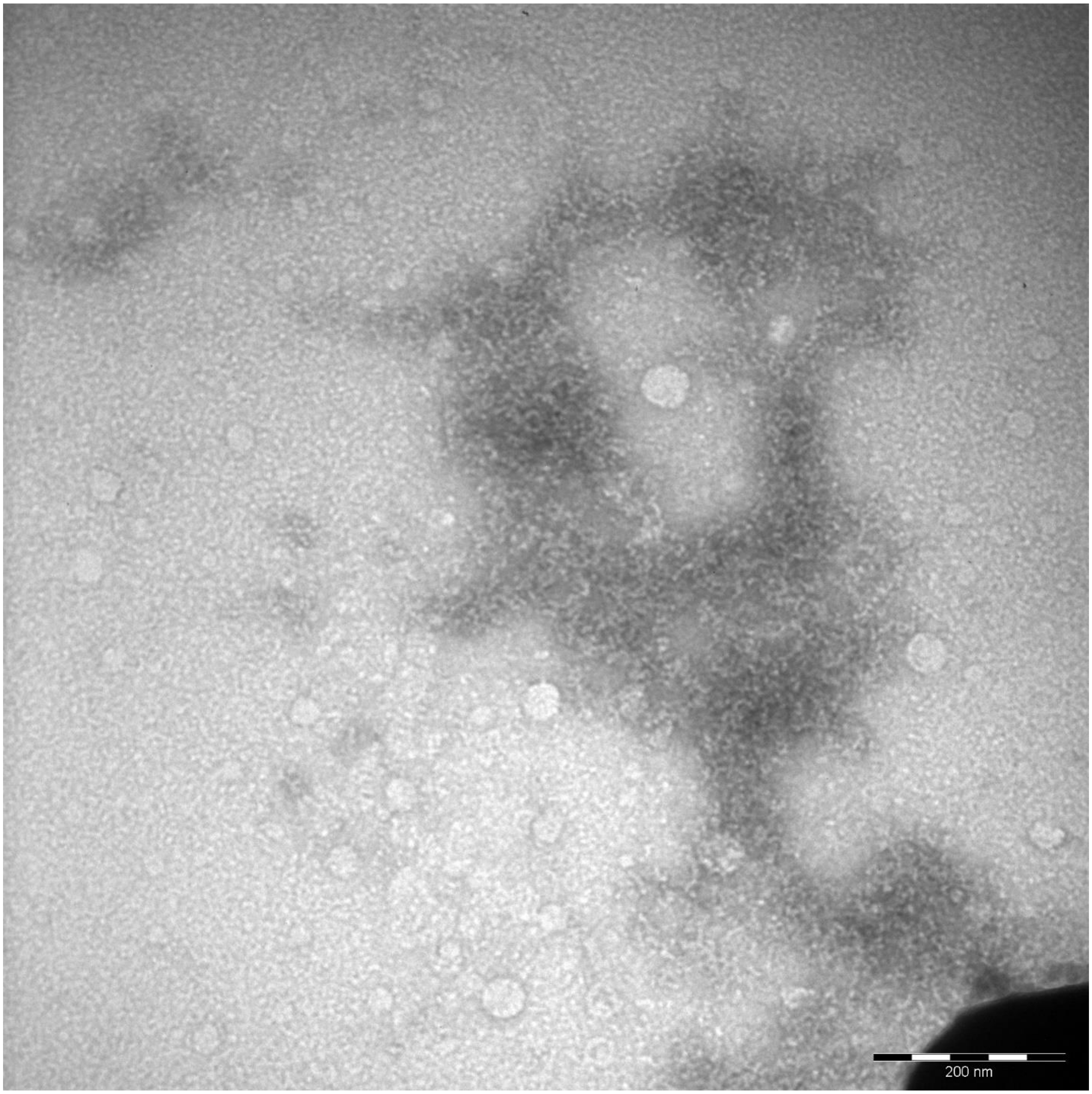
200 nm

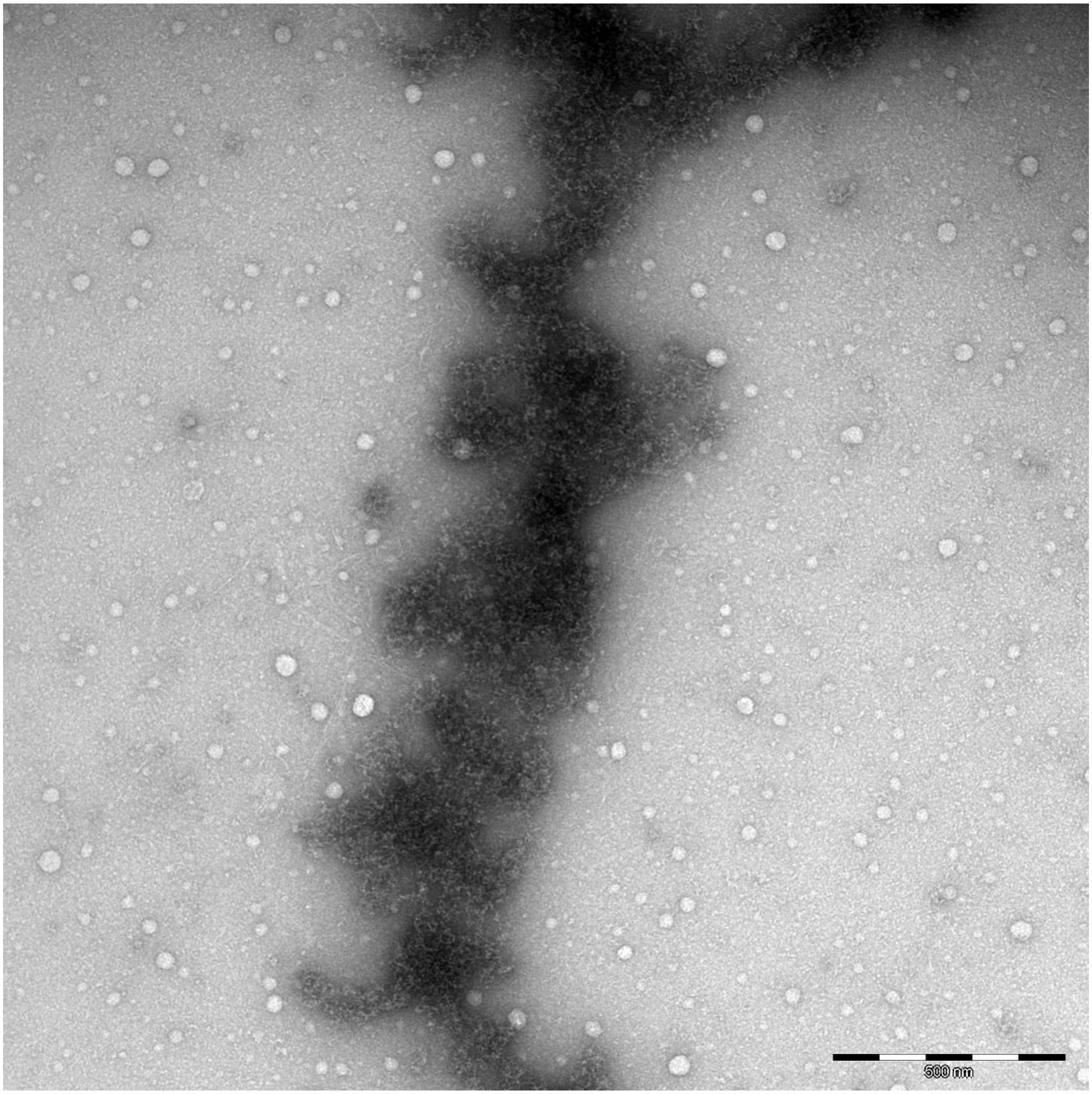
500 nm

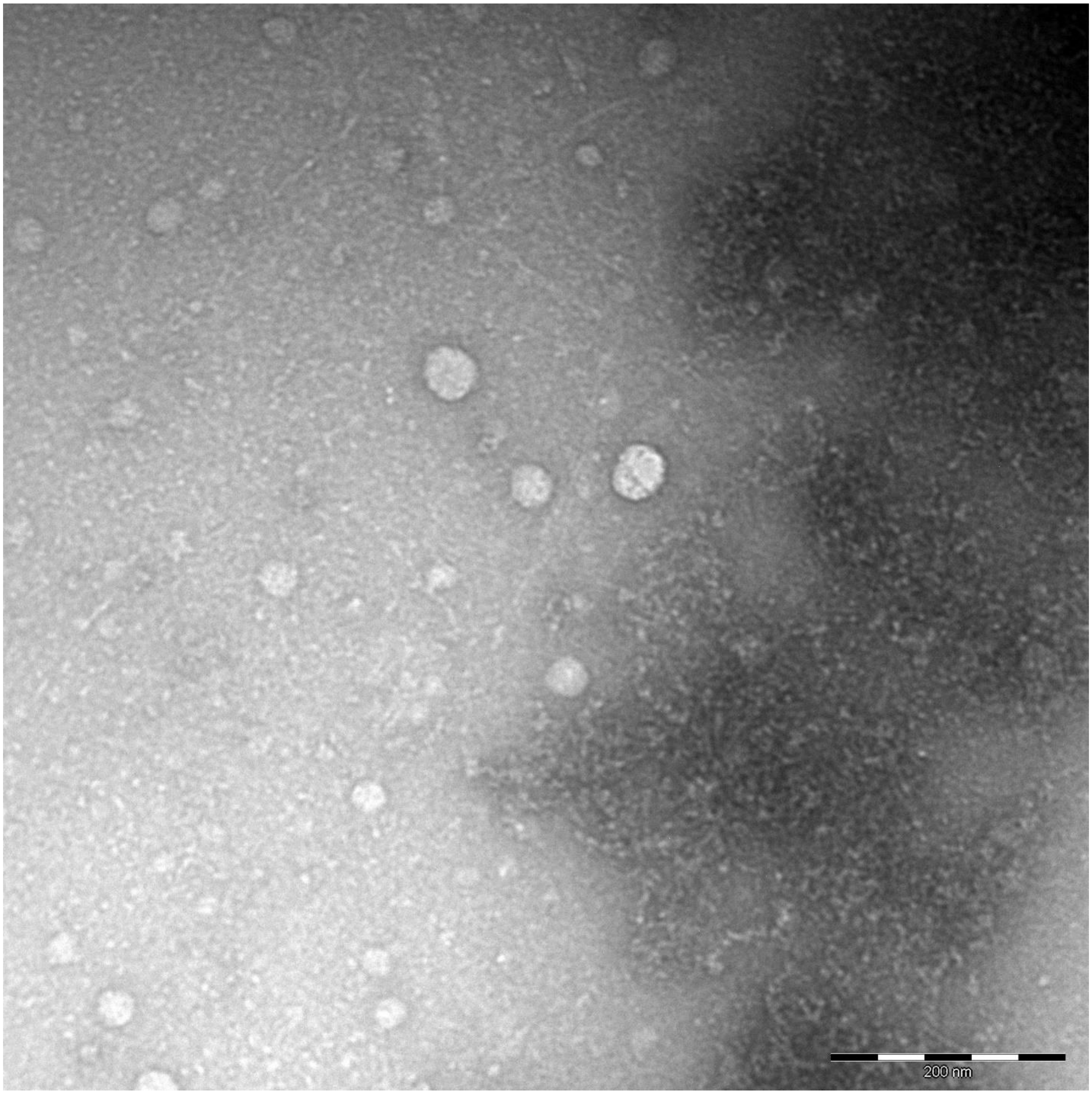
200 nm

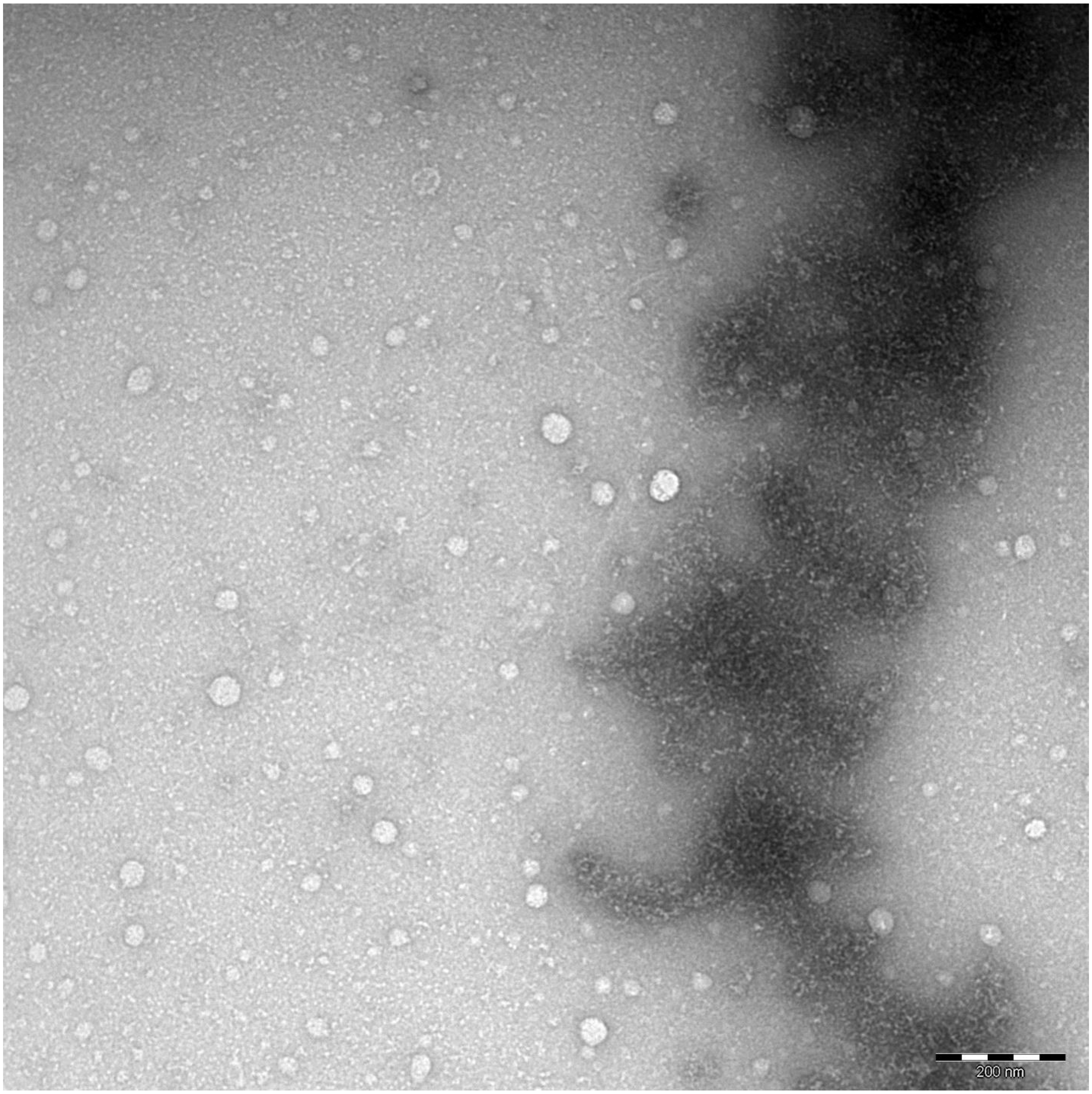
200 nm

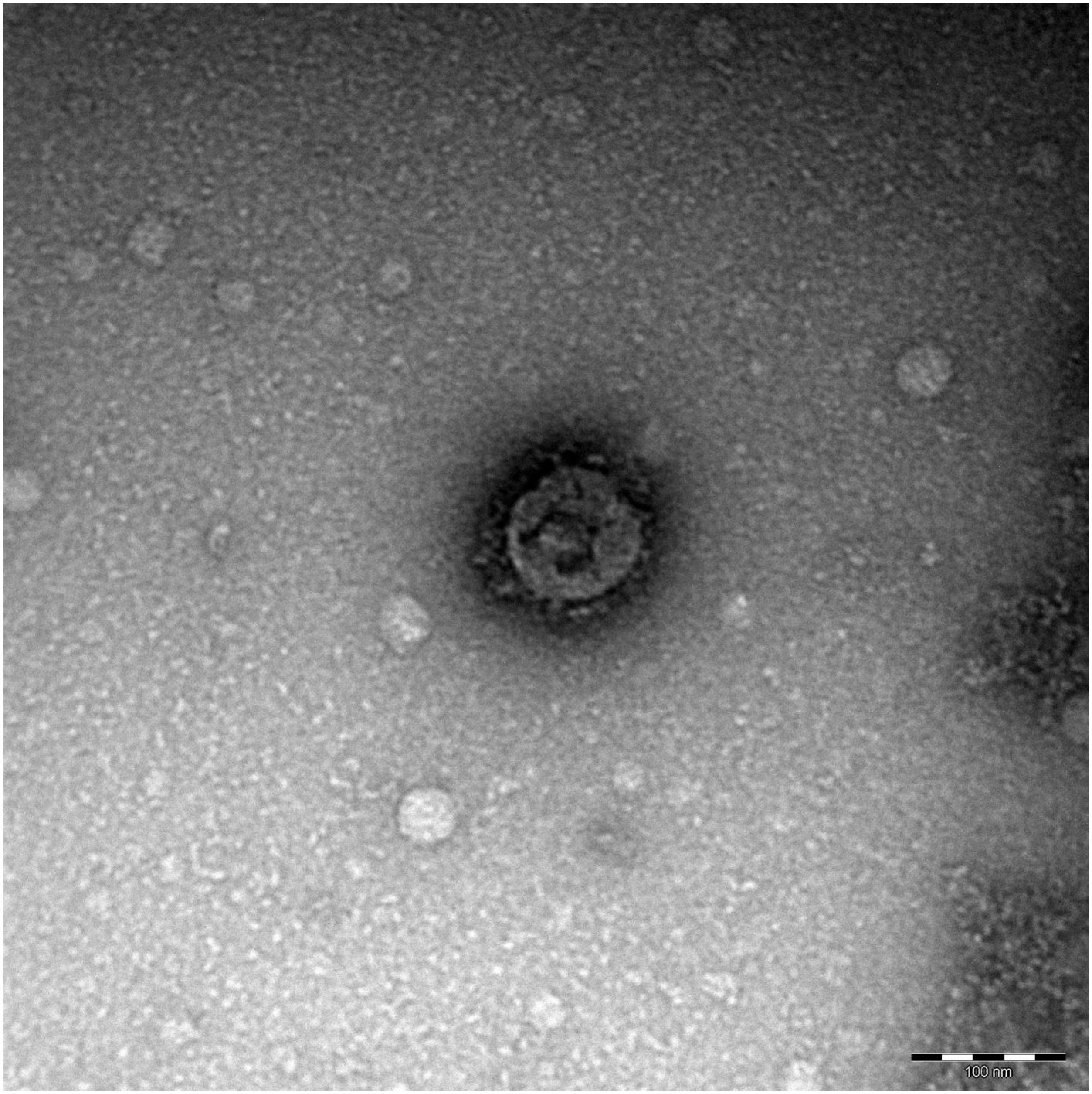
100 nm

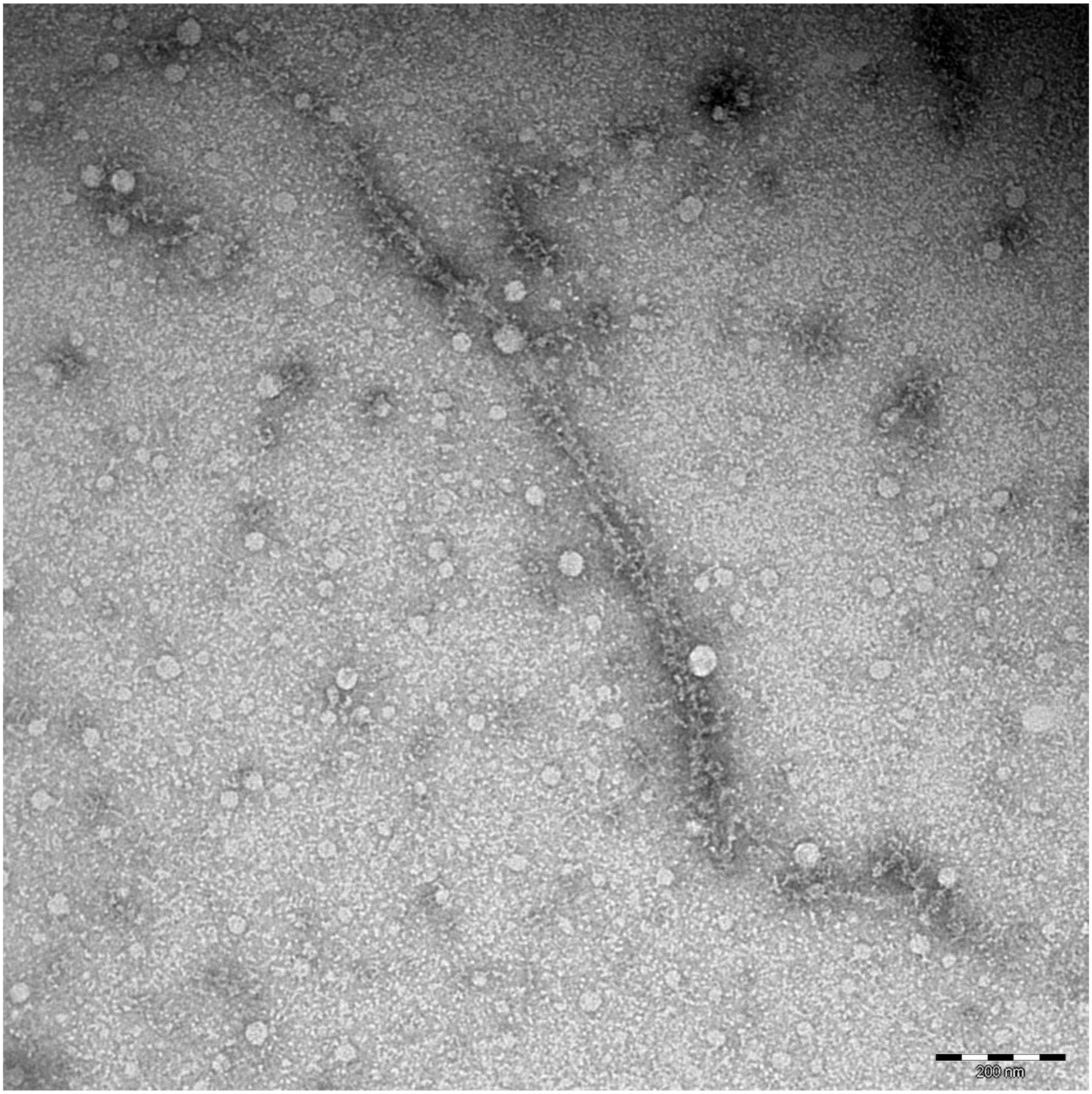
200 nm

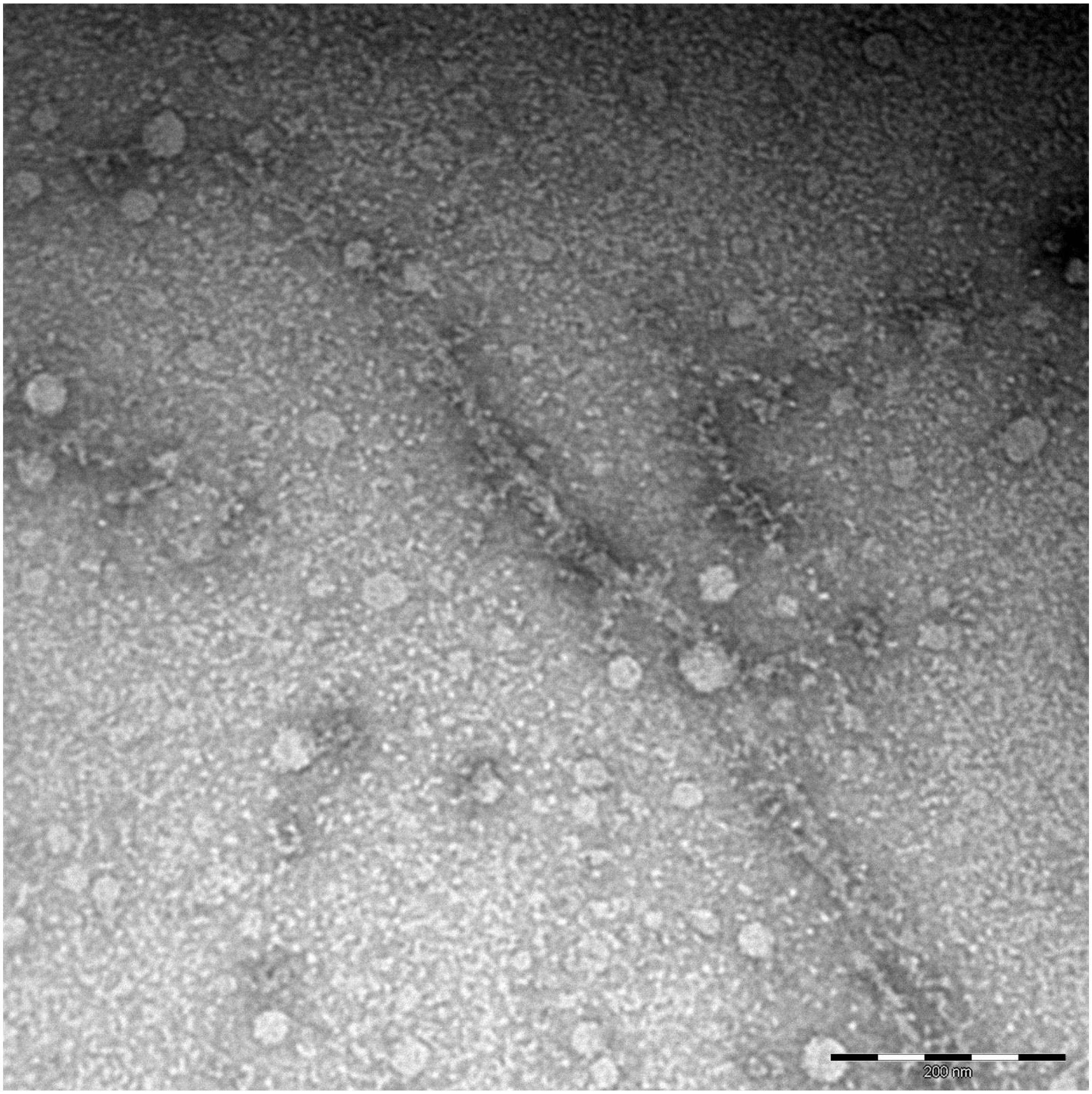
200 nm

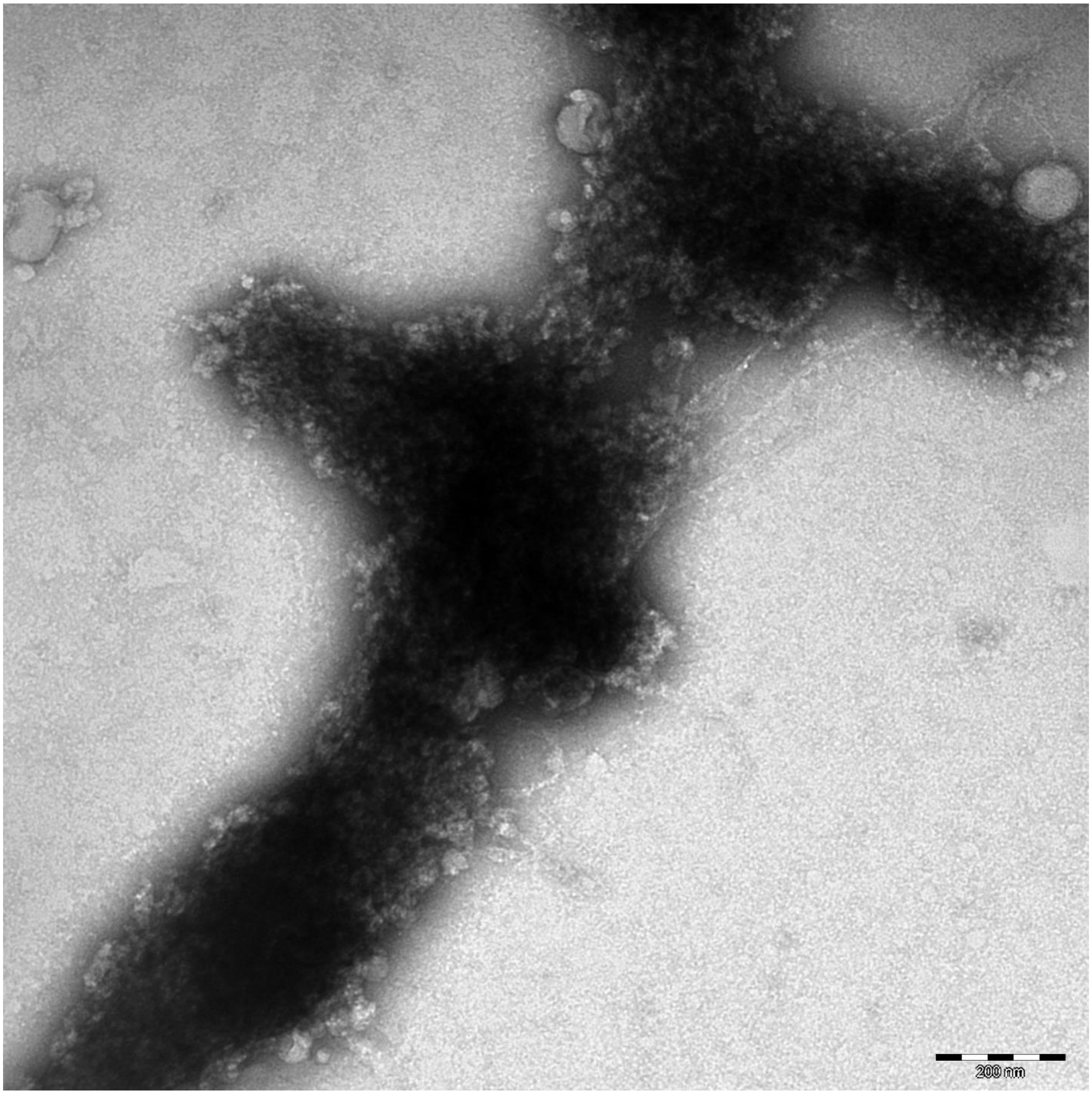
200 nm

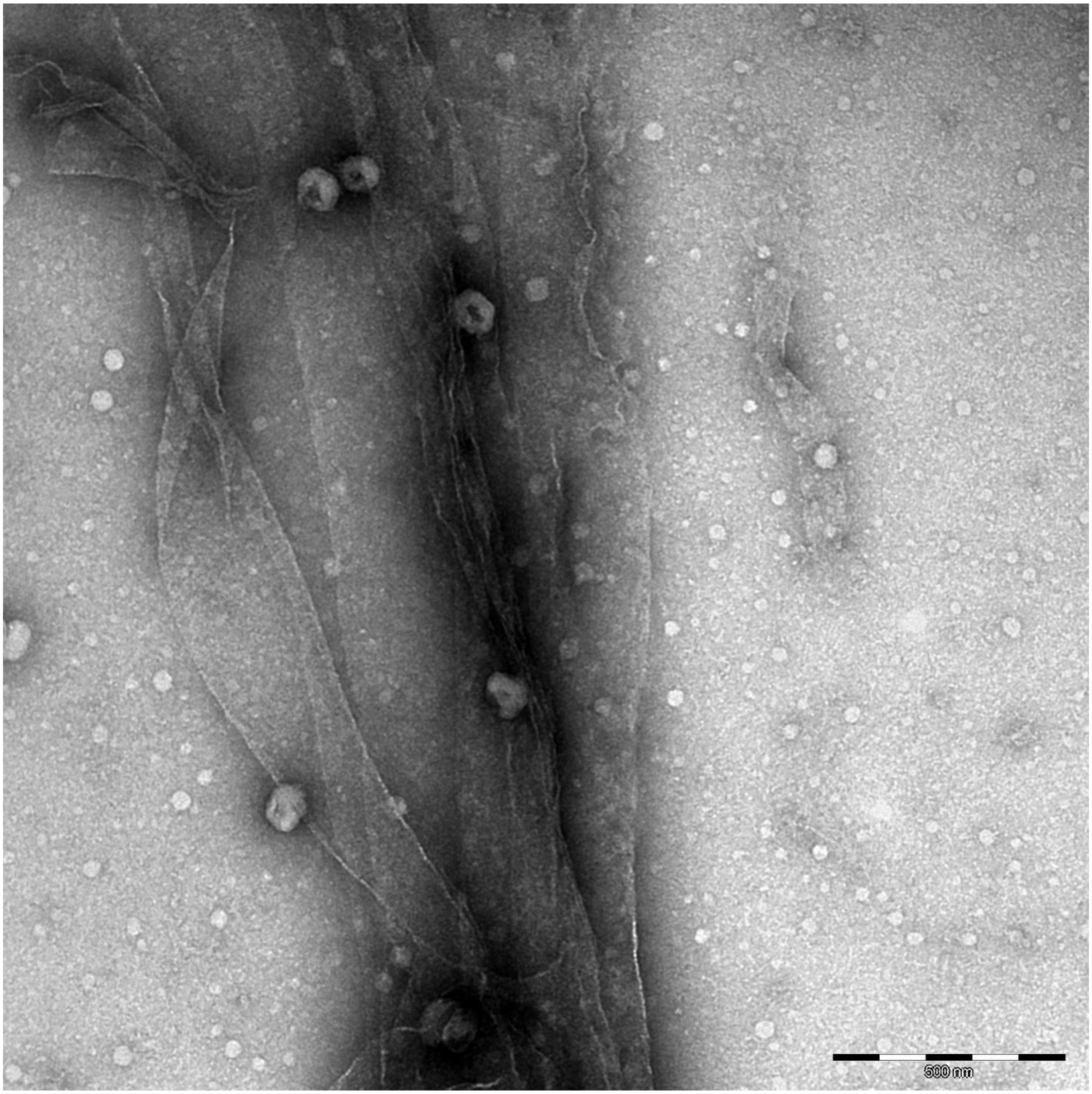
500 nm

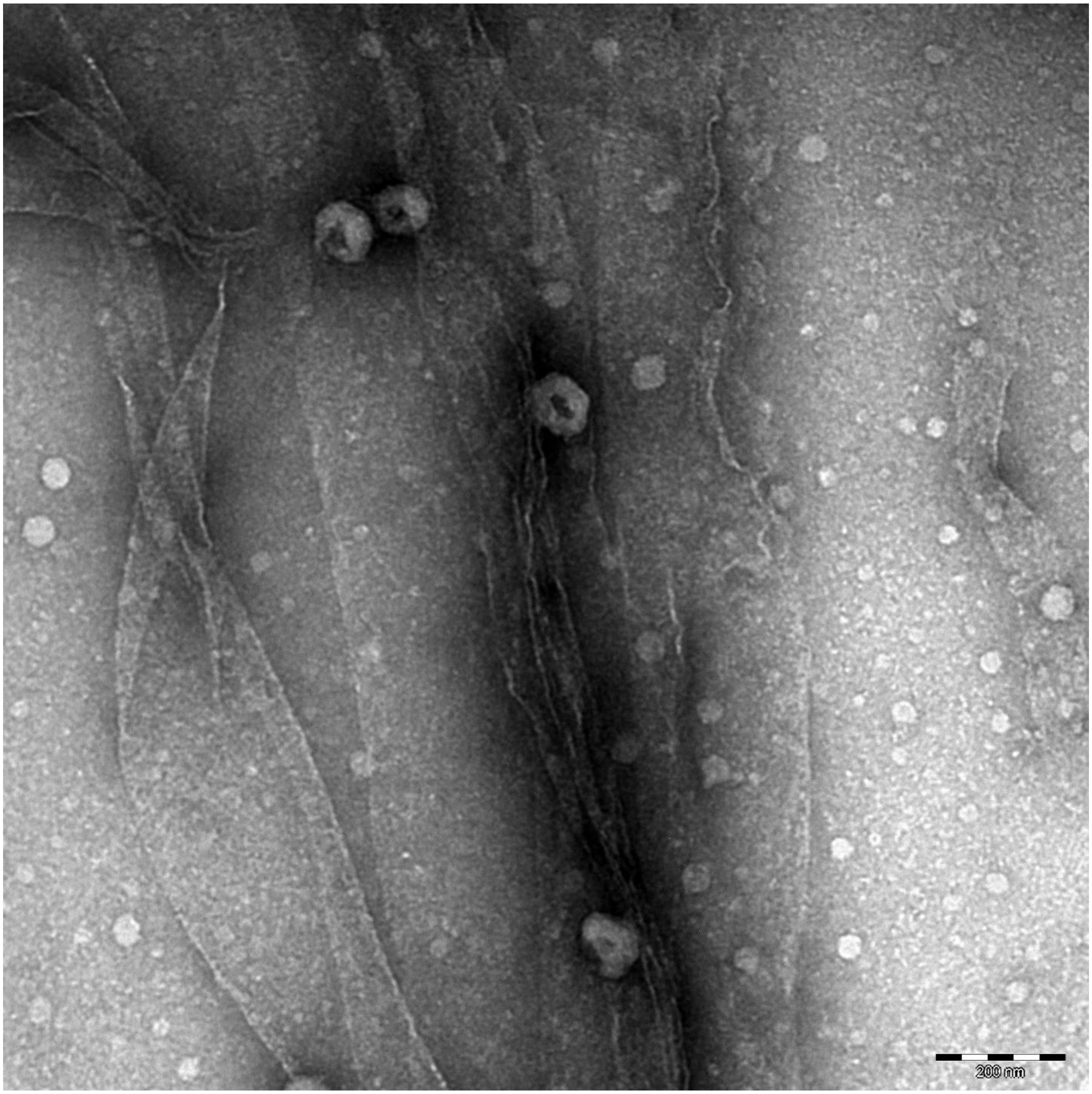
200 nm

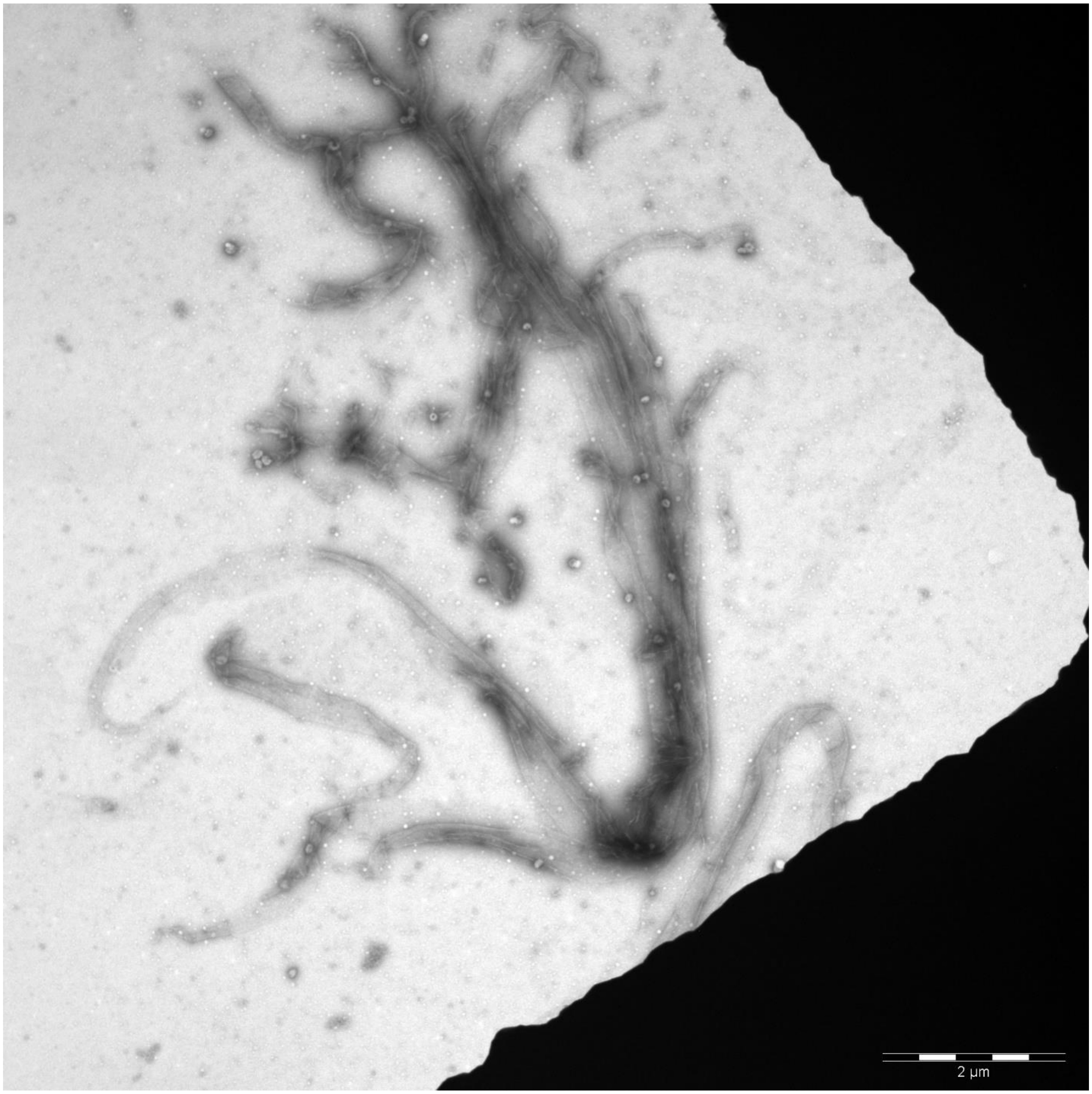
2 µm

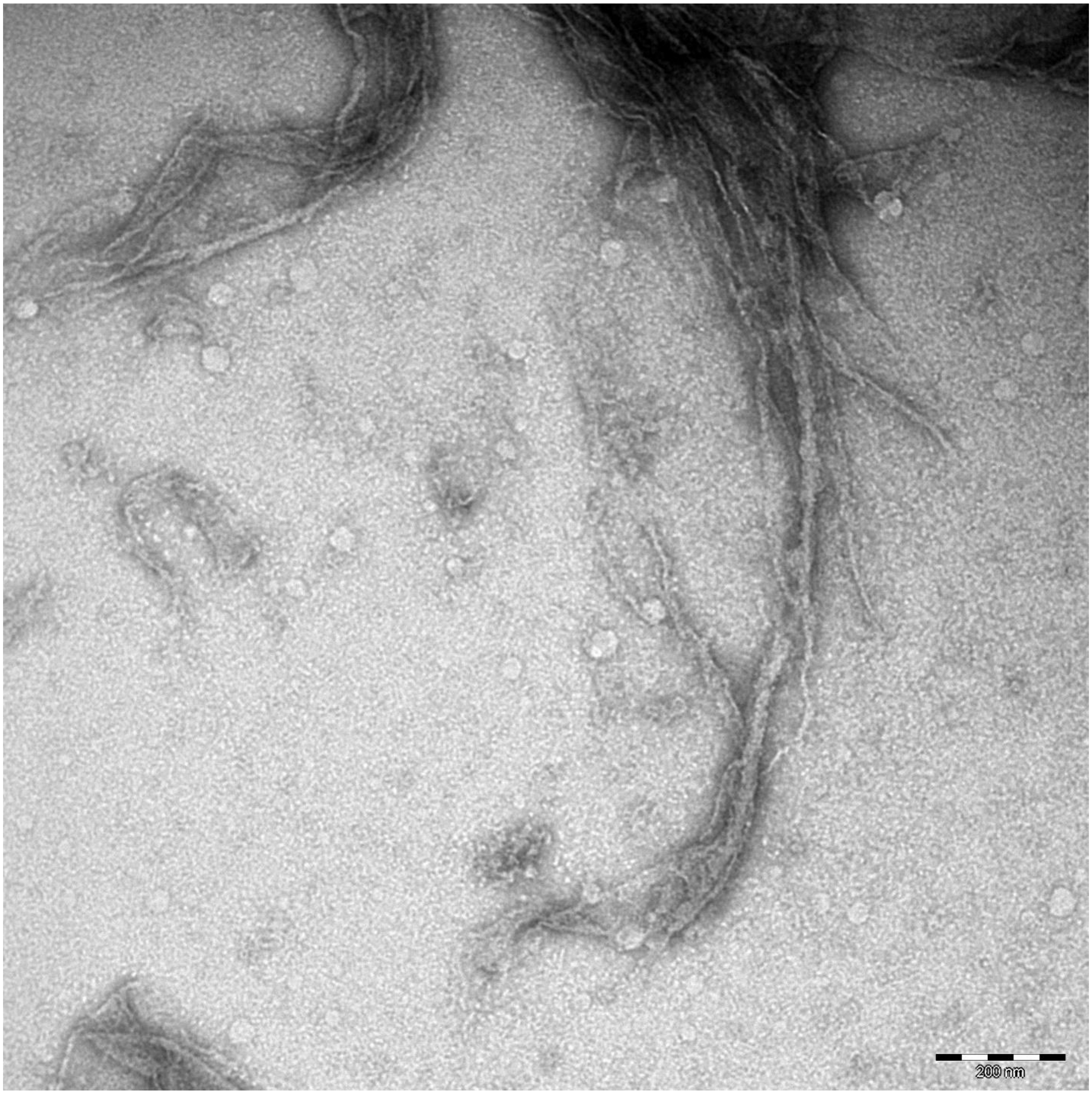
200 nm

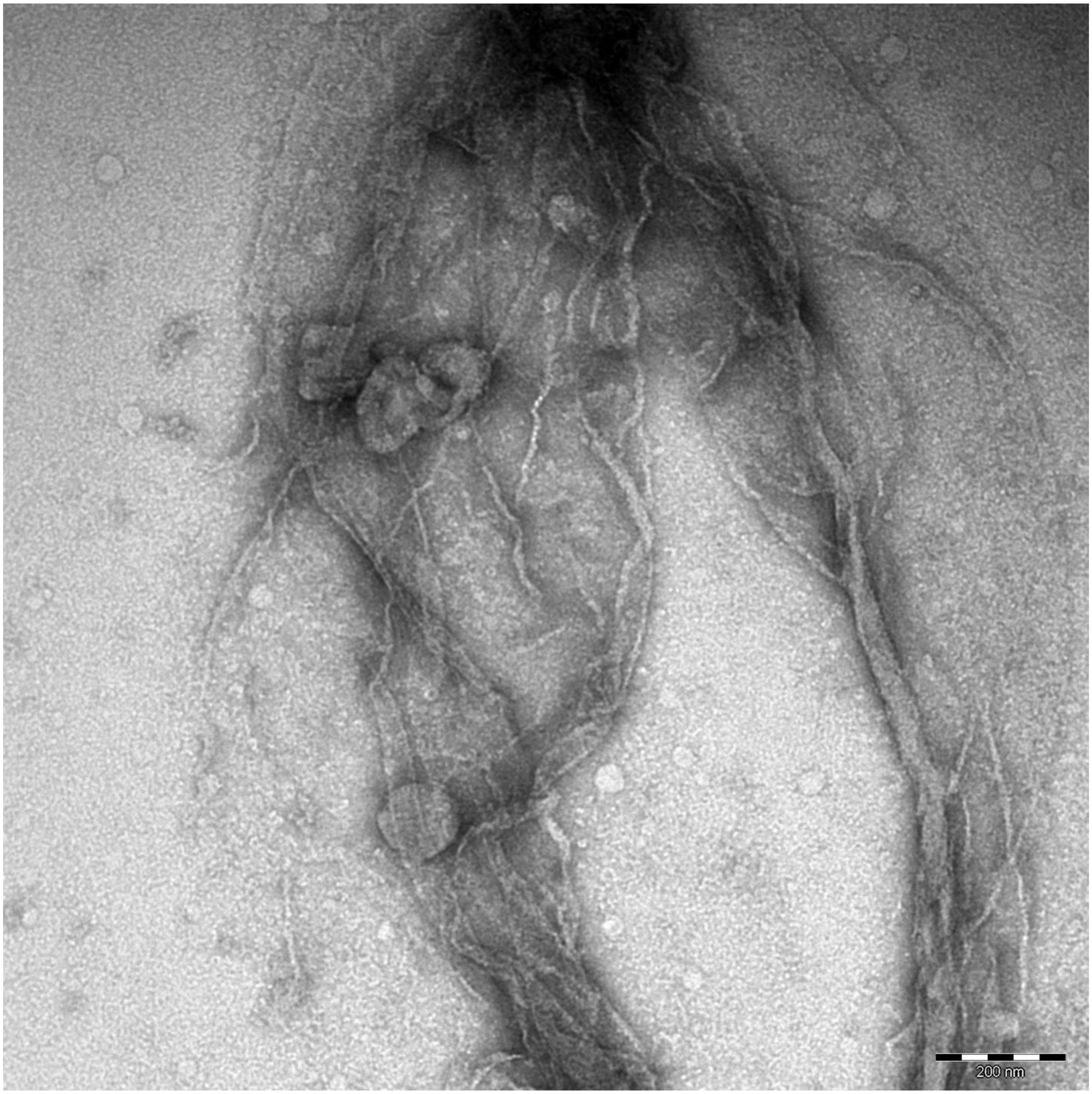
200 nm

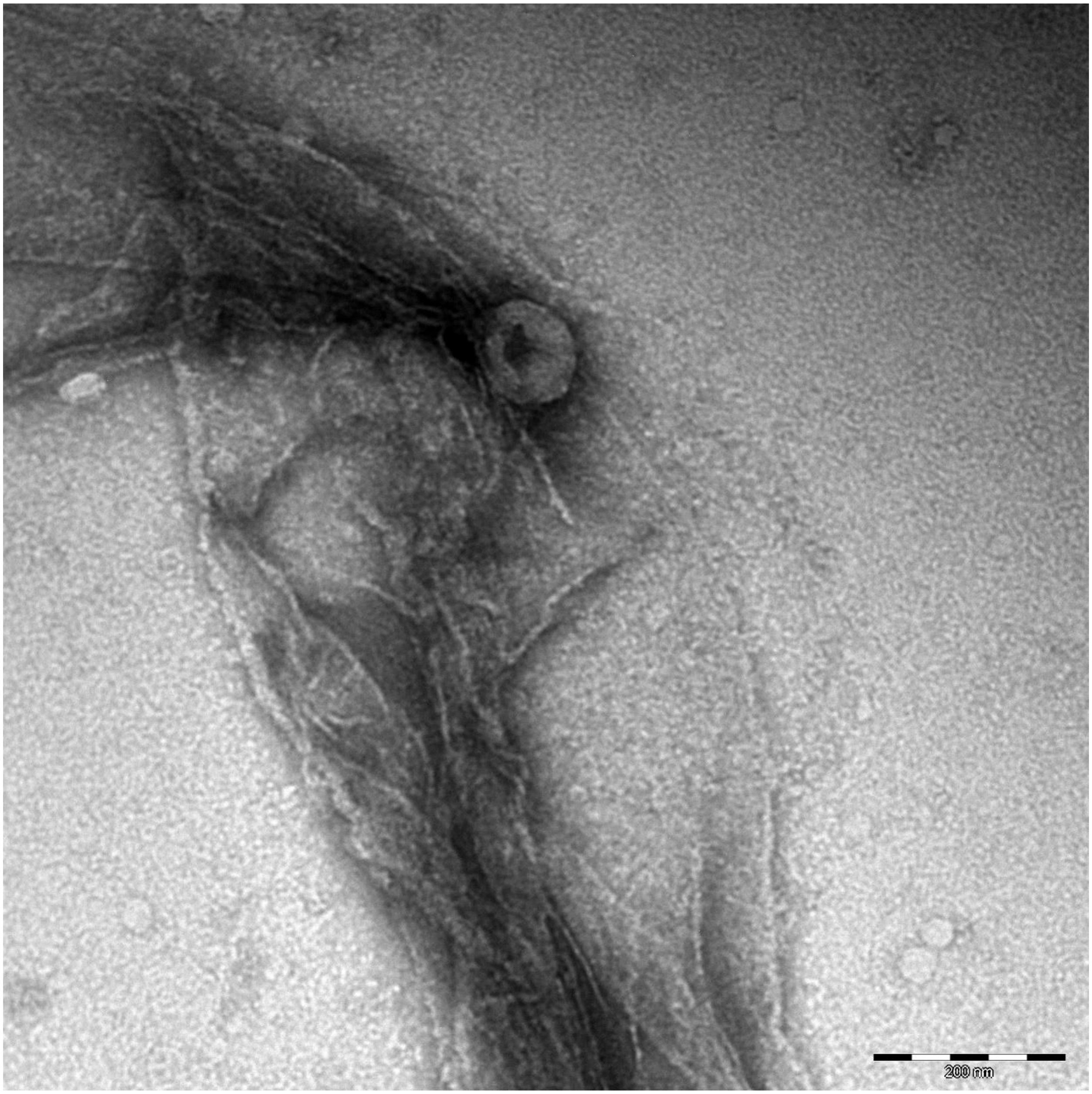
200 nm

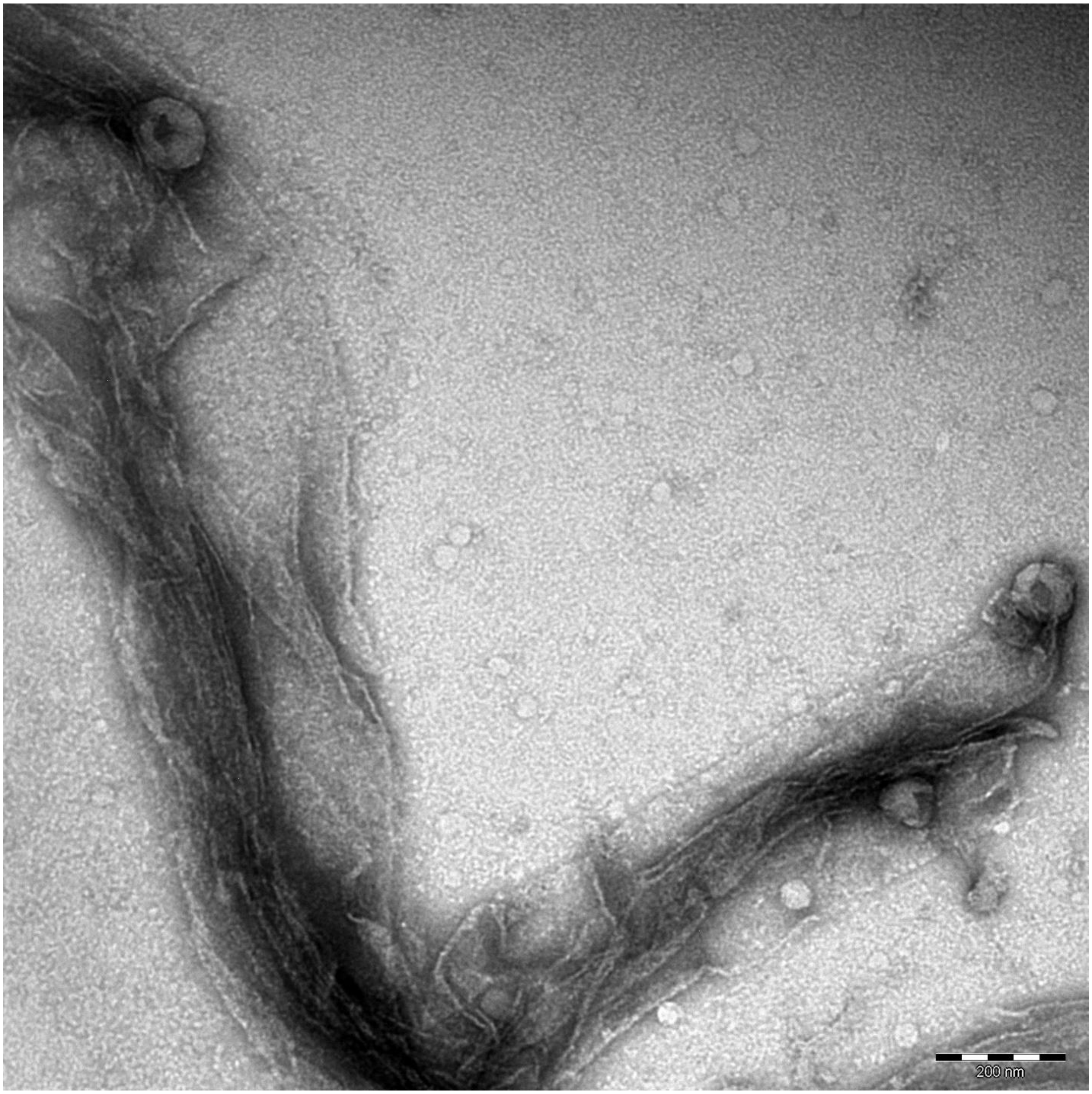
200 nm

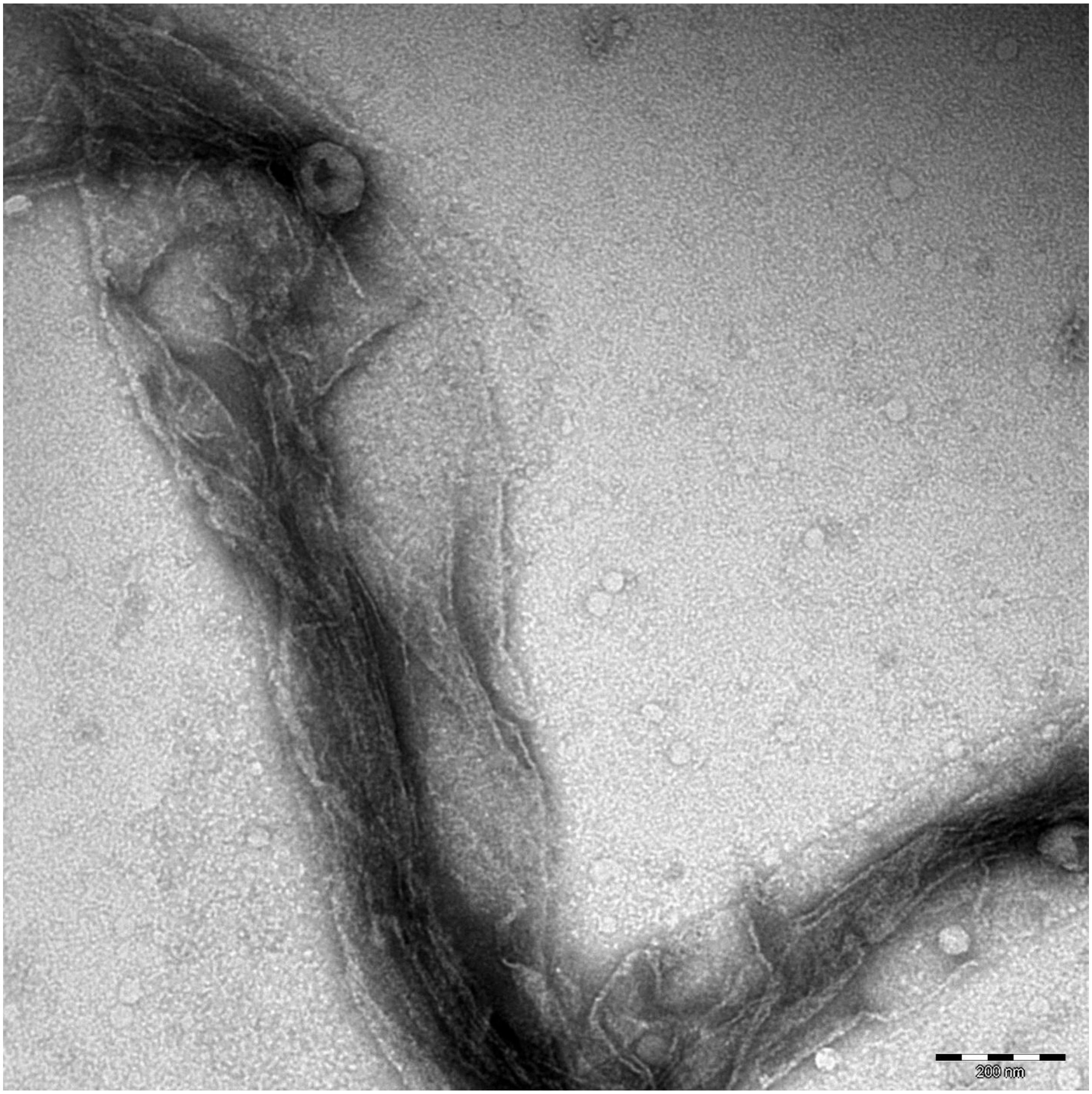
200 nm

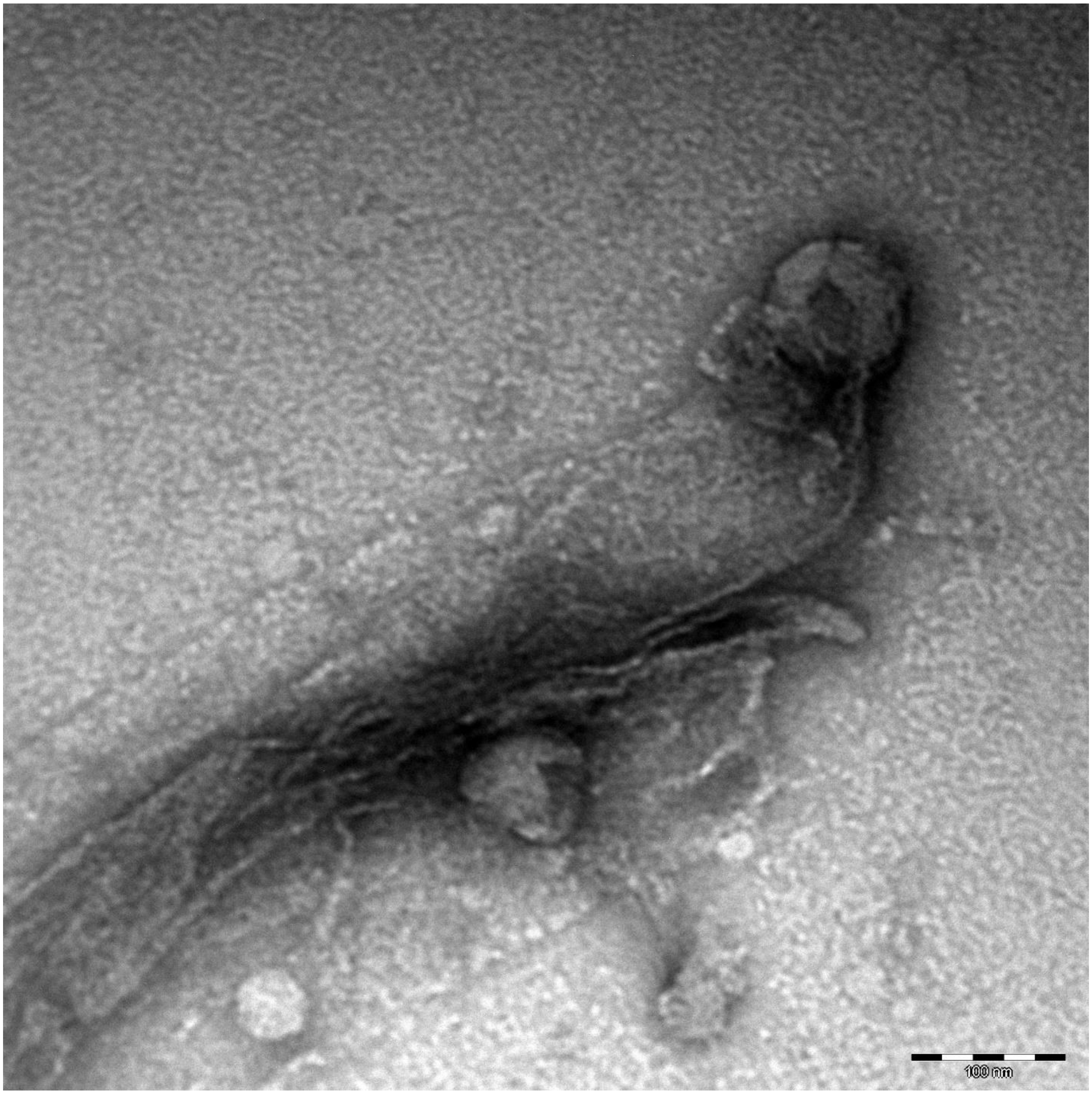
100 nm

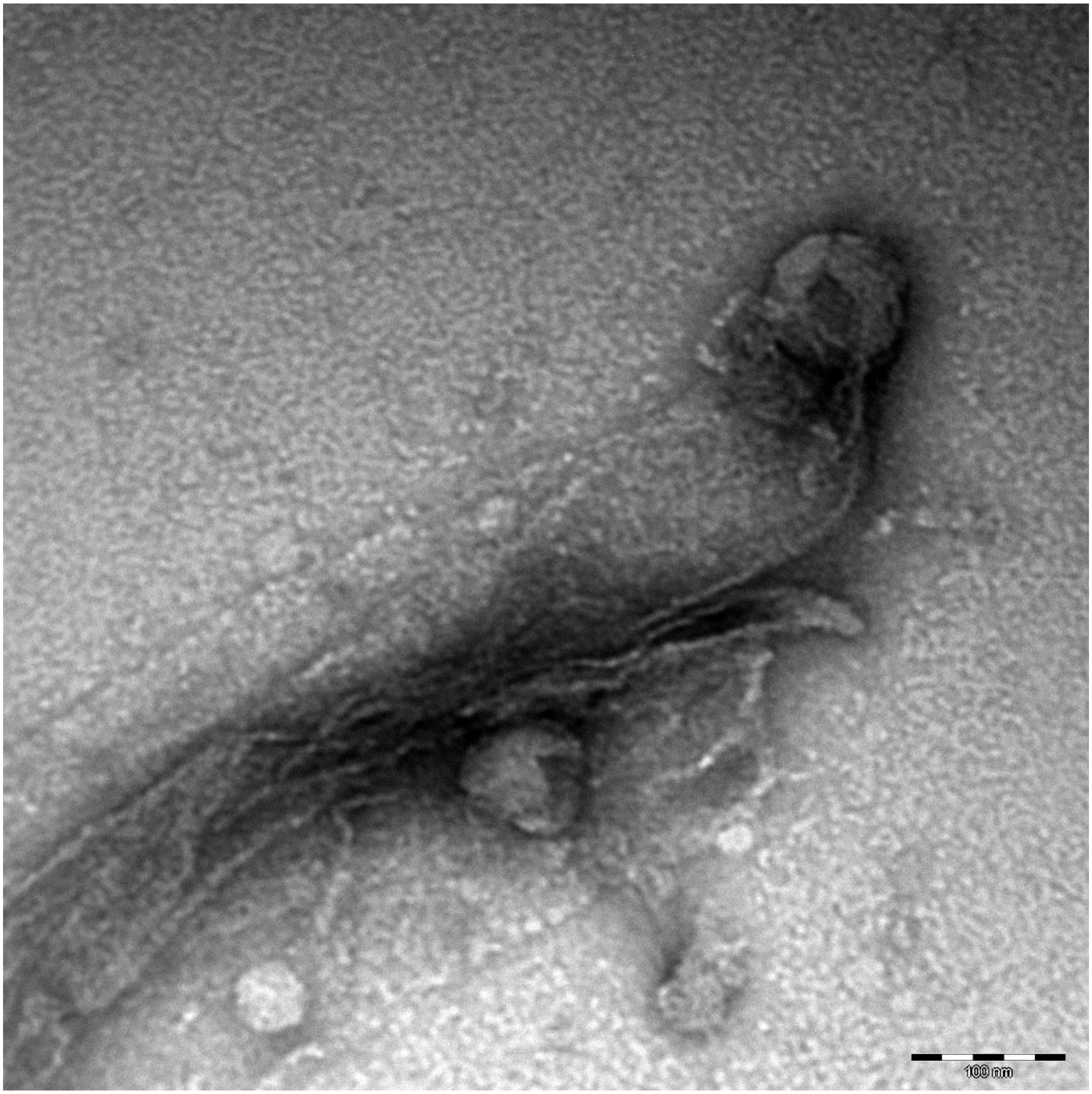
100 nm

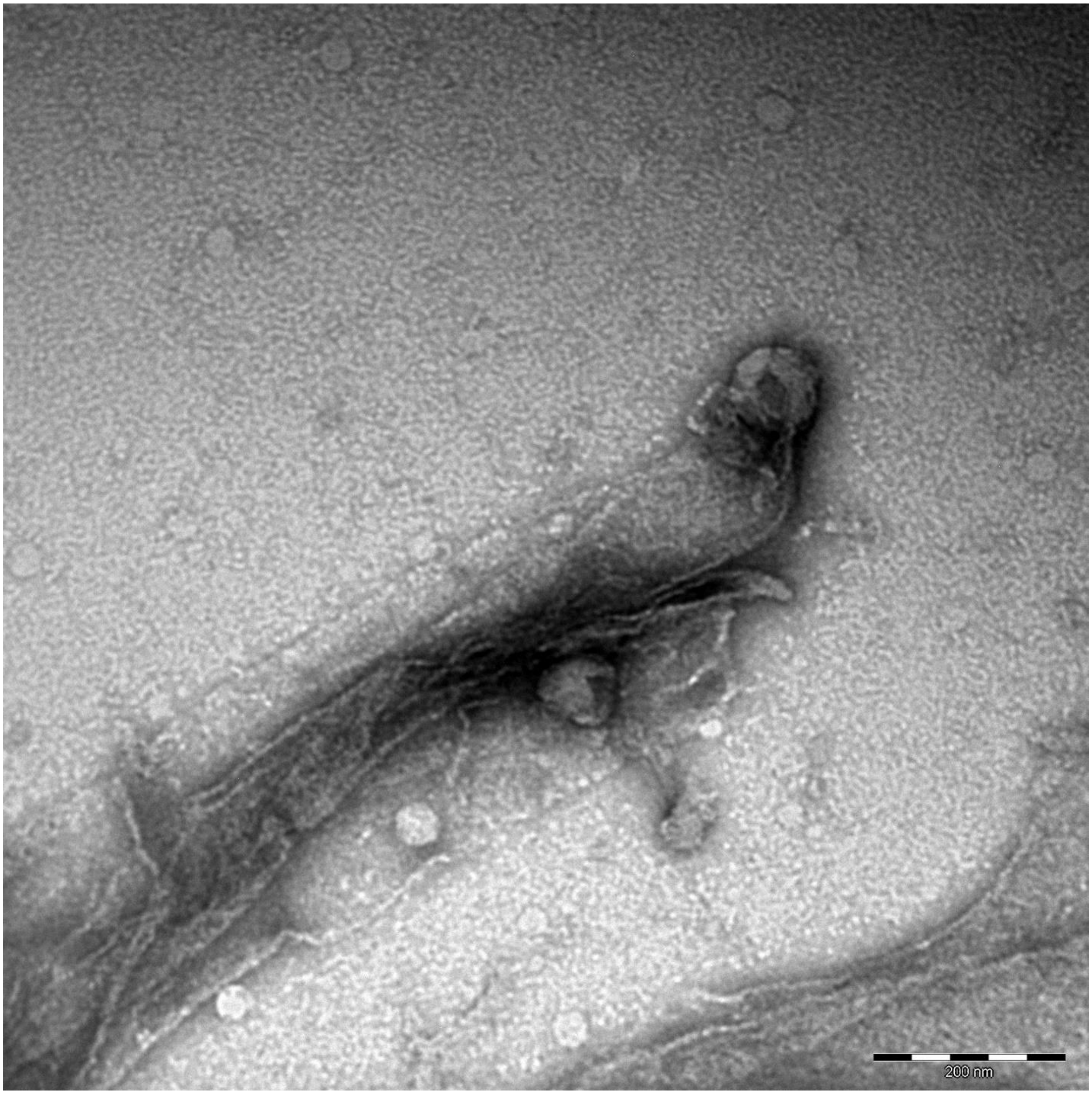
200 nm

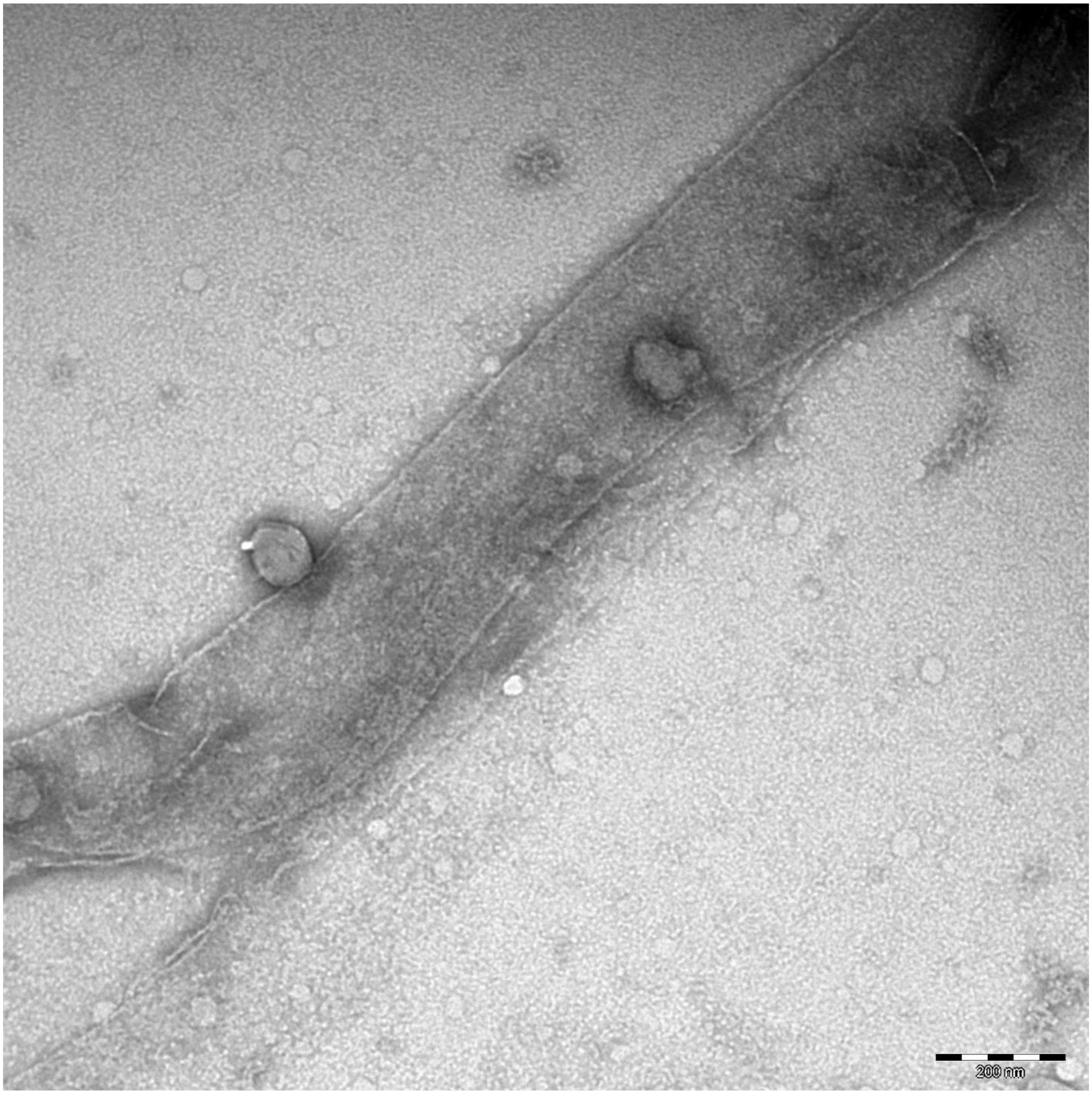
200 nm

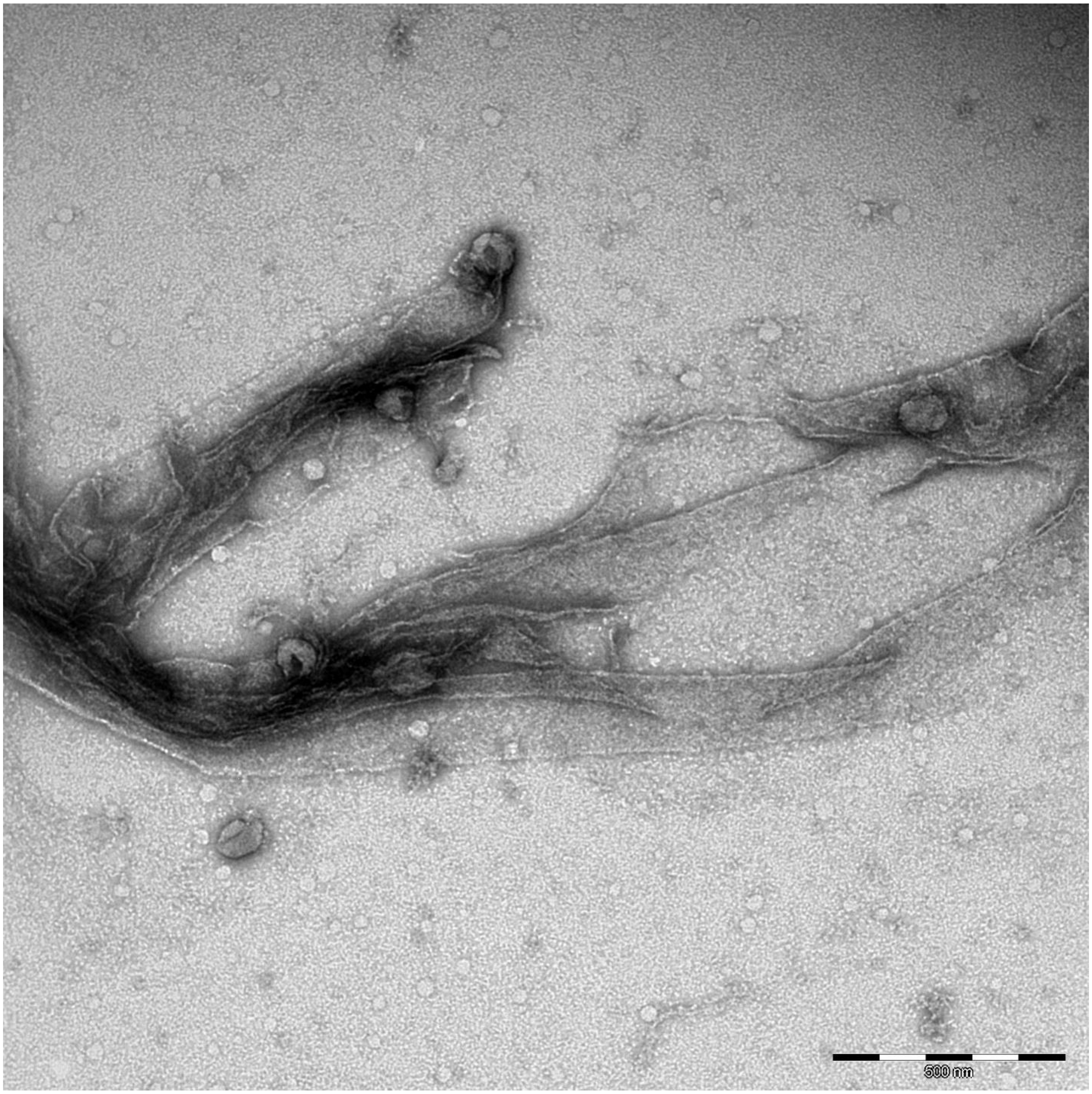
500 nm

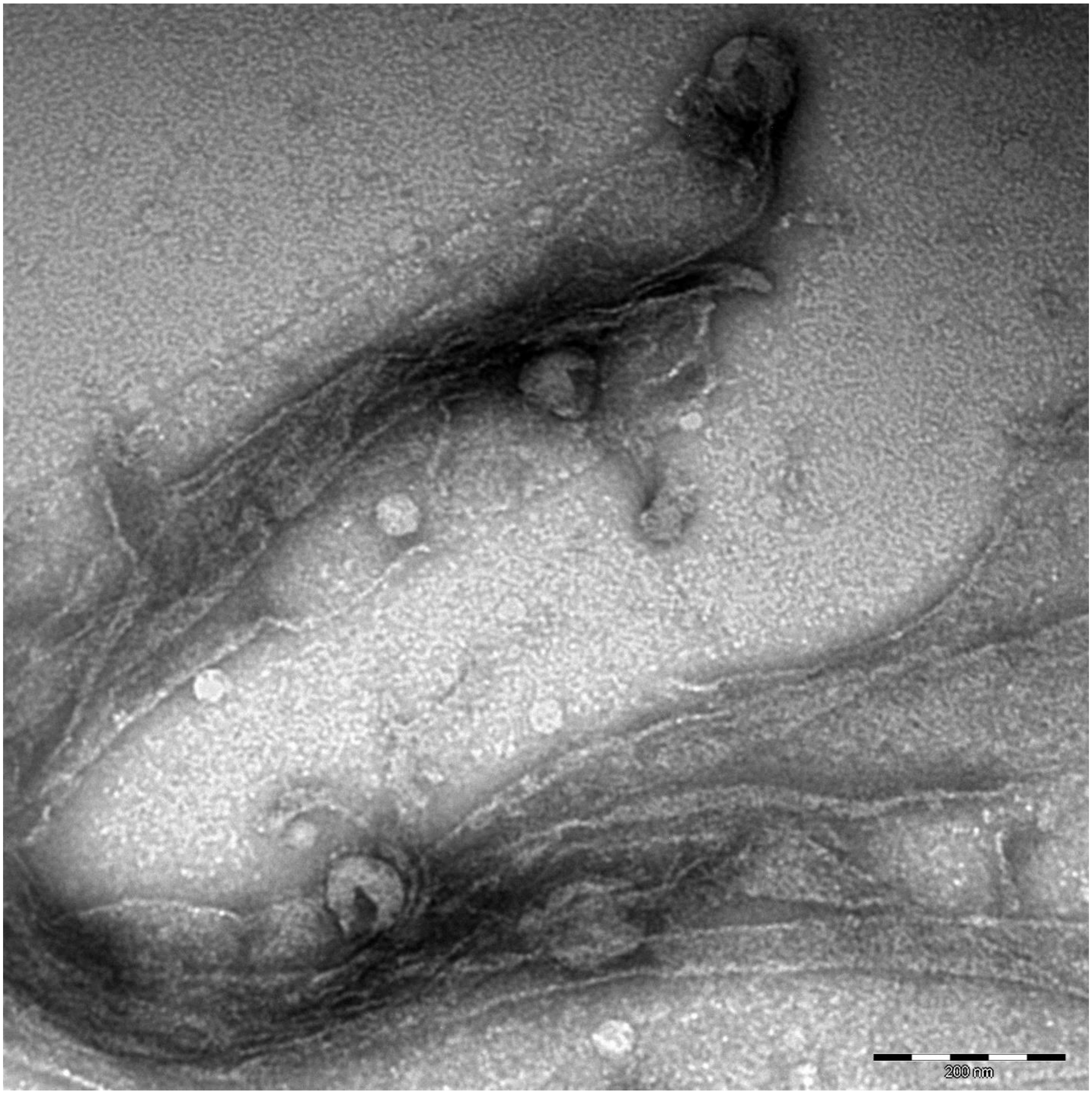
200 nm

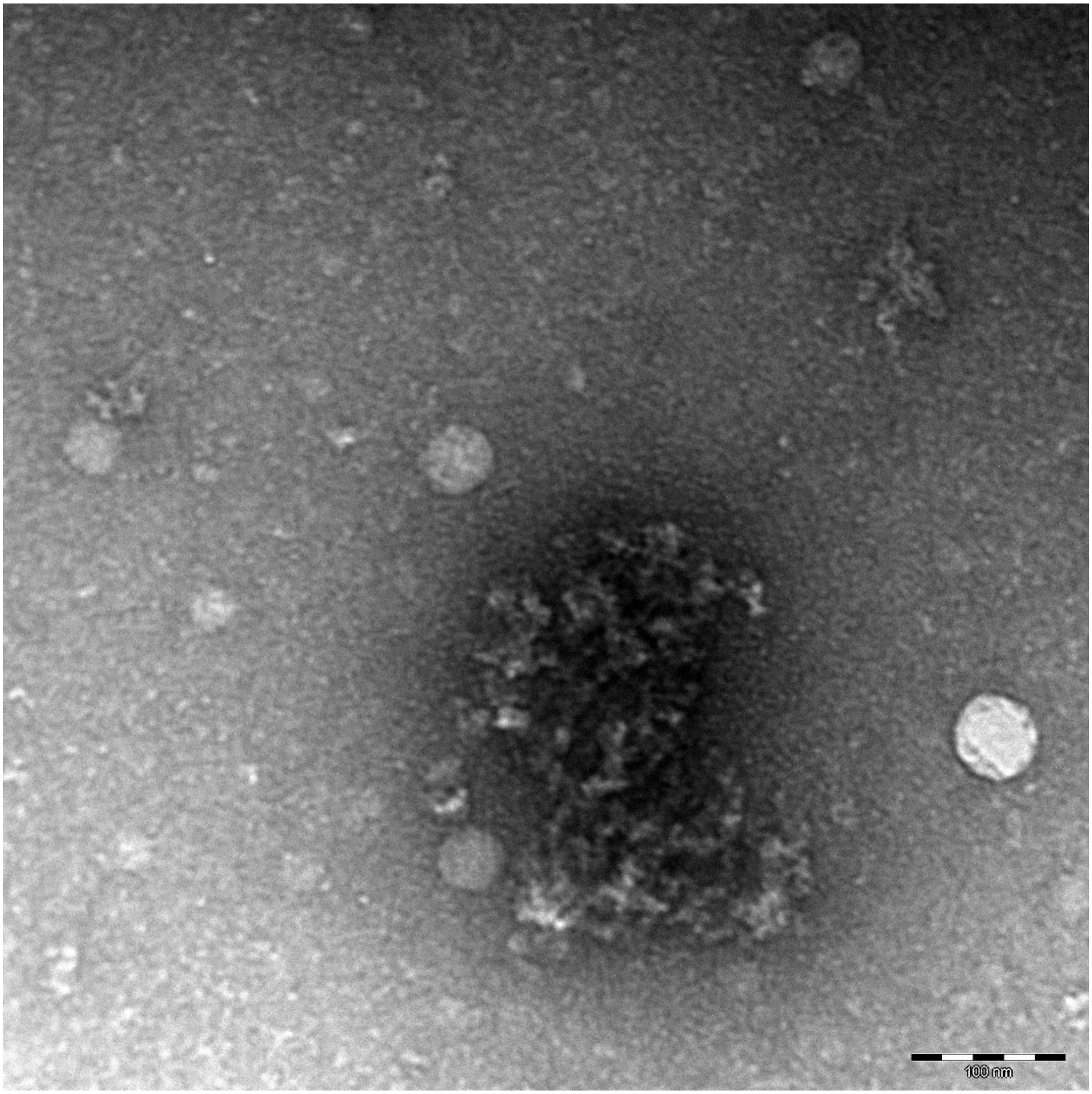
100 nm

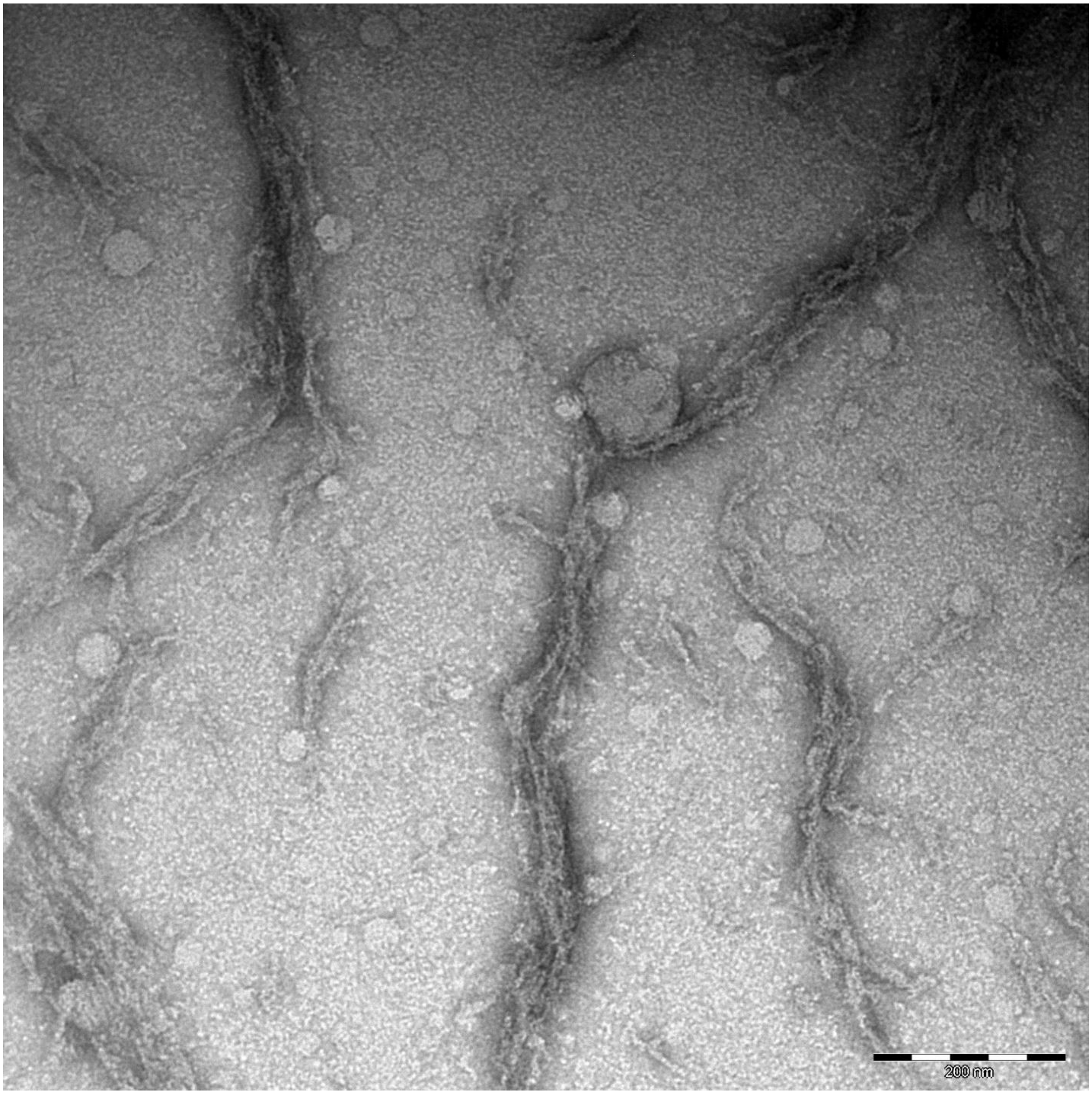
200 nm

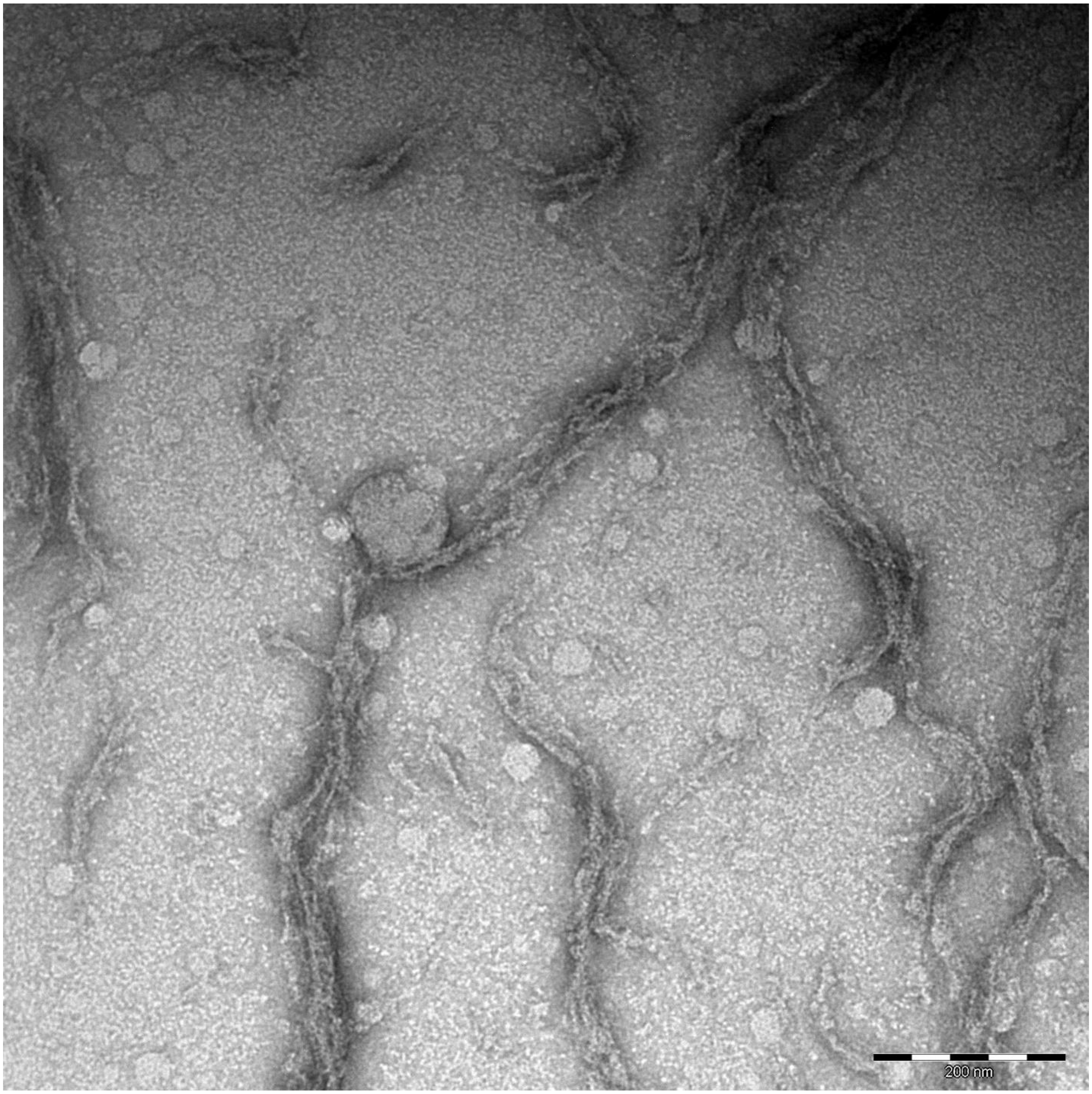
200 nm

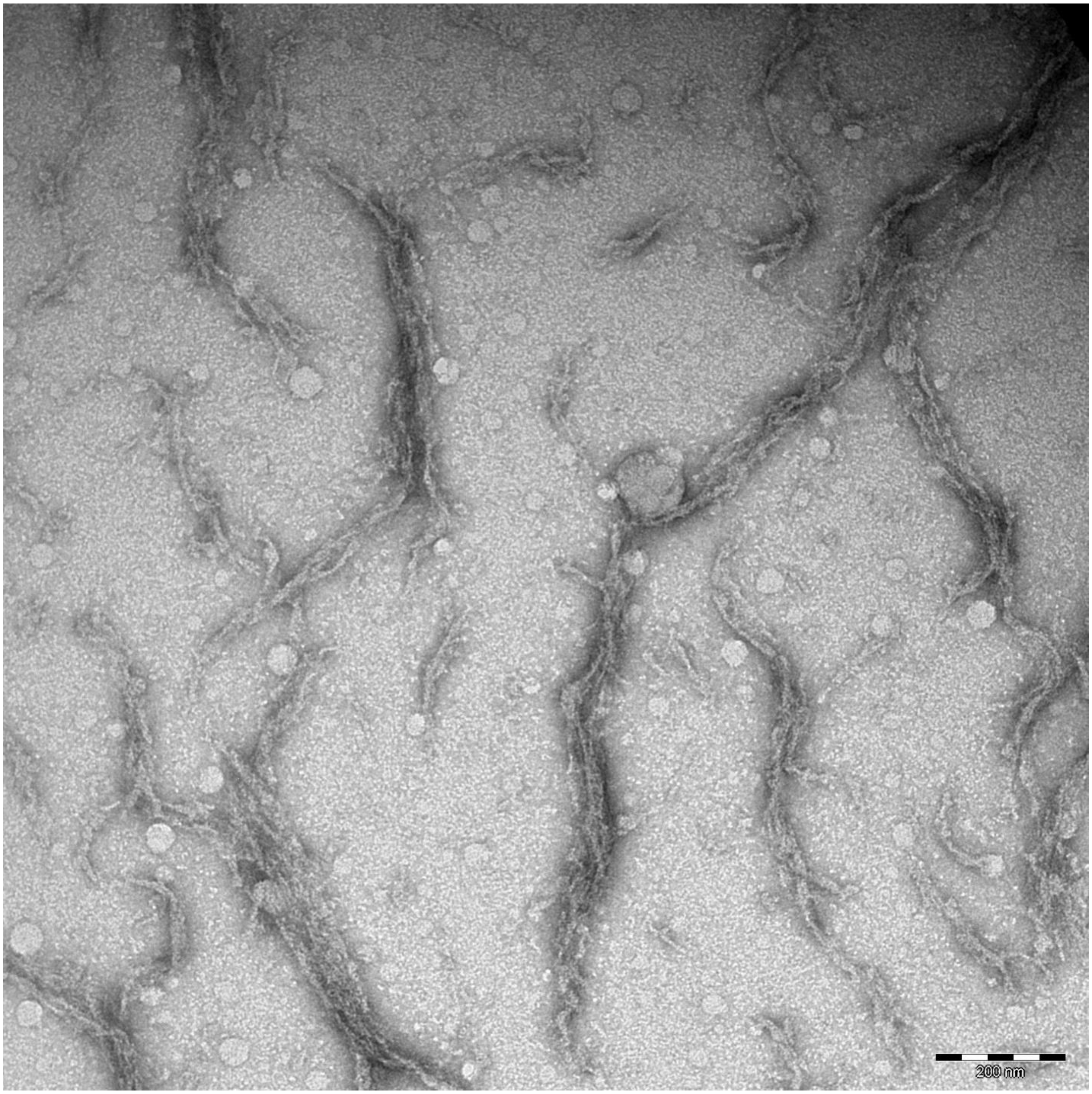
200 nm

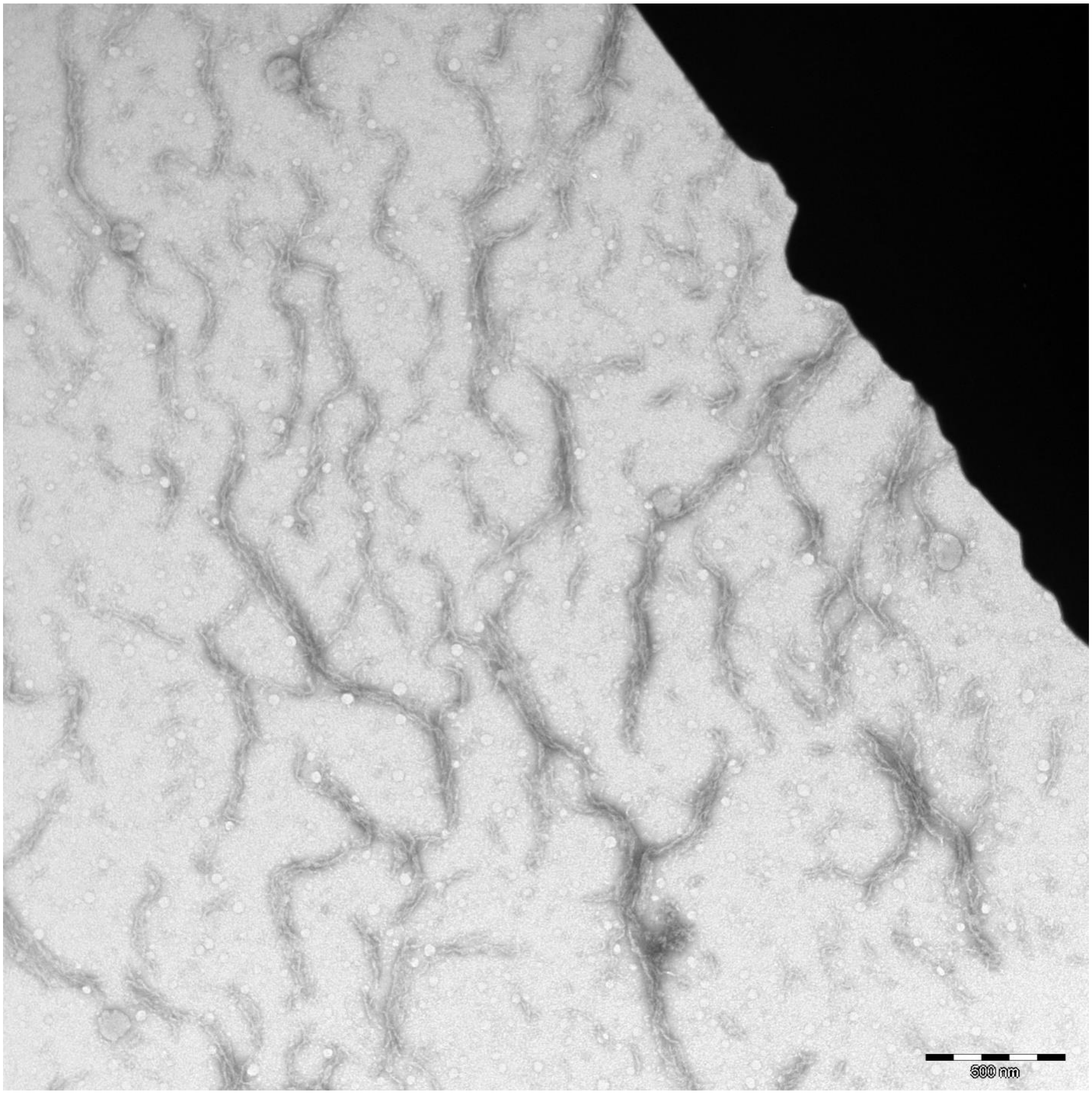
500 nm

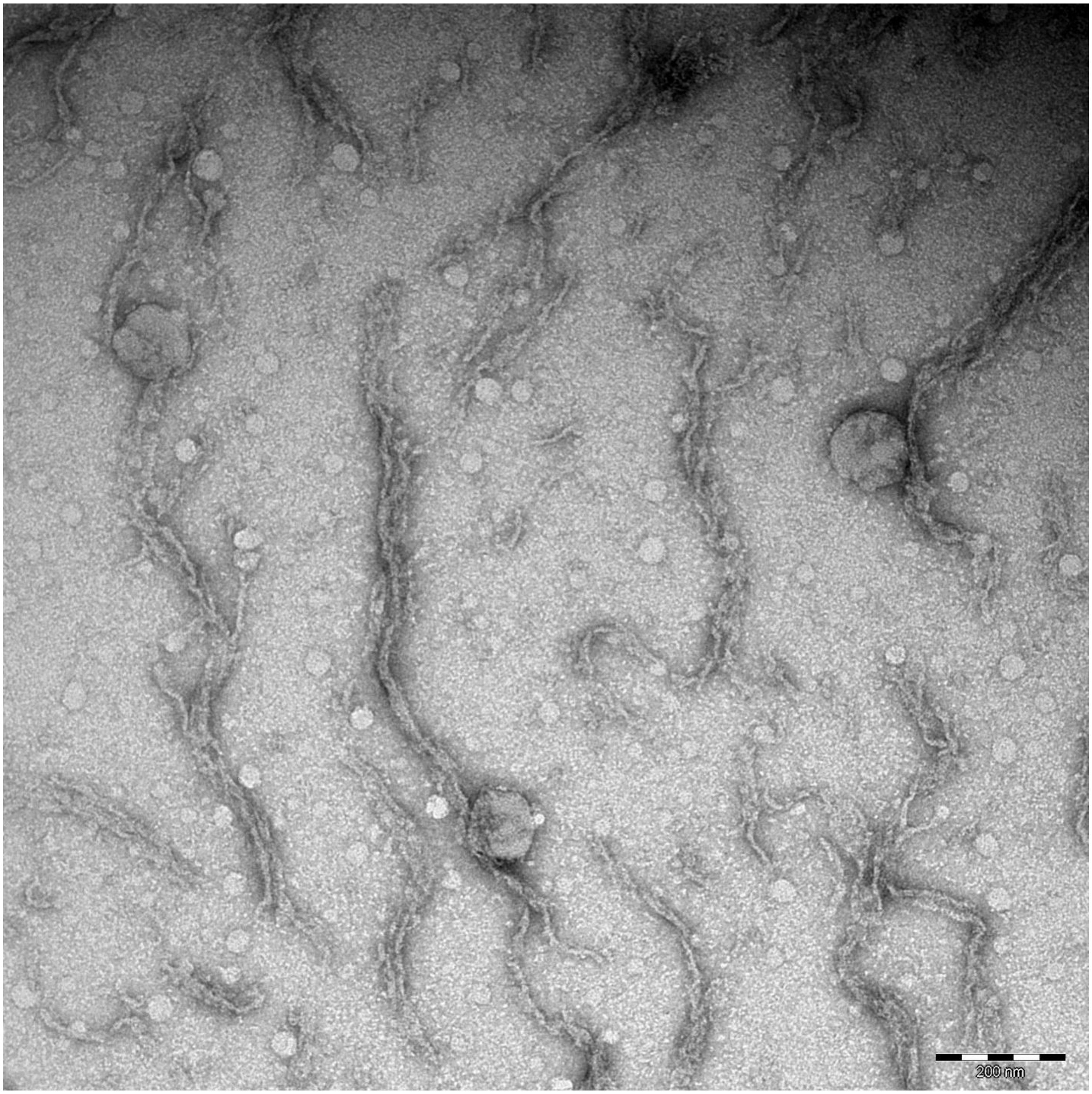
200 nm

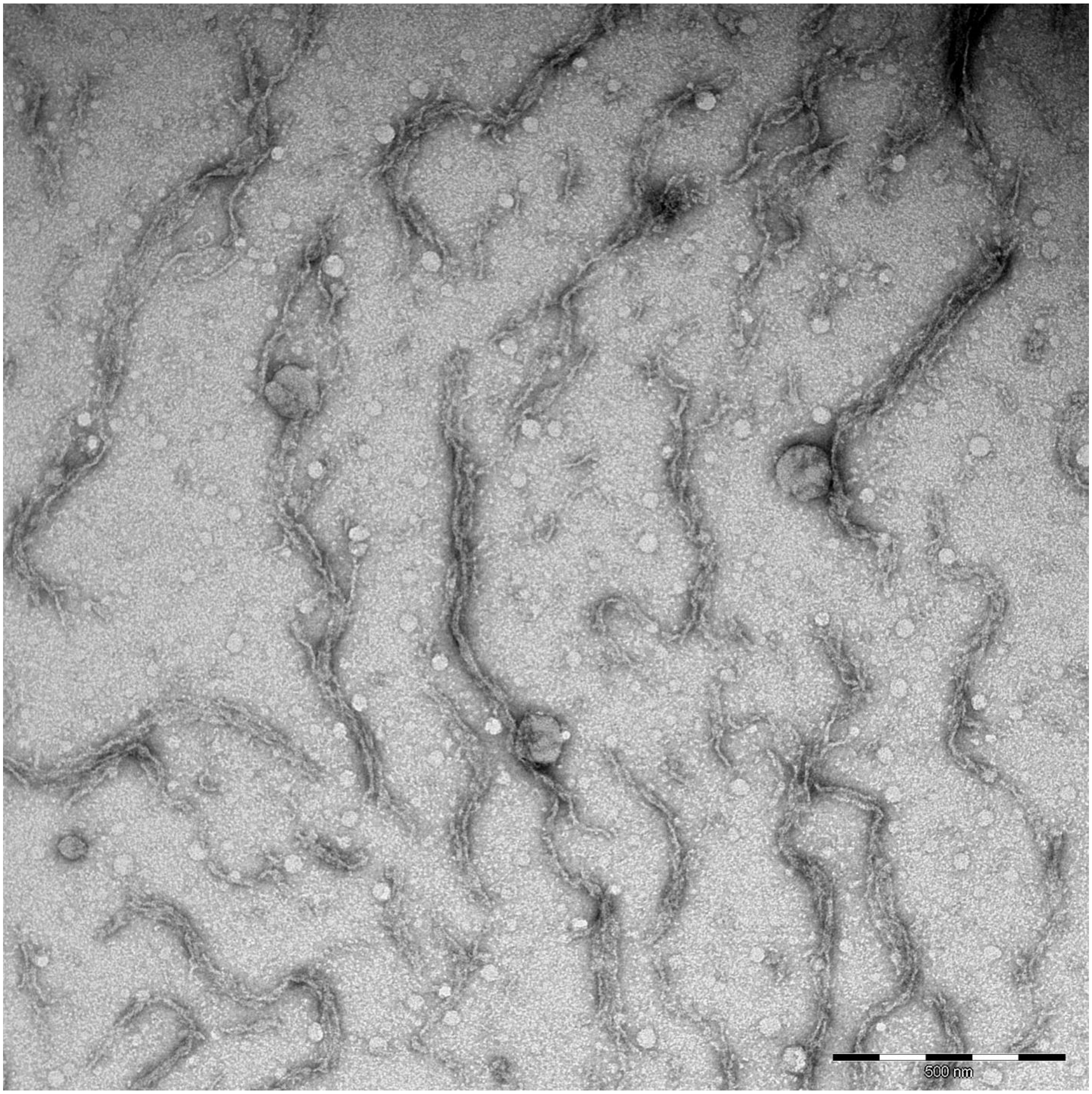
500 nm

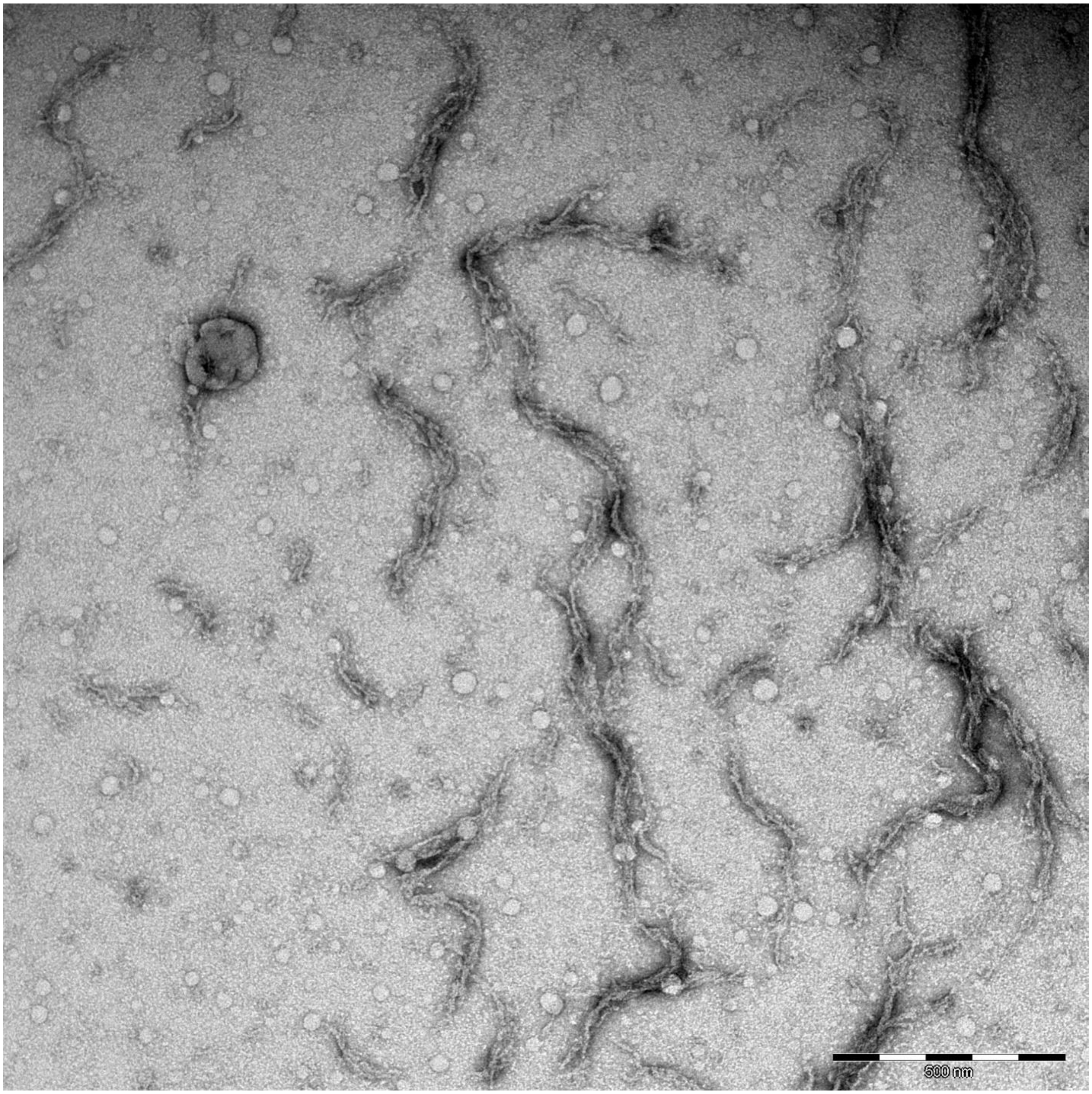
500 nm

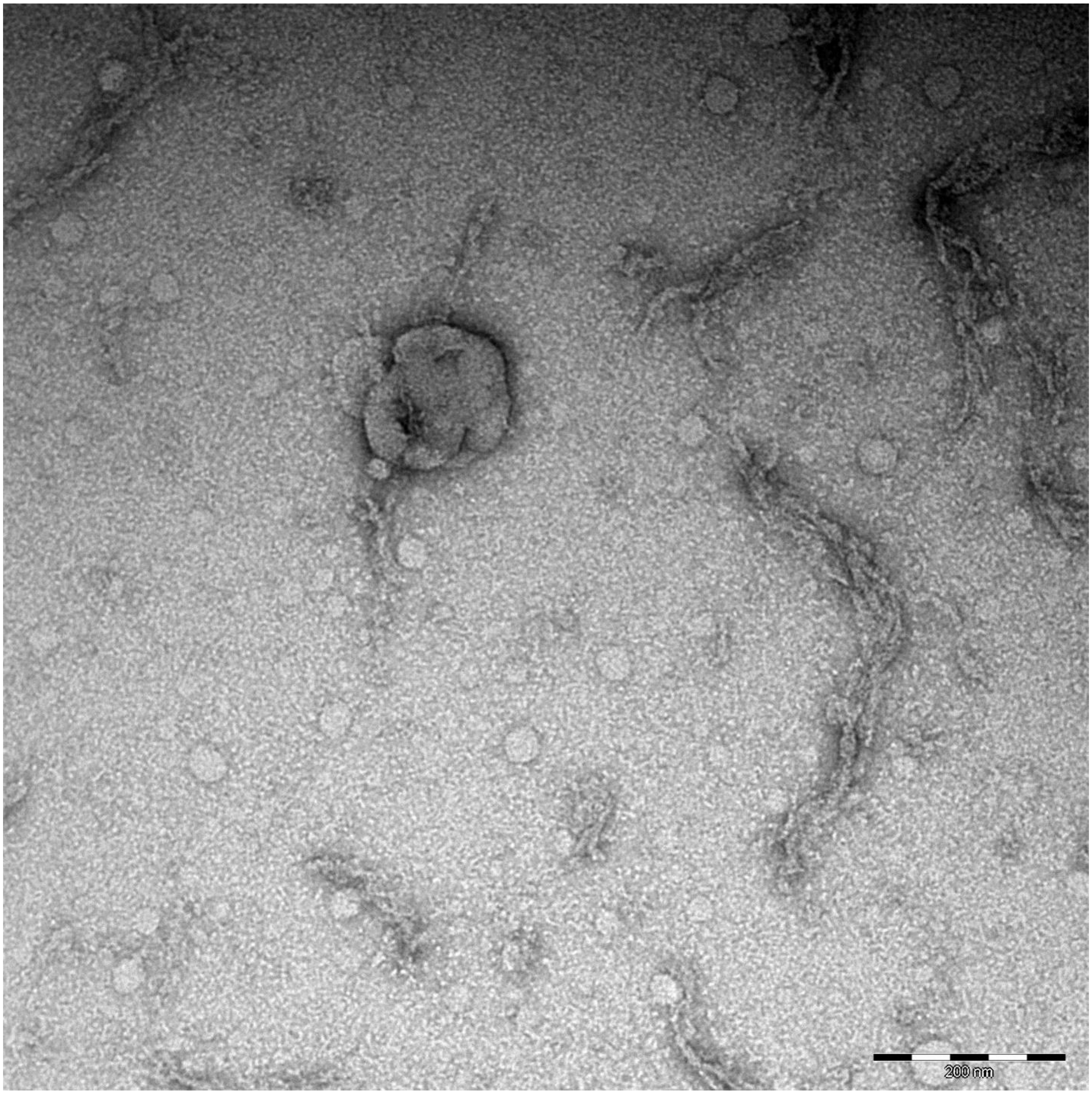
200 nm

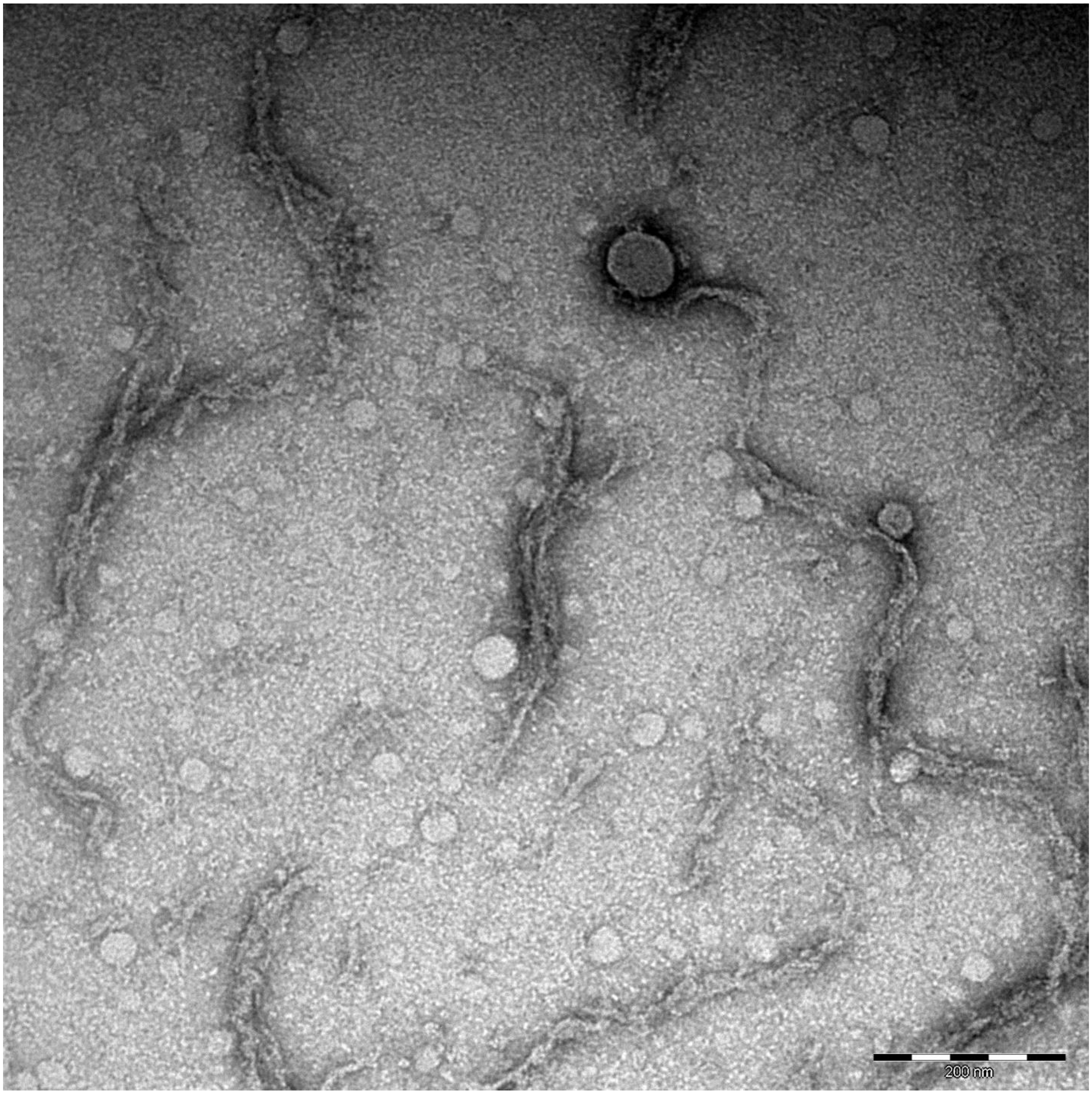
200 nm

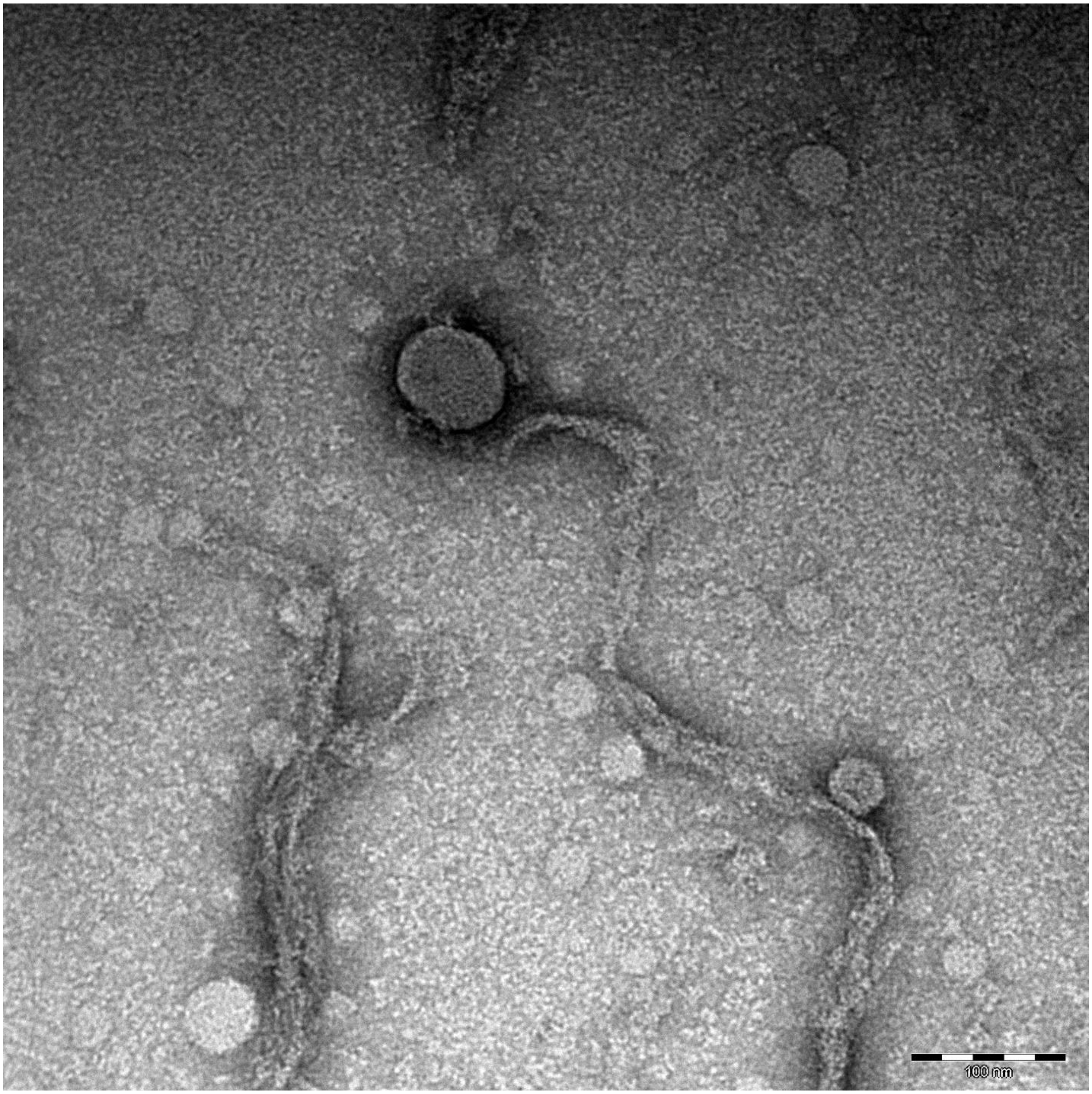
100 nm

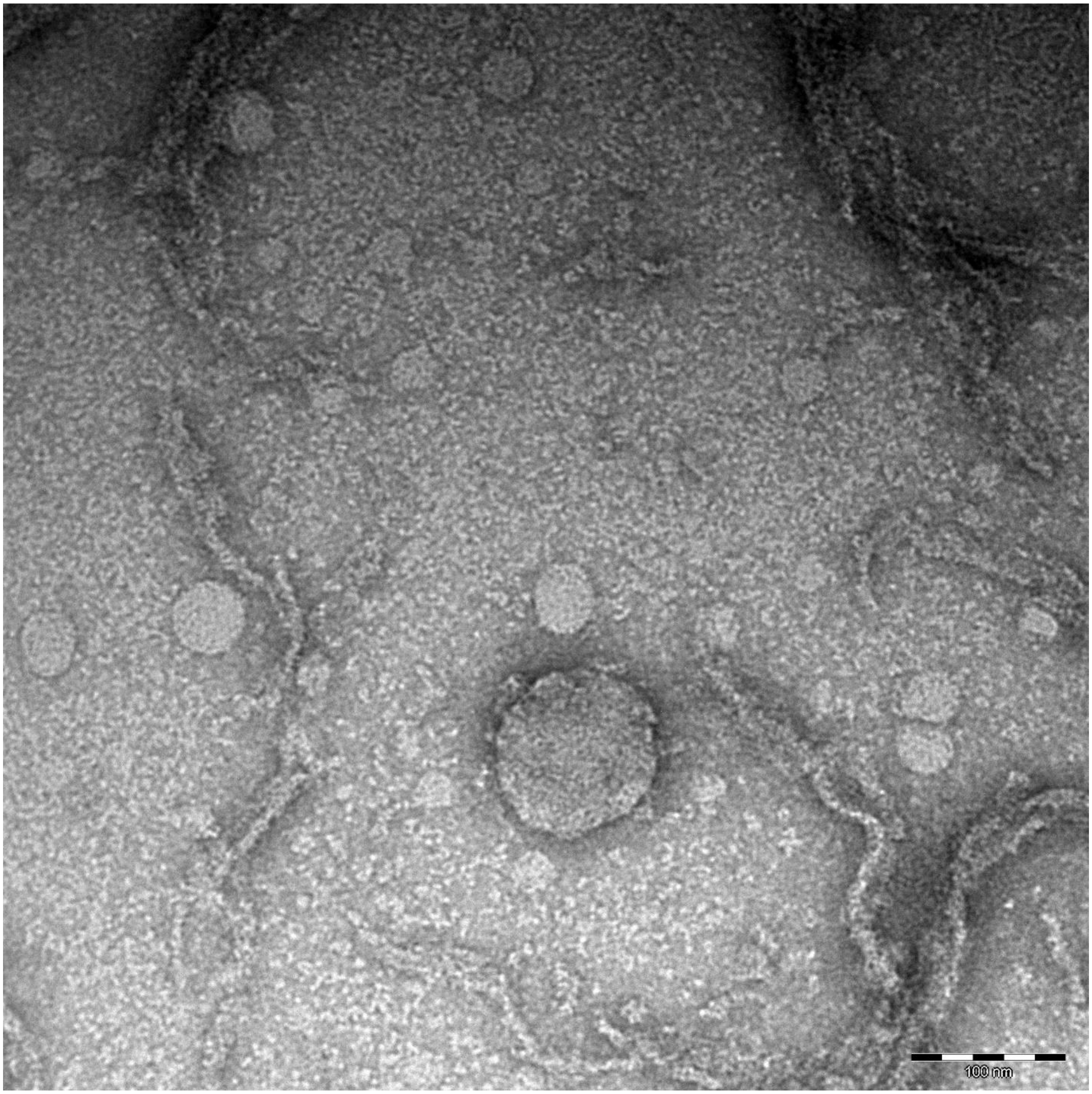
100 nm

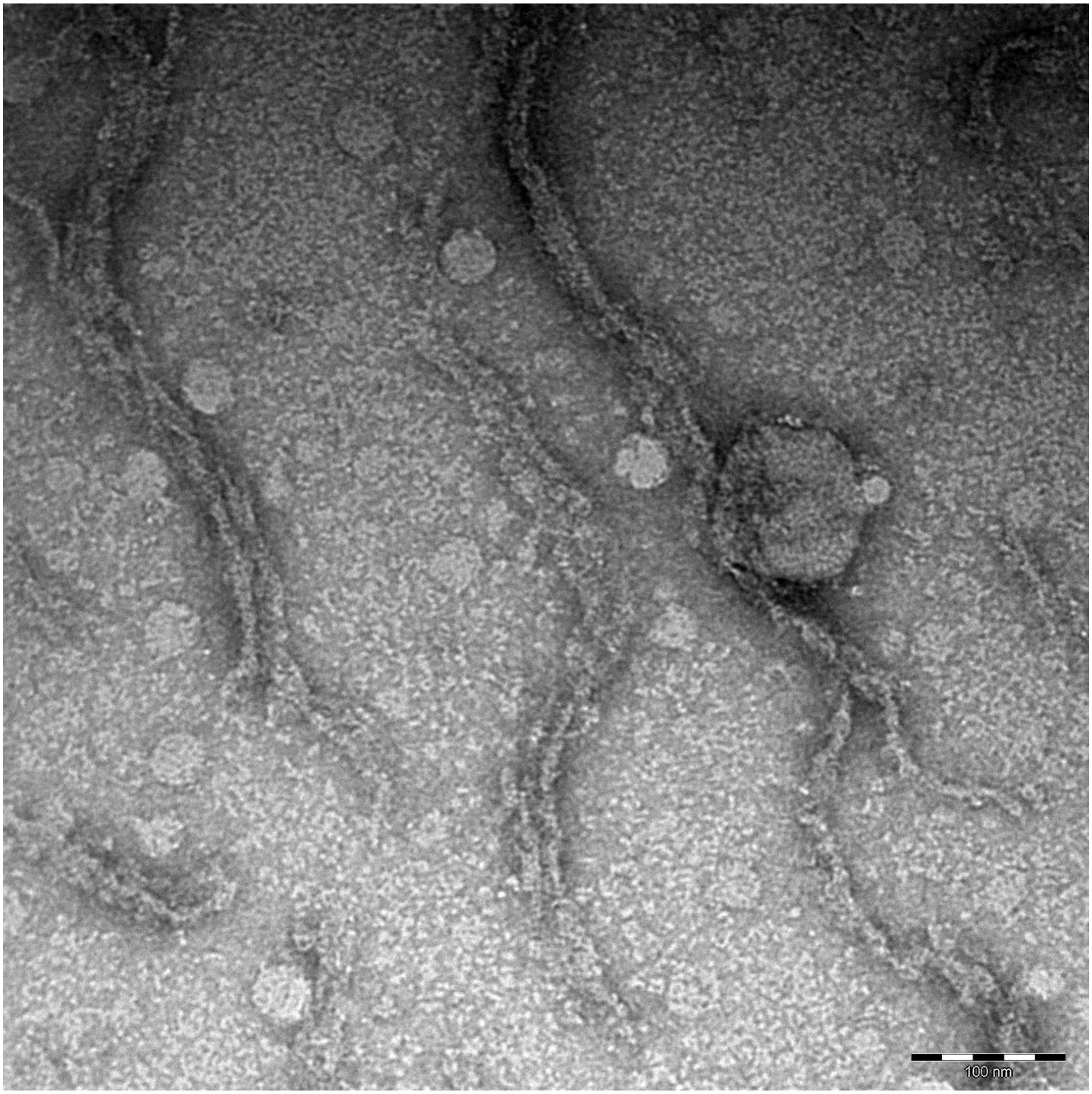
100 nm
