## Supplementary Figure 4 for "SARS-CoV-2 and HSV-1 Induce Amyloid Aggregation in Human CSF Resulting in Drastic Soluble Protein Depletion"

**Supplementary Figure 4. Amyloid fibrils formed upon HSV-1 incubation with human CSF.** Negatively stained TEM images of HSV-1 incubated with CSF for 48 h at 37 °C. Raw pictures.

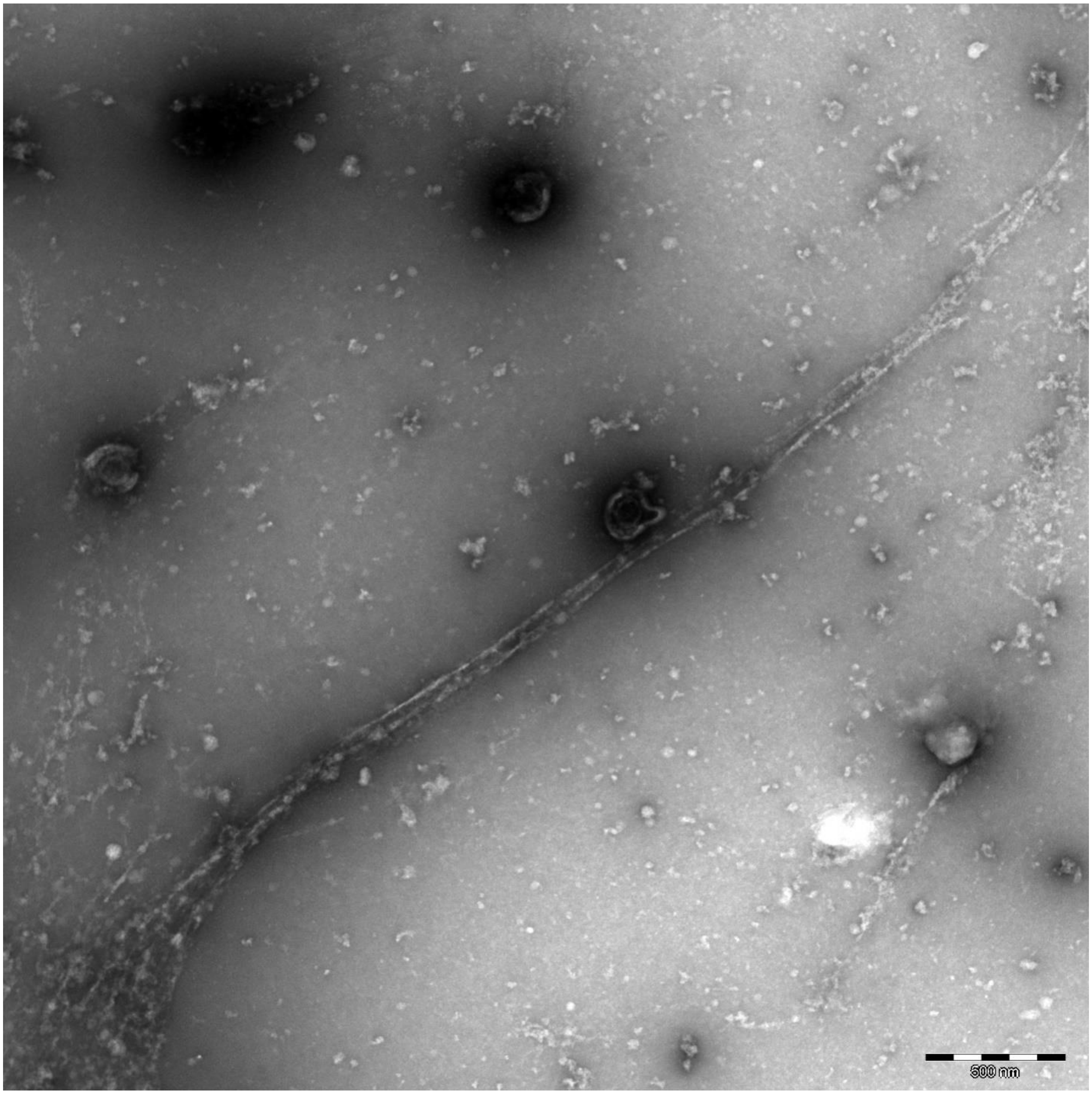

500 nm

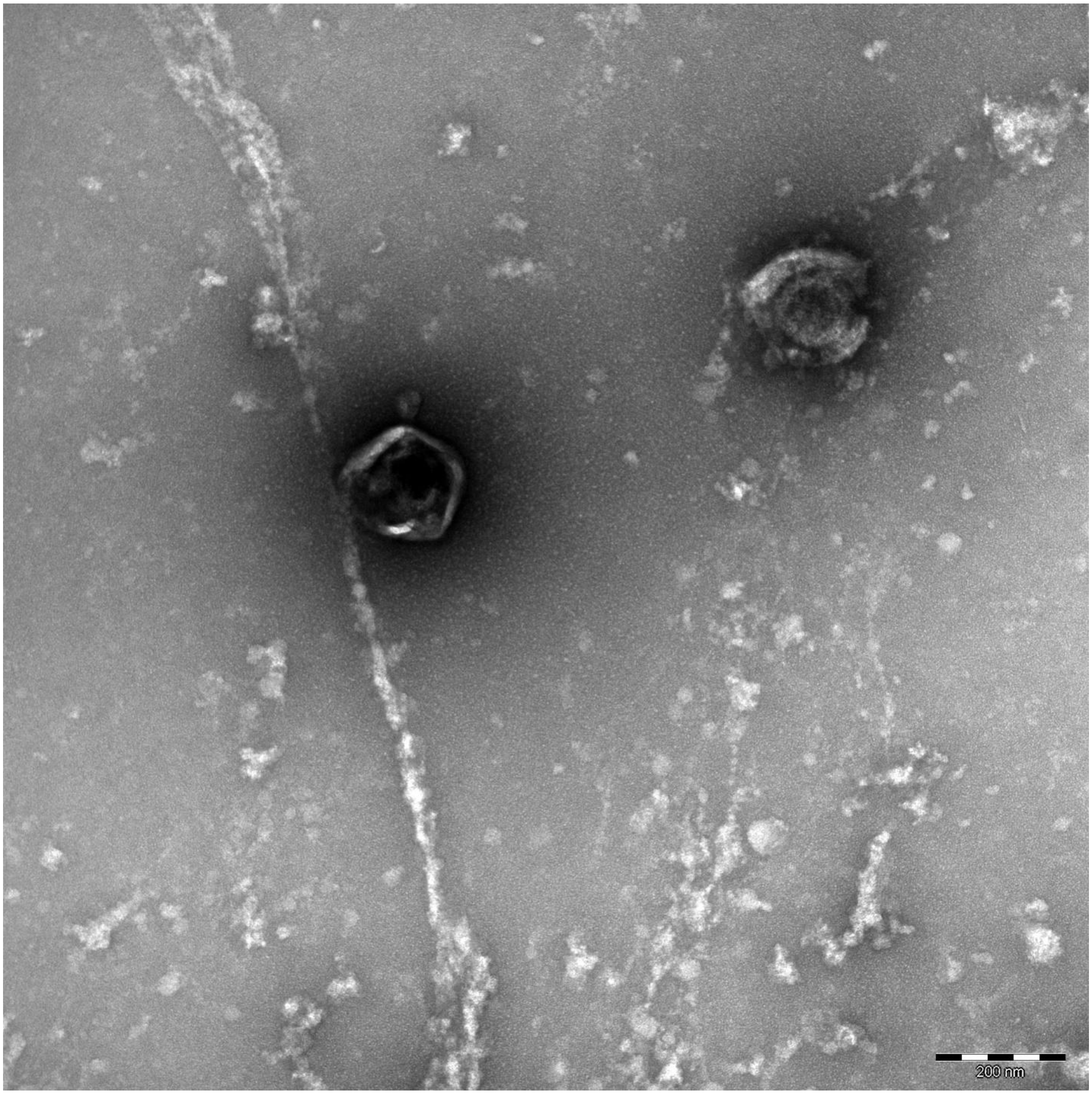
200 nm

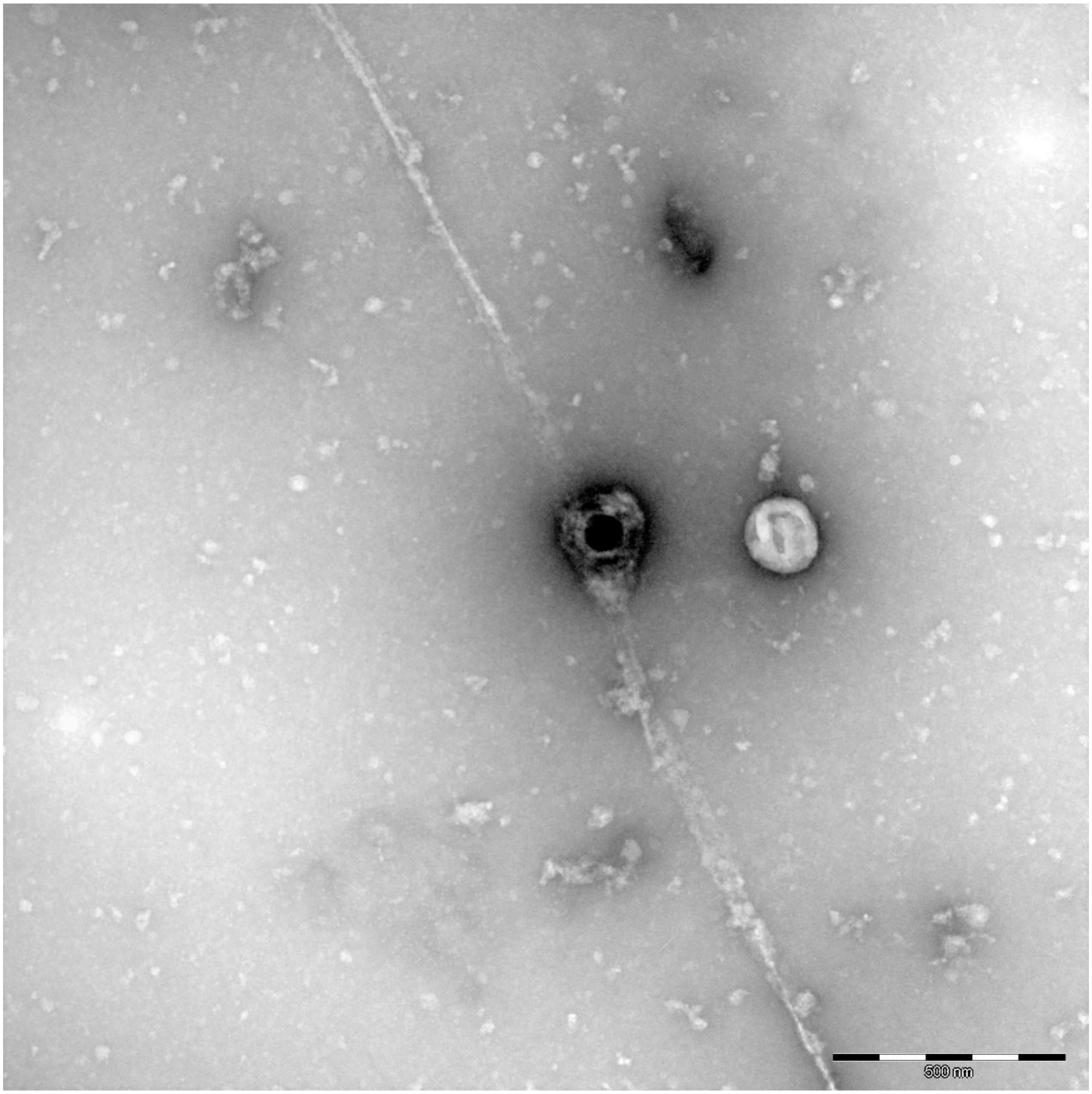
500 nm

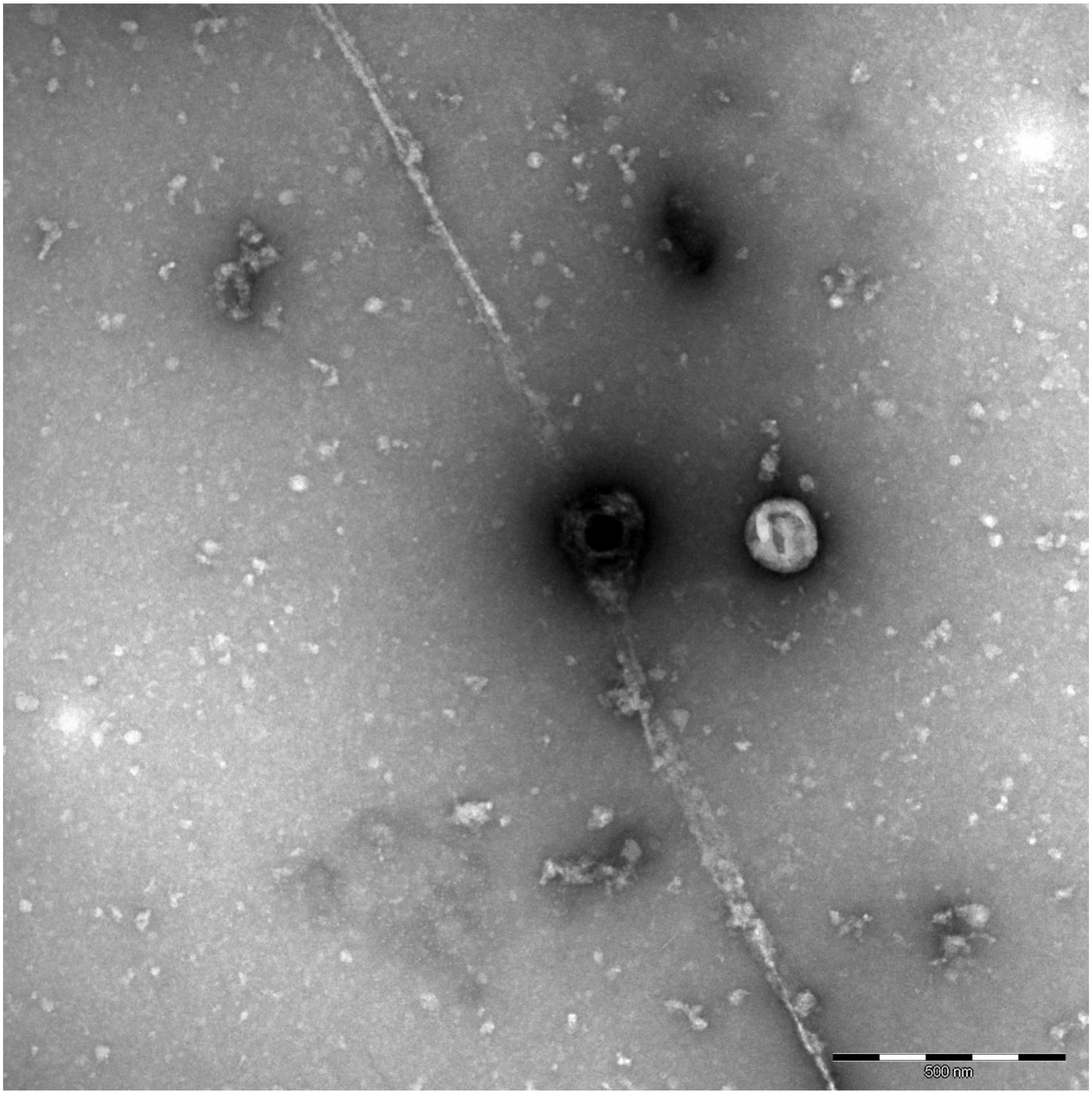
500 nm

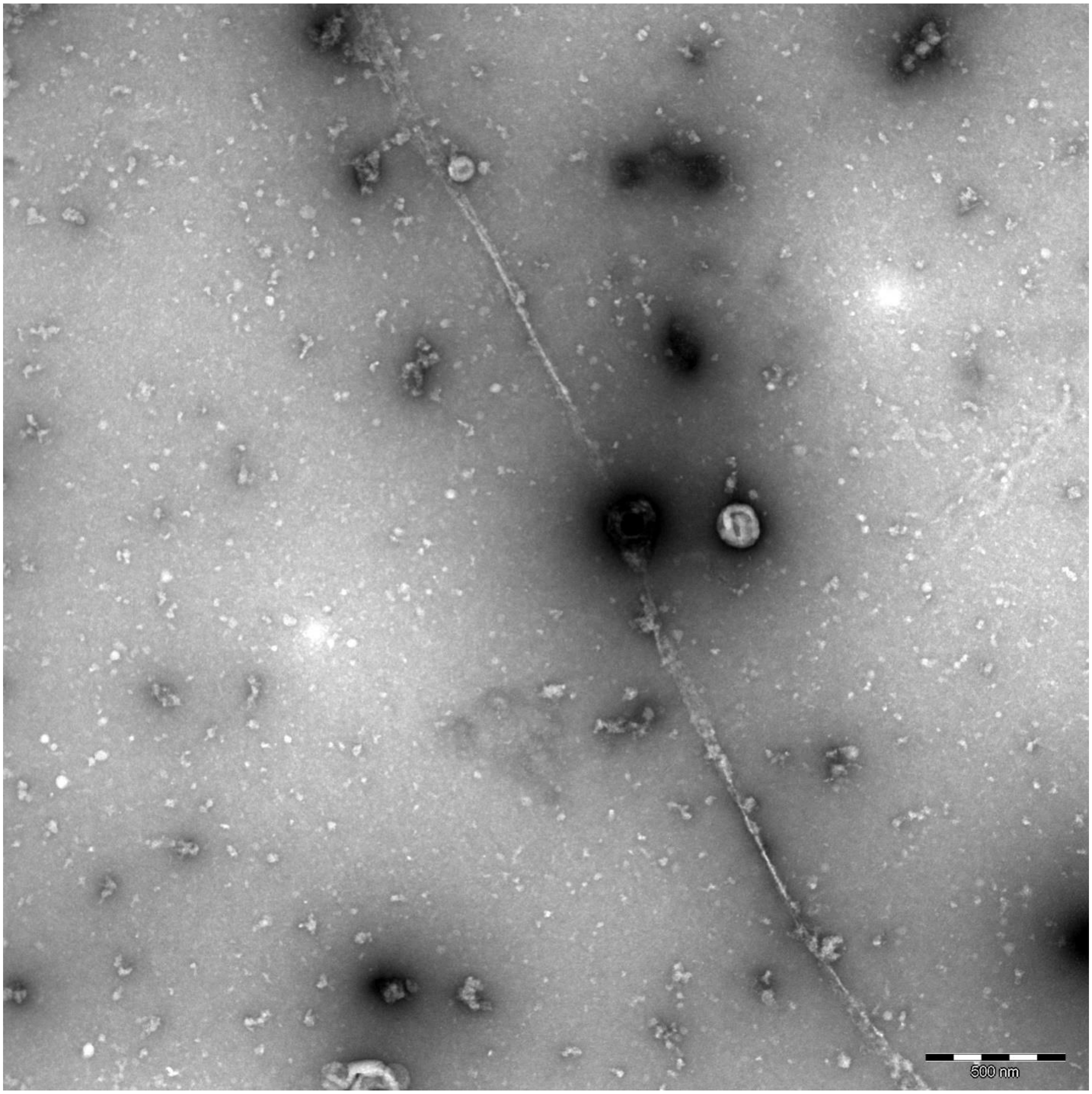
500 nm

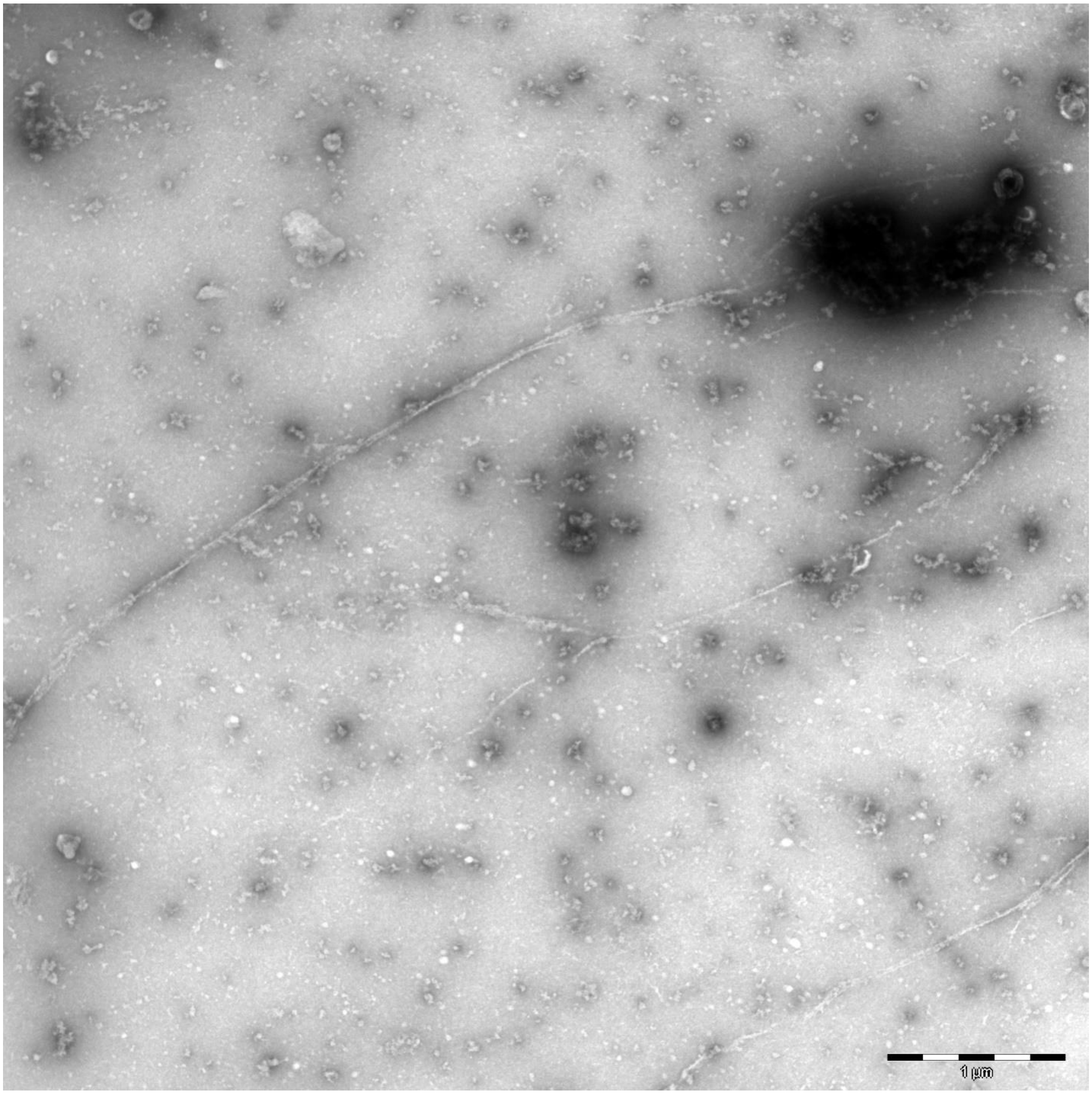
1 µm

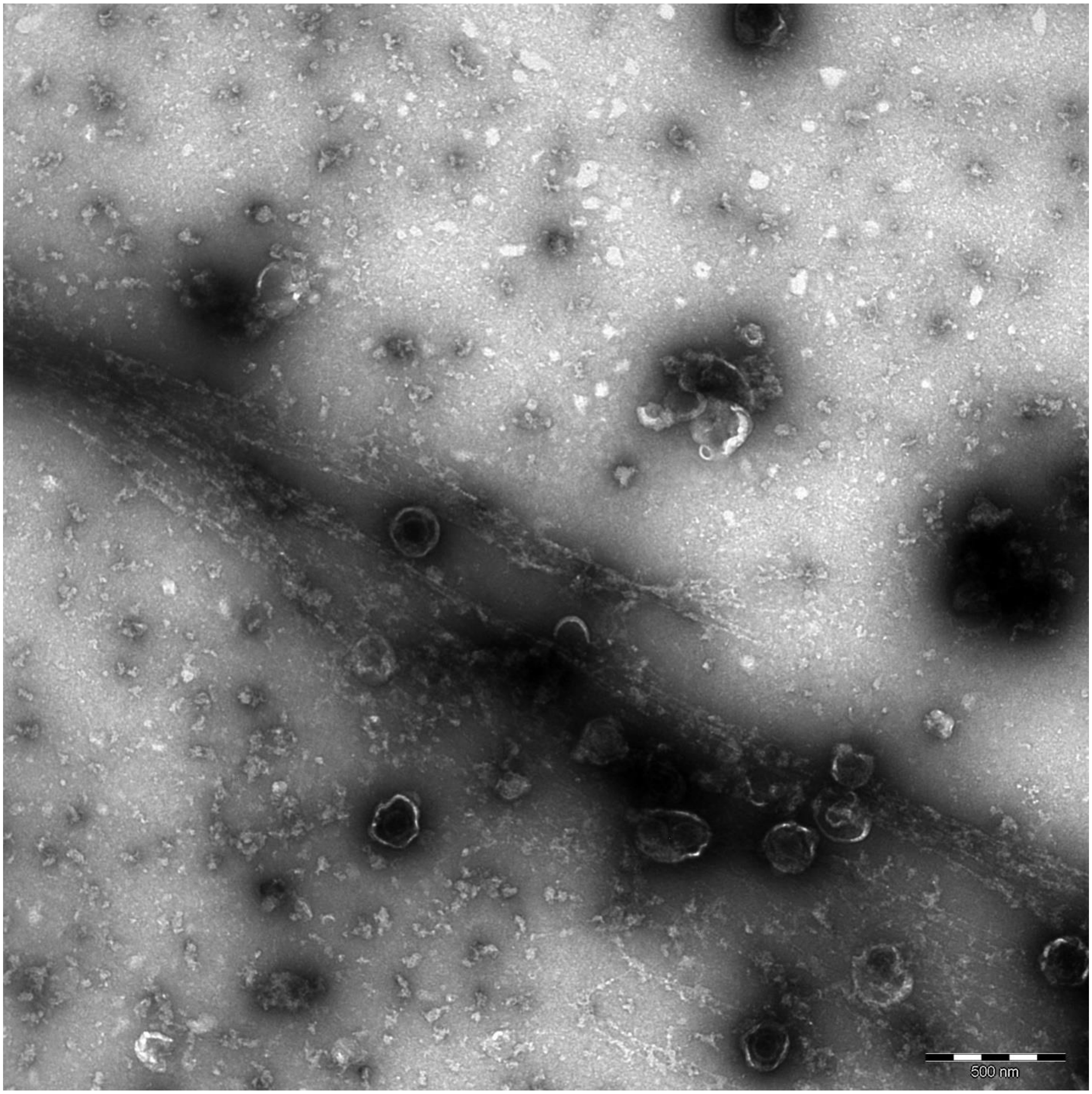
500 nm

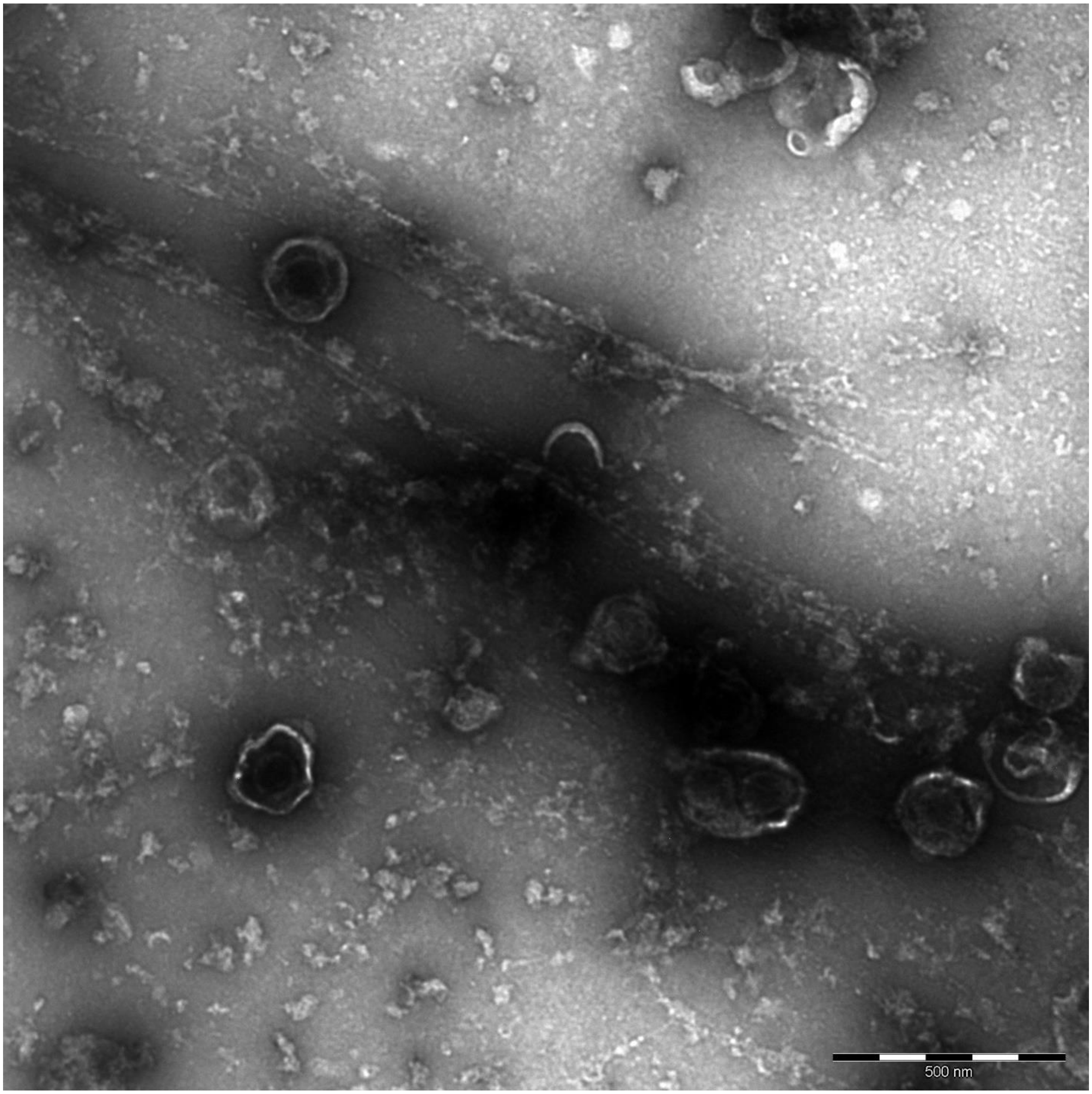
500 nm

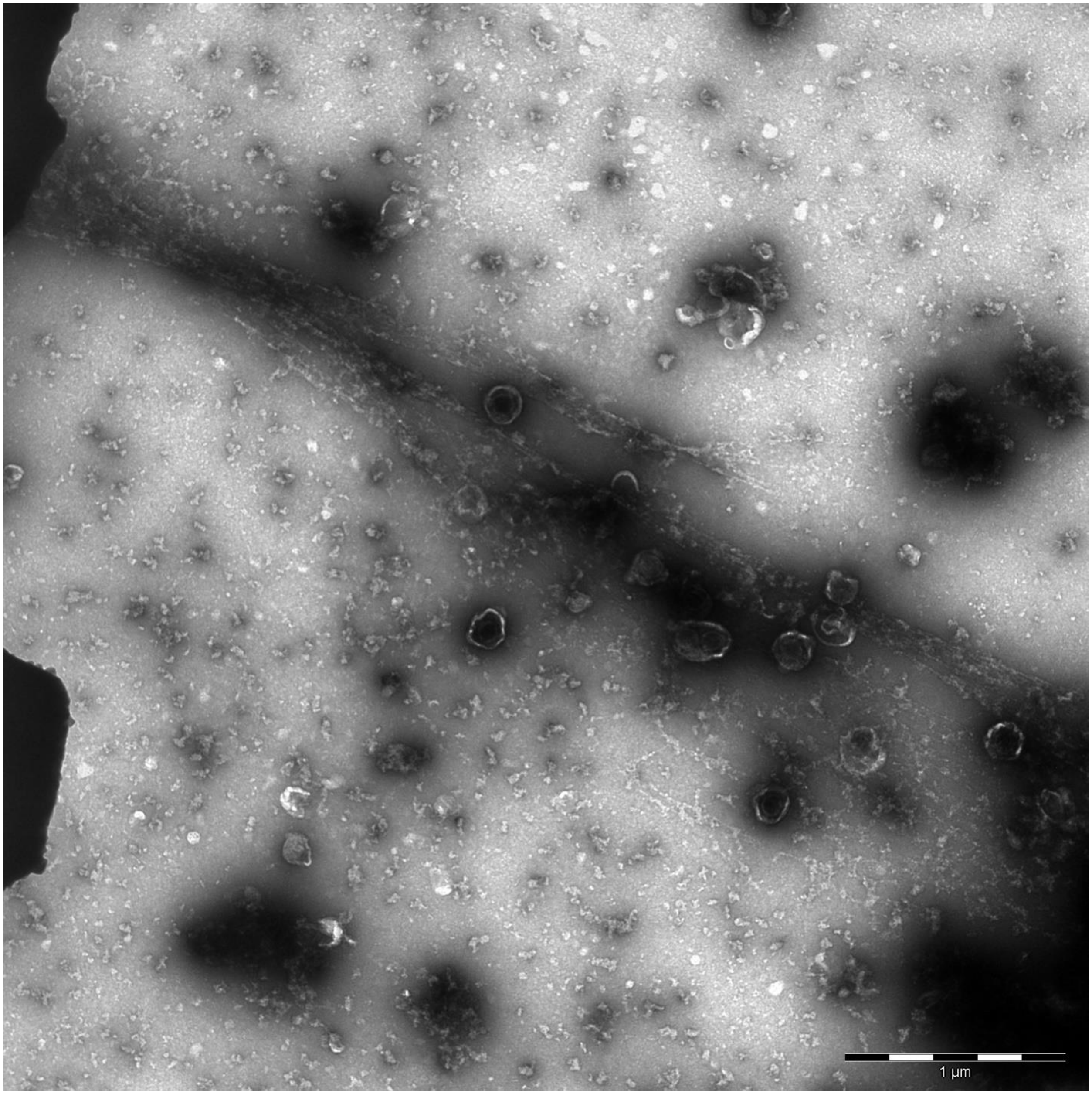
1 µm

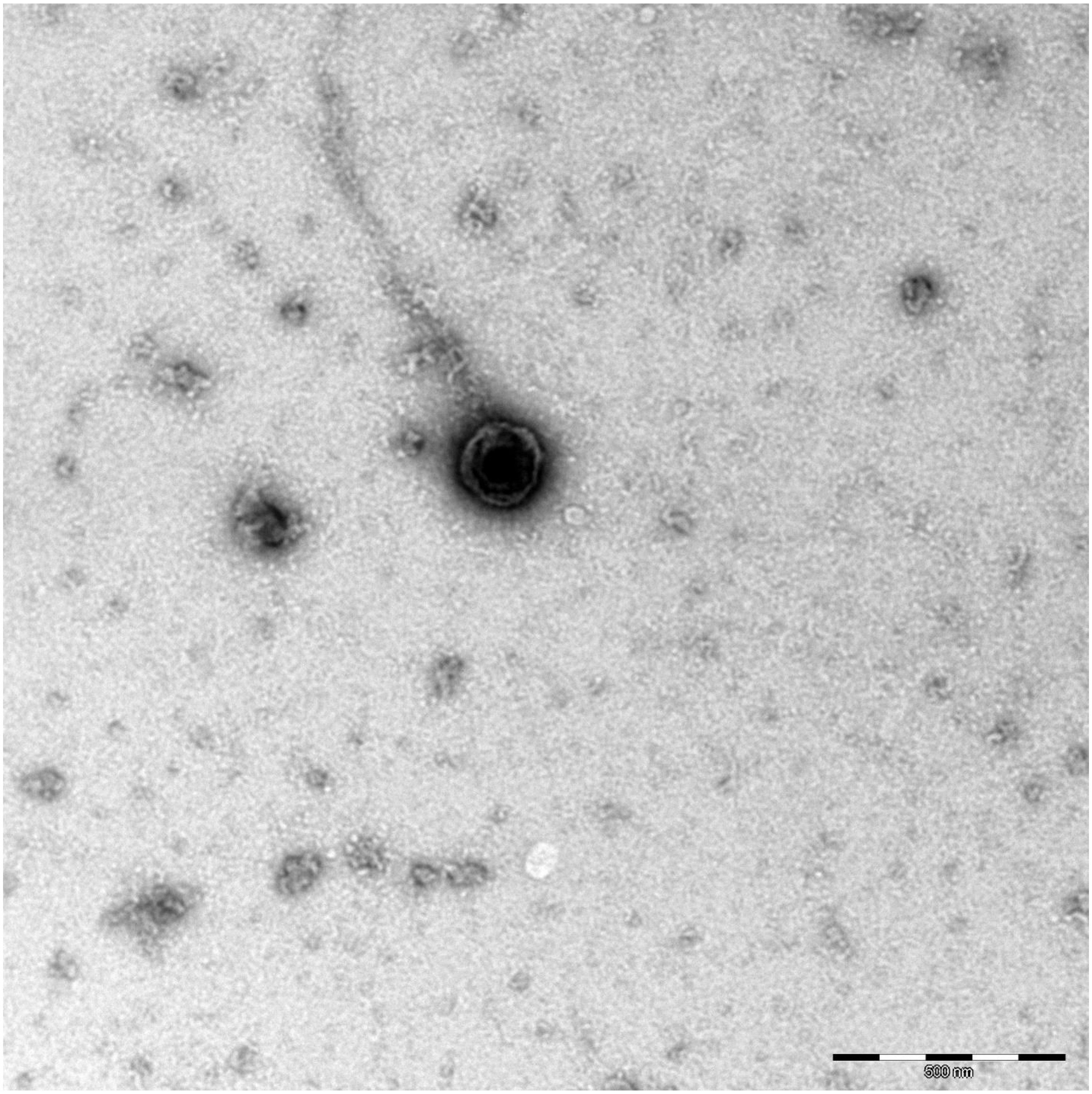
500 nm

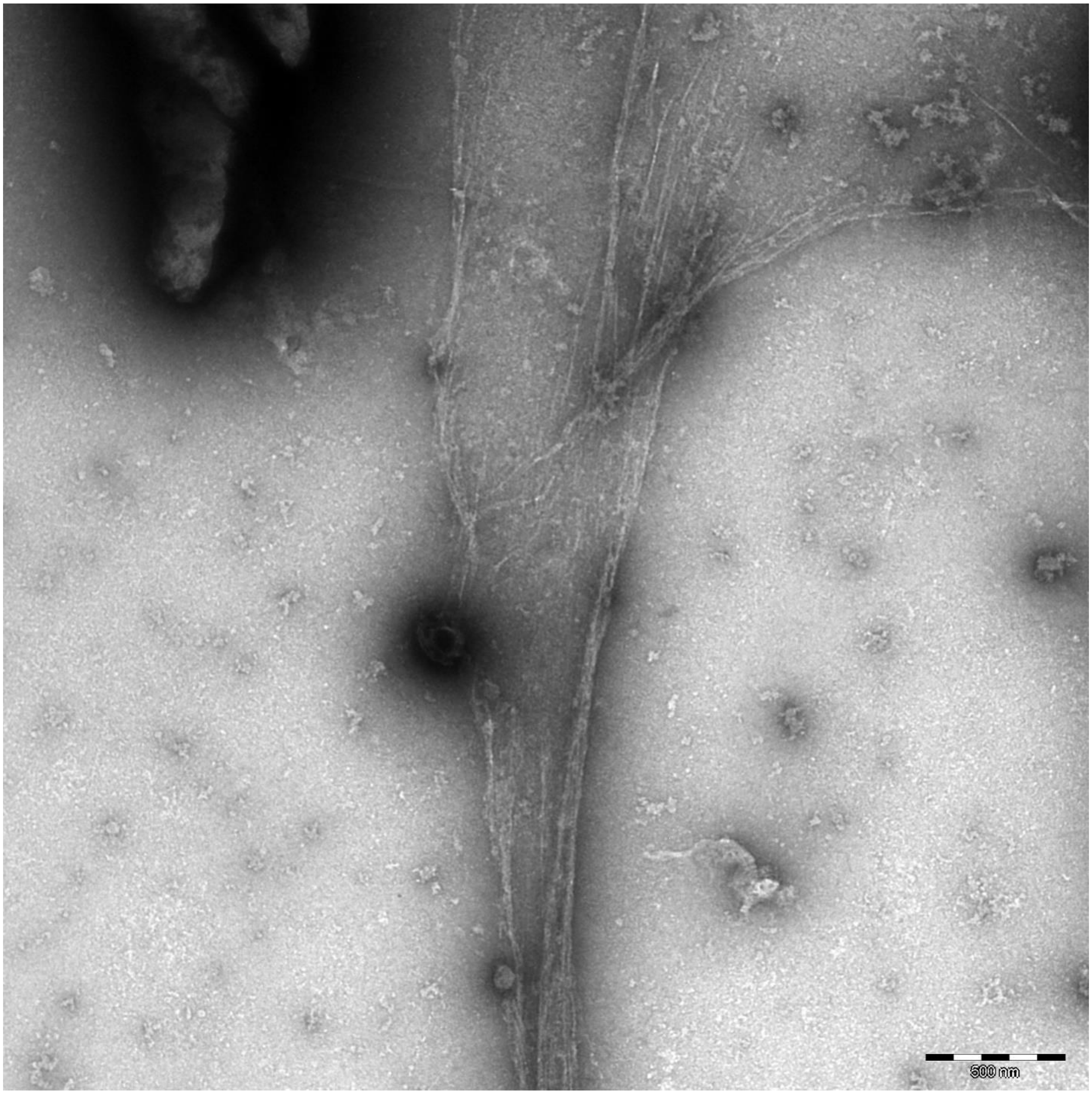
500 nm

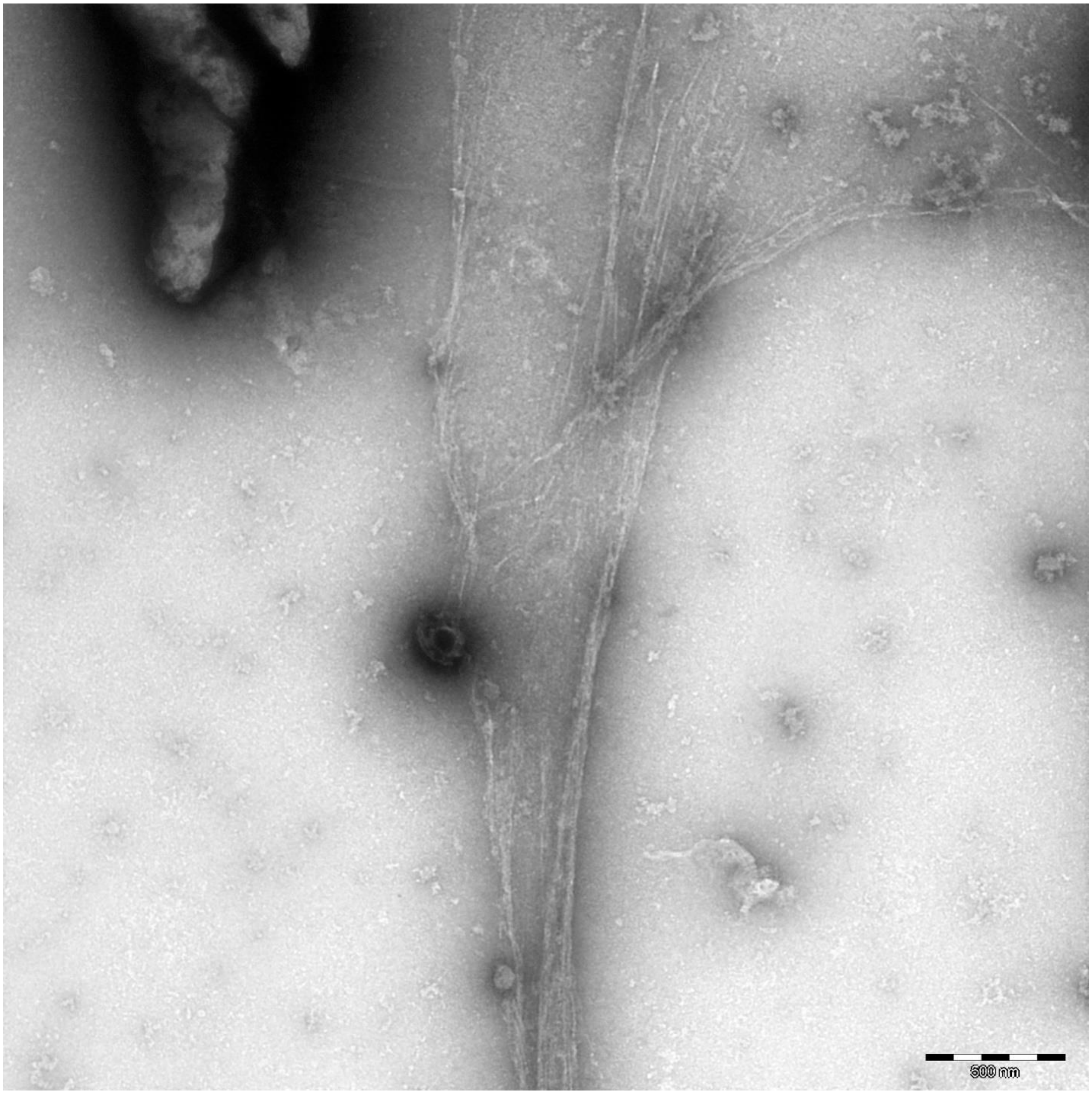
500 nm

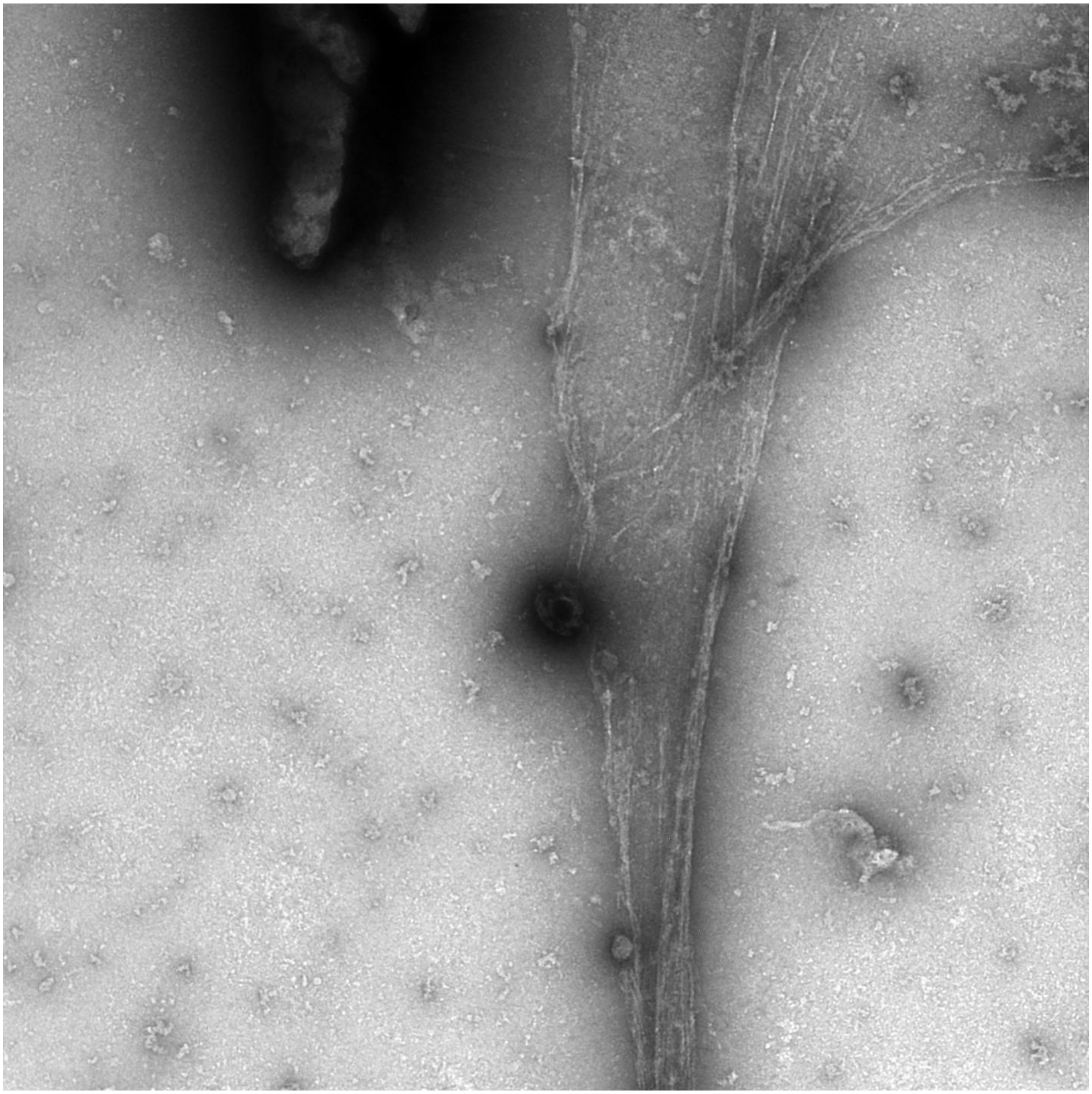

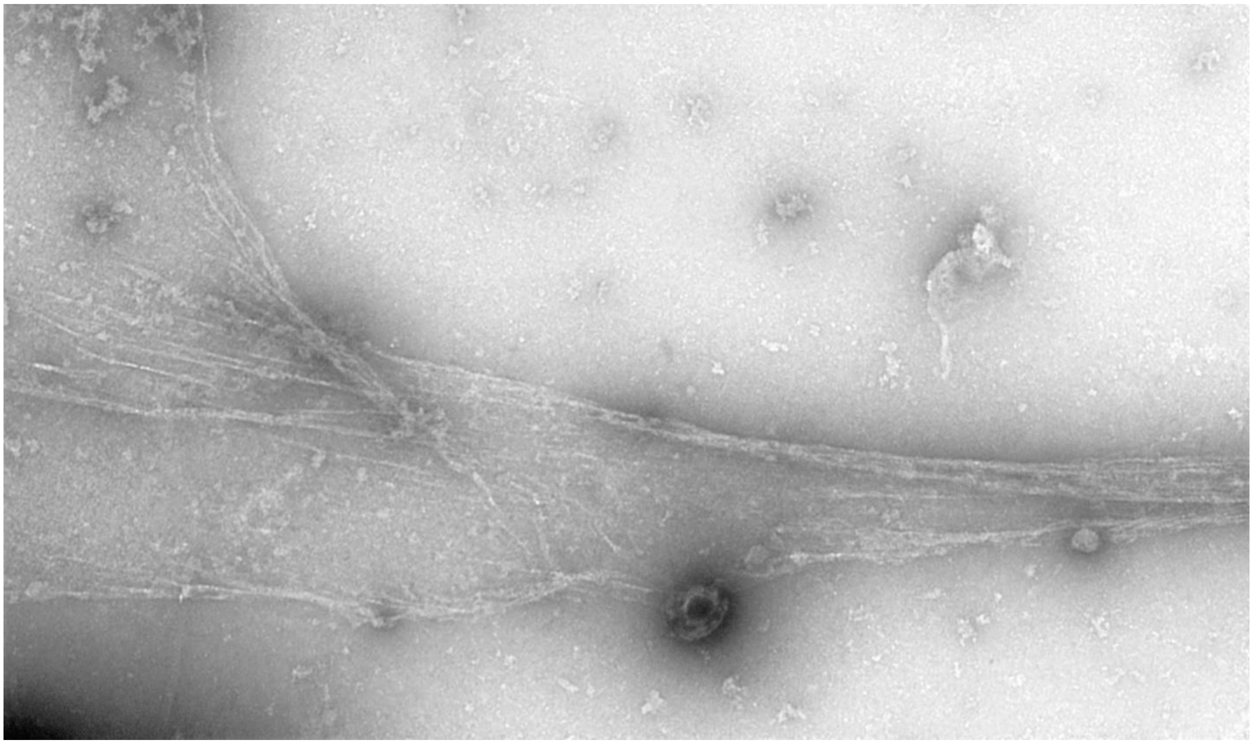

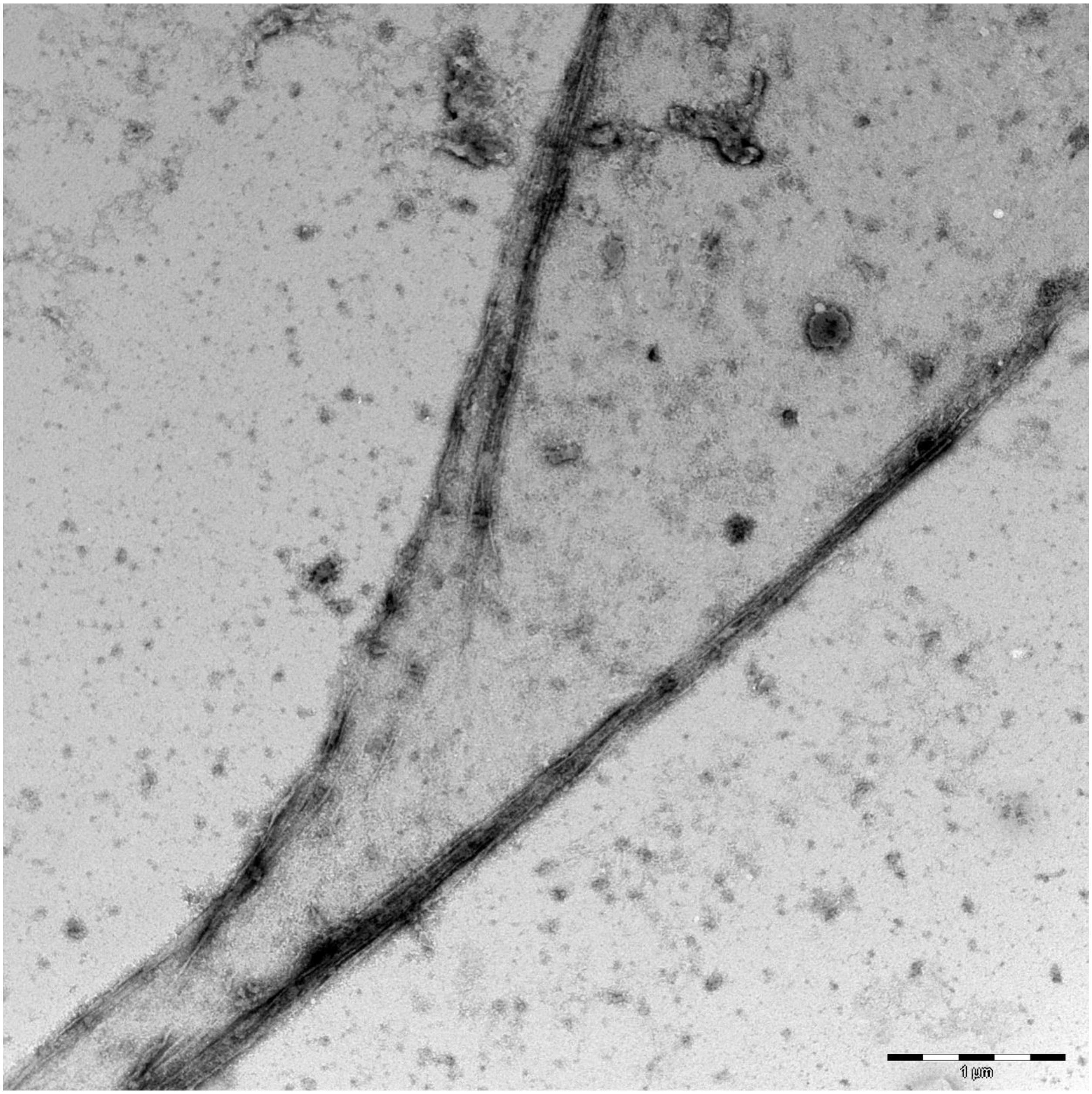
1 µm

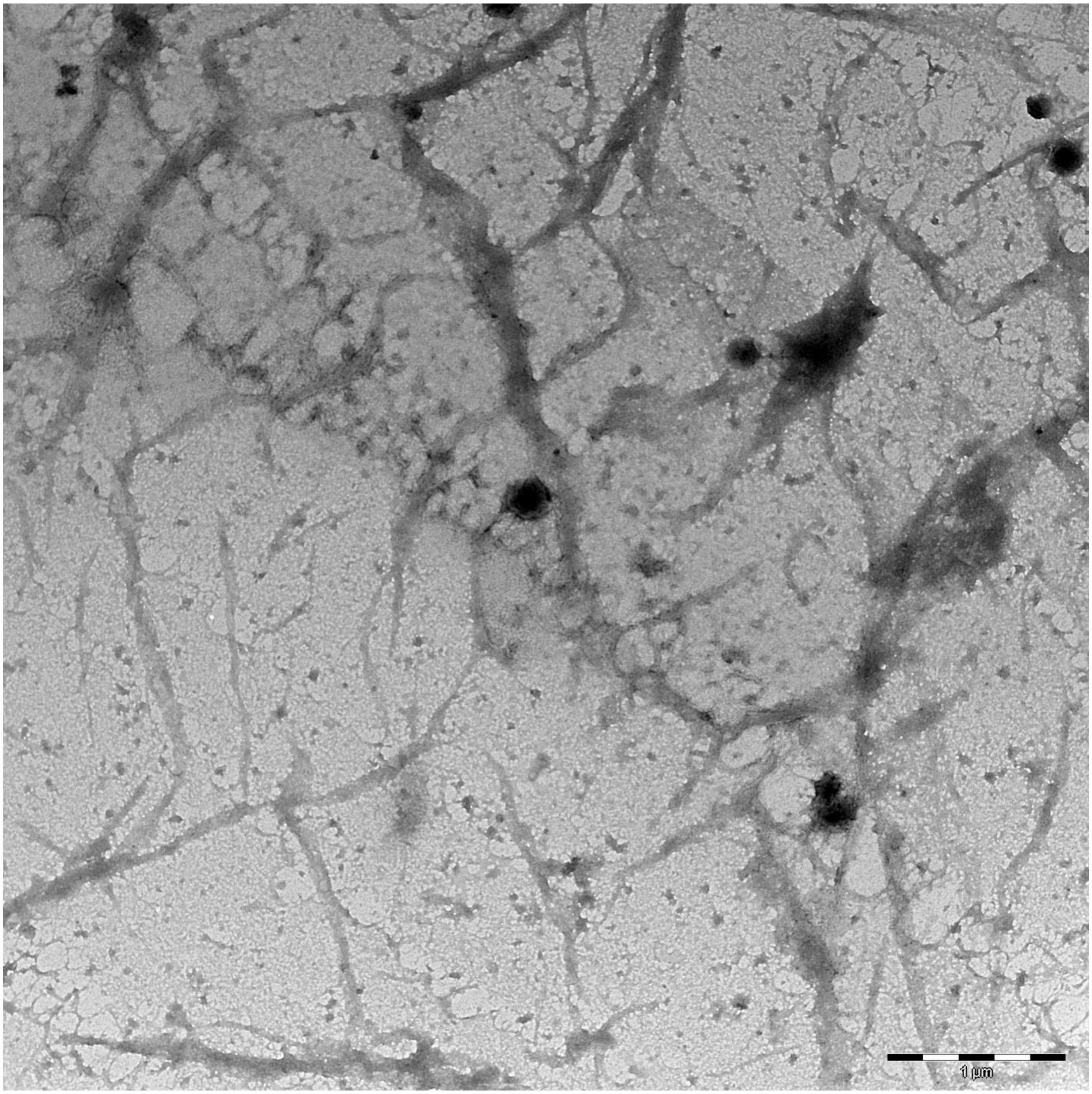
1 µm

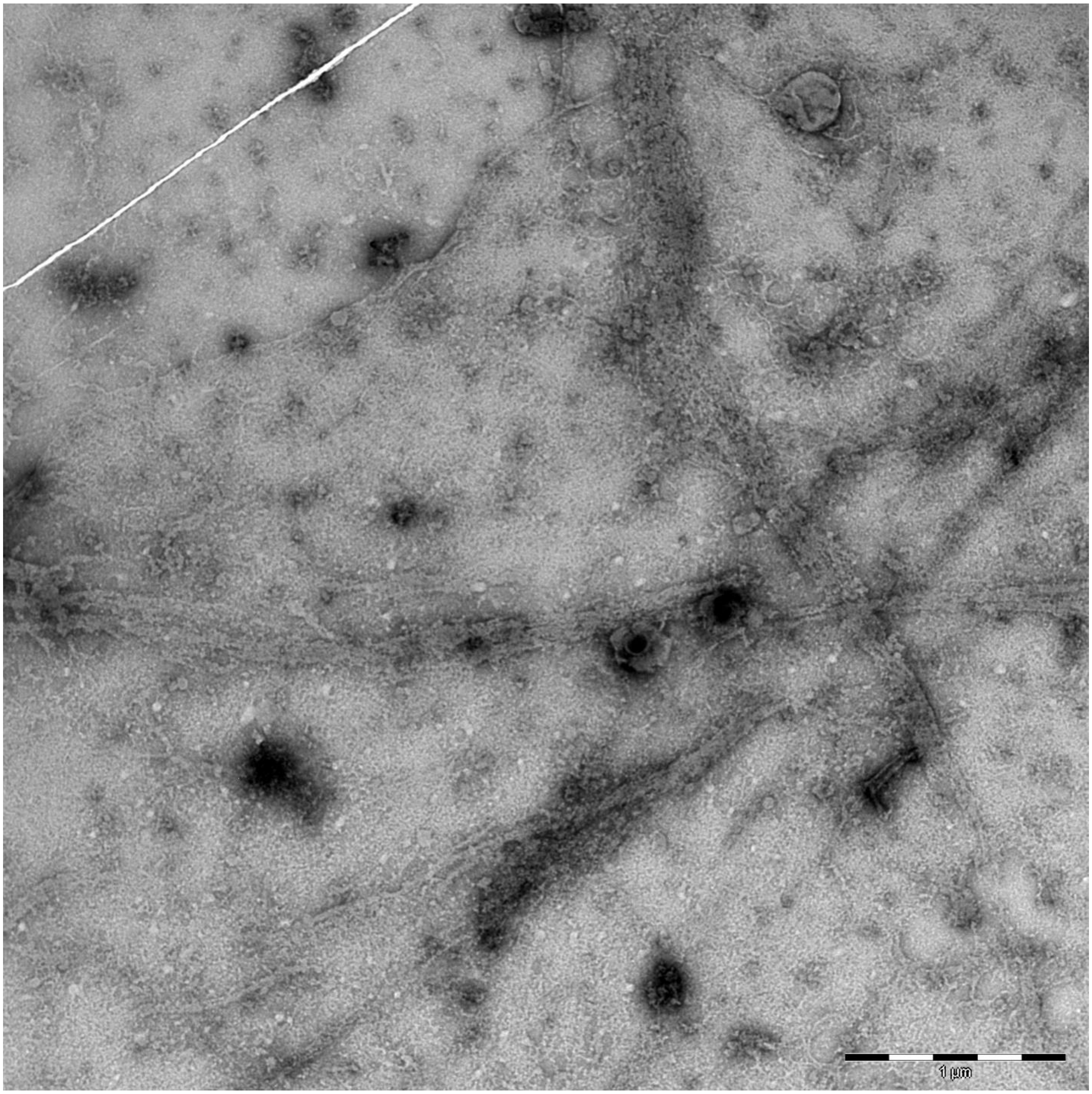
1 µm

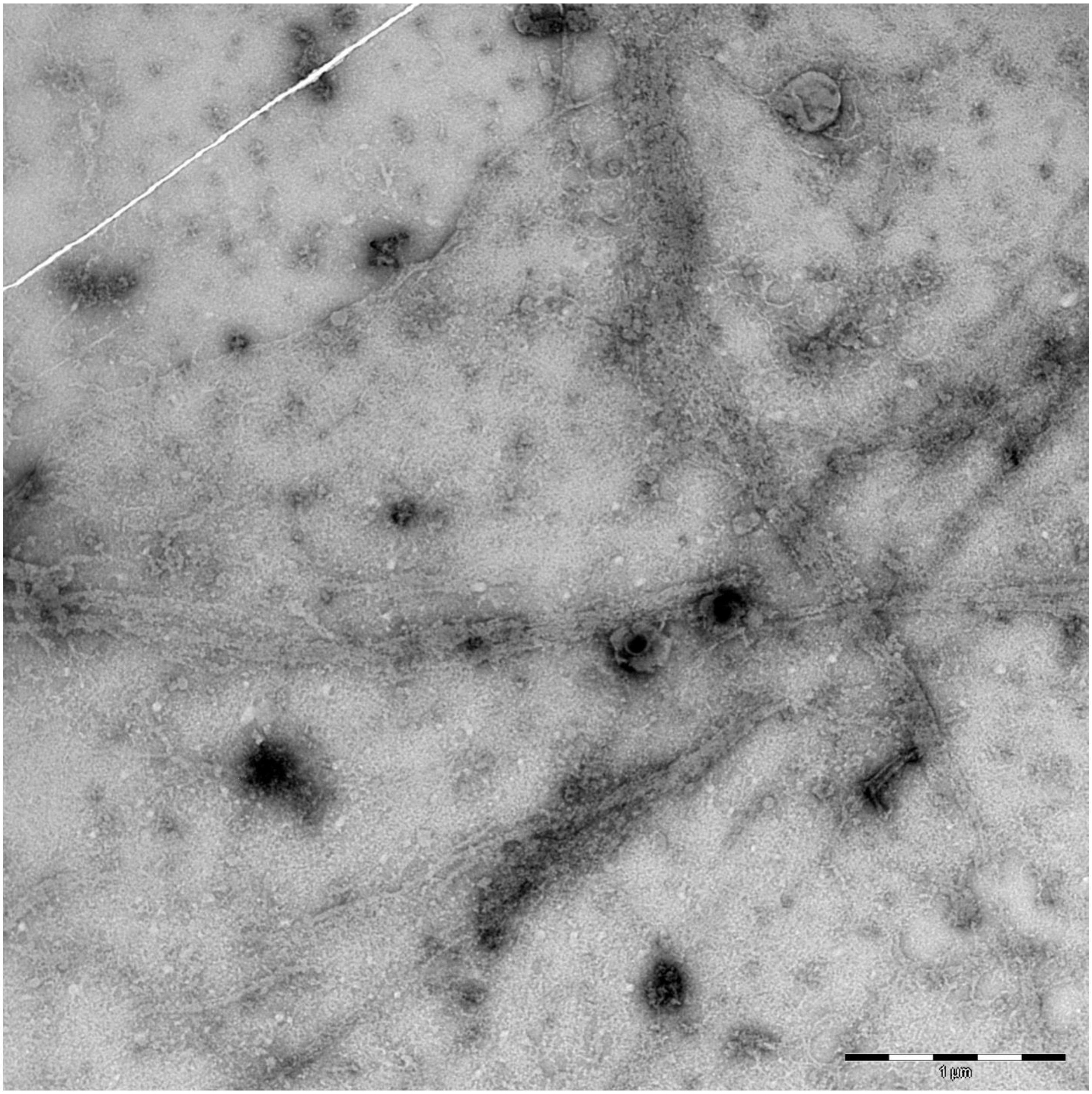
1 µm

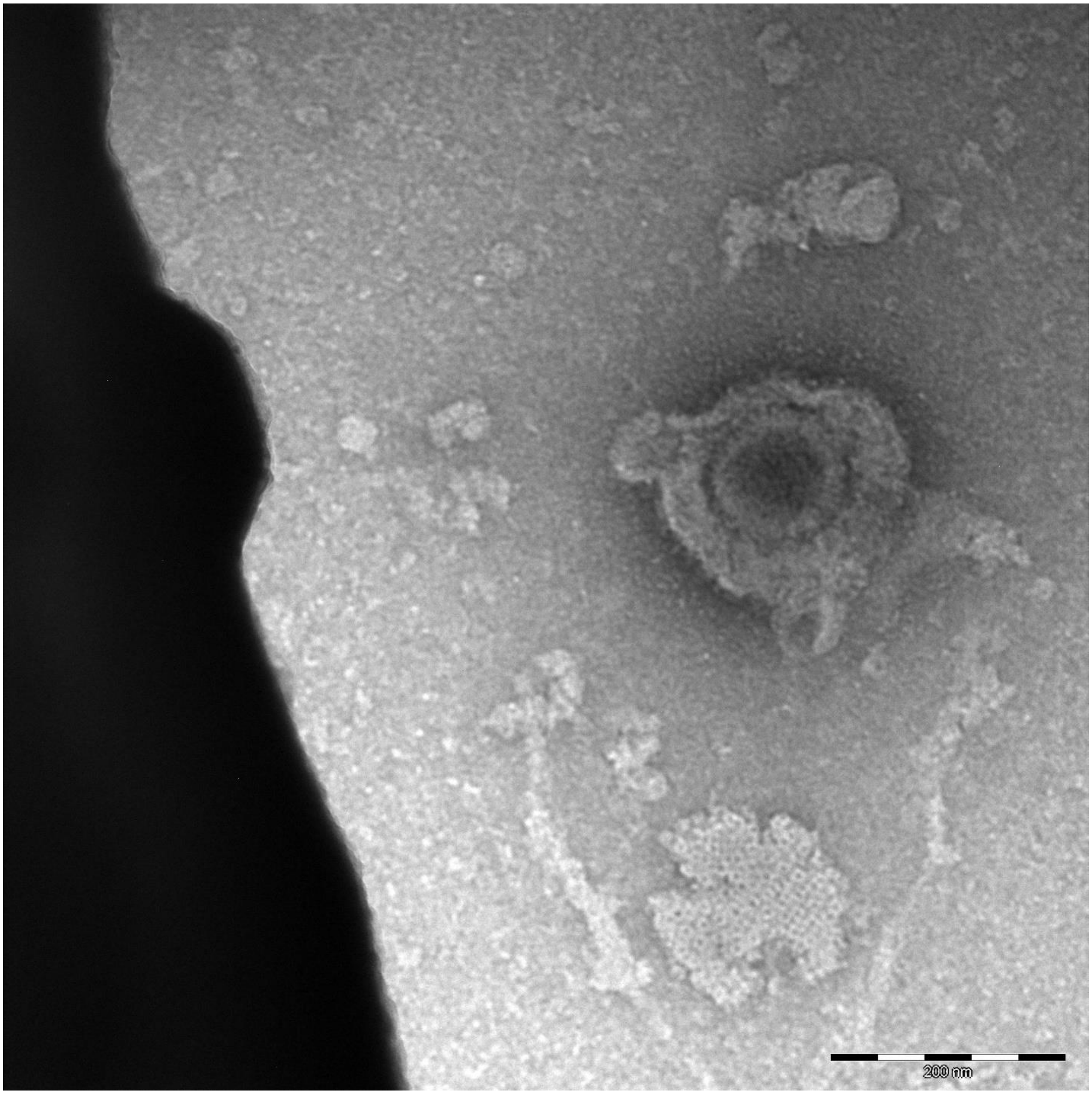
200 nm

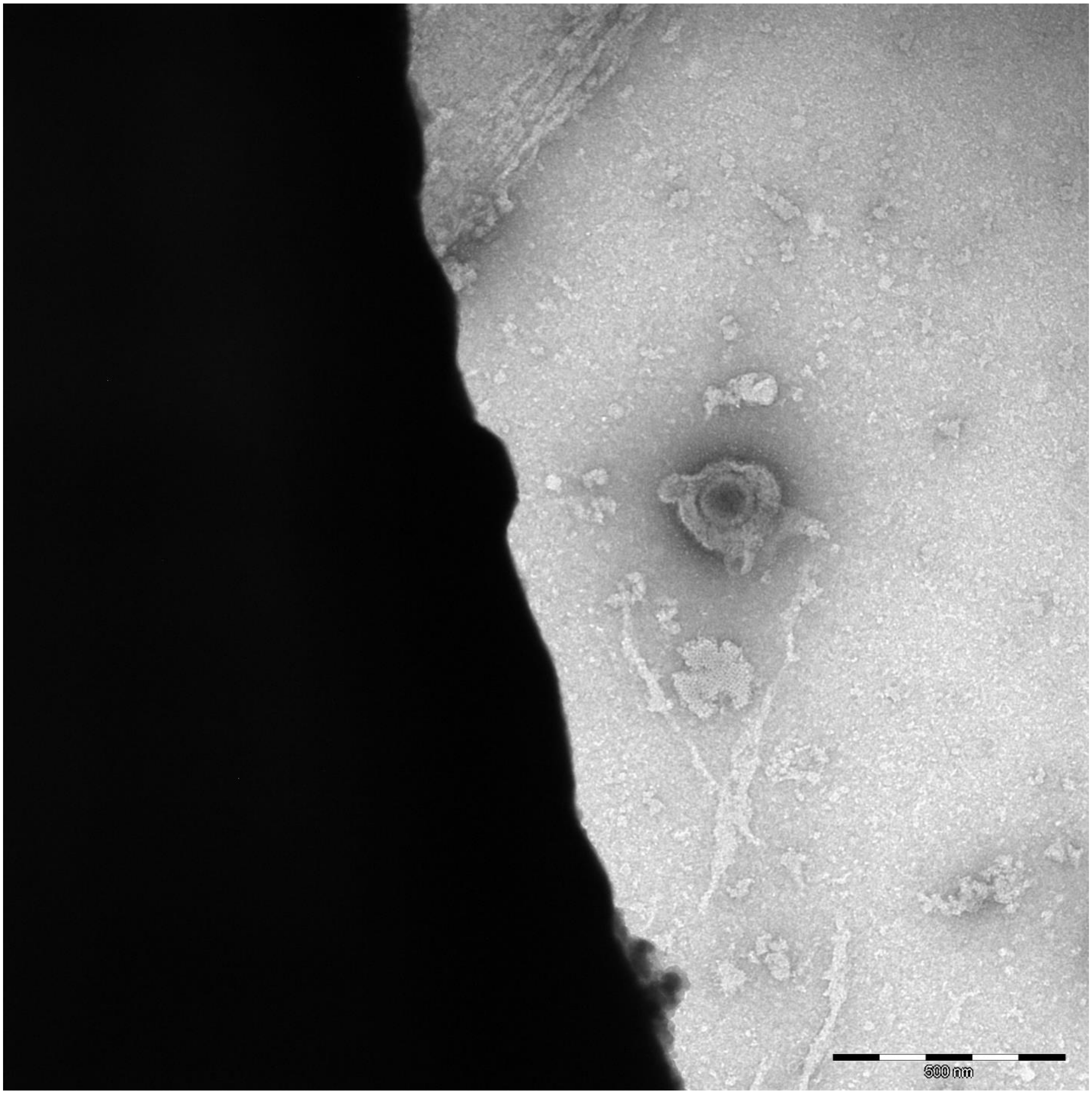
500 nm

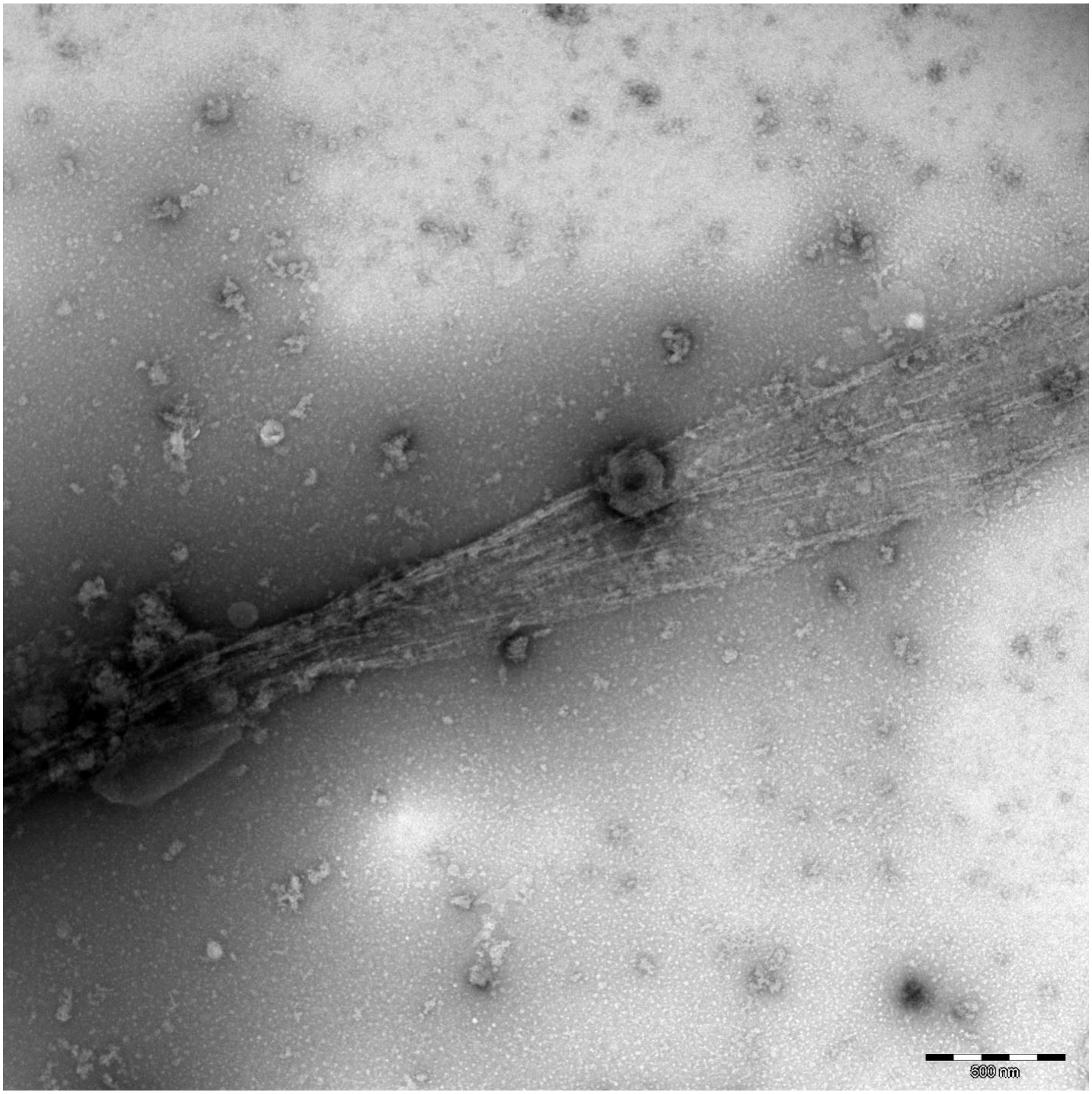
500 nm

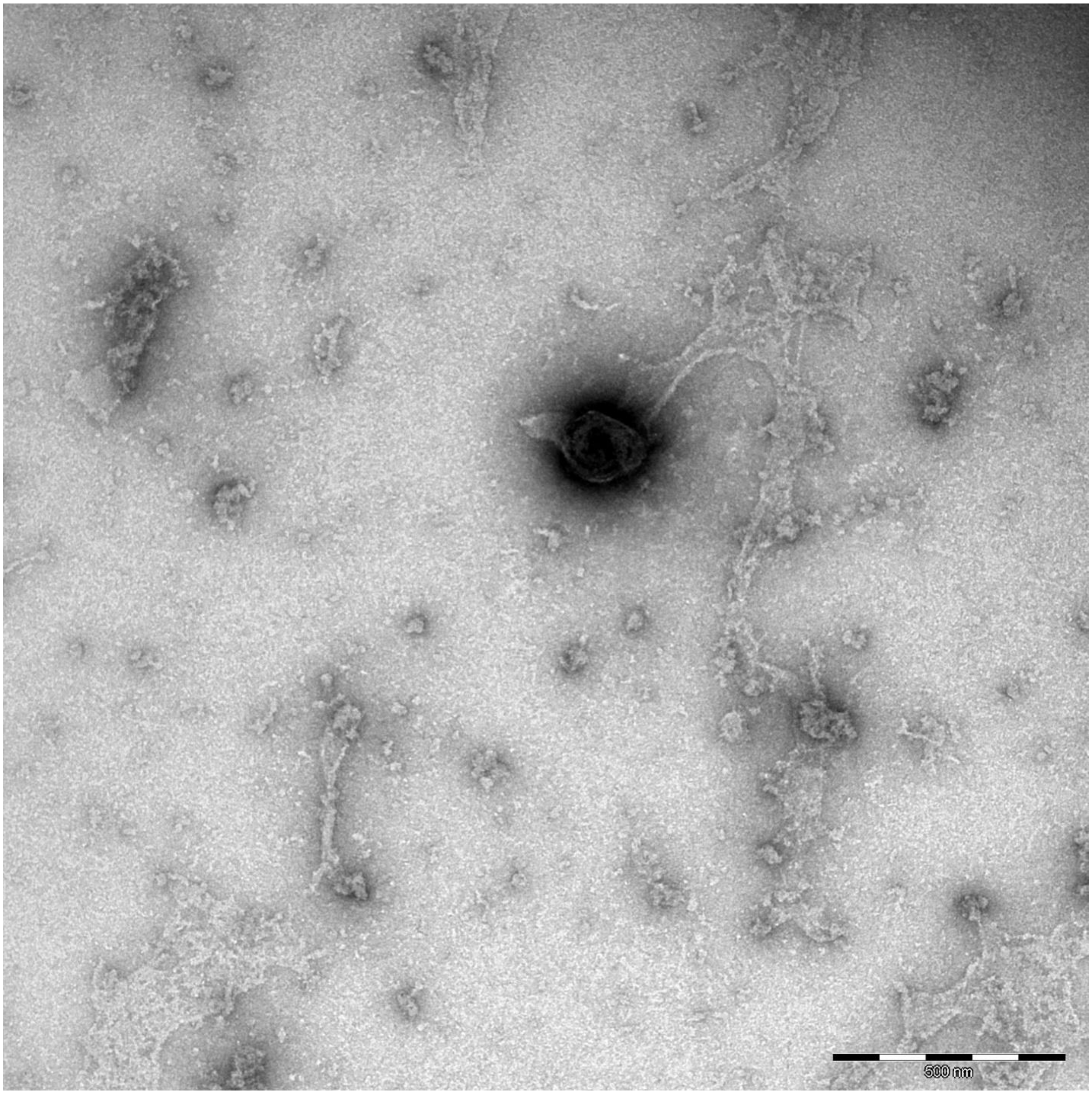
500 nm

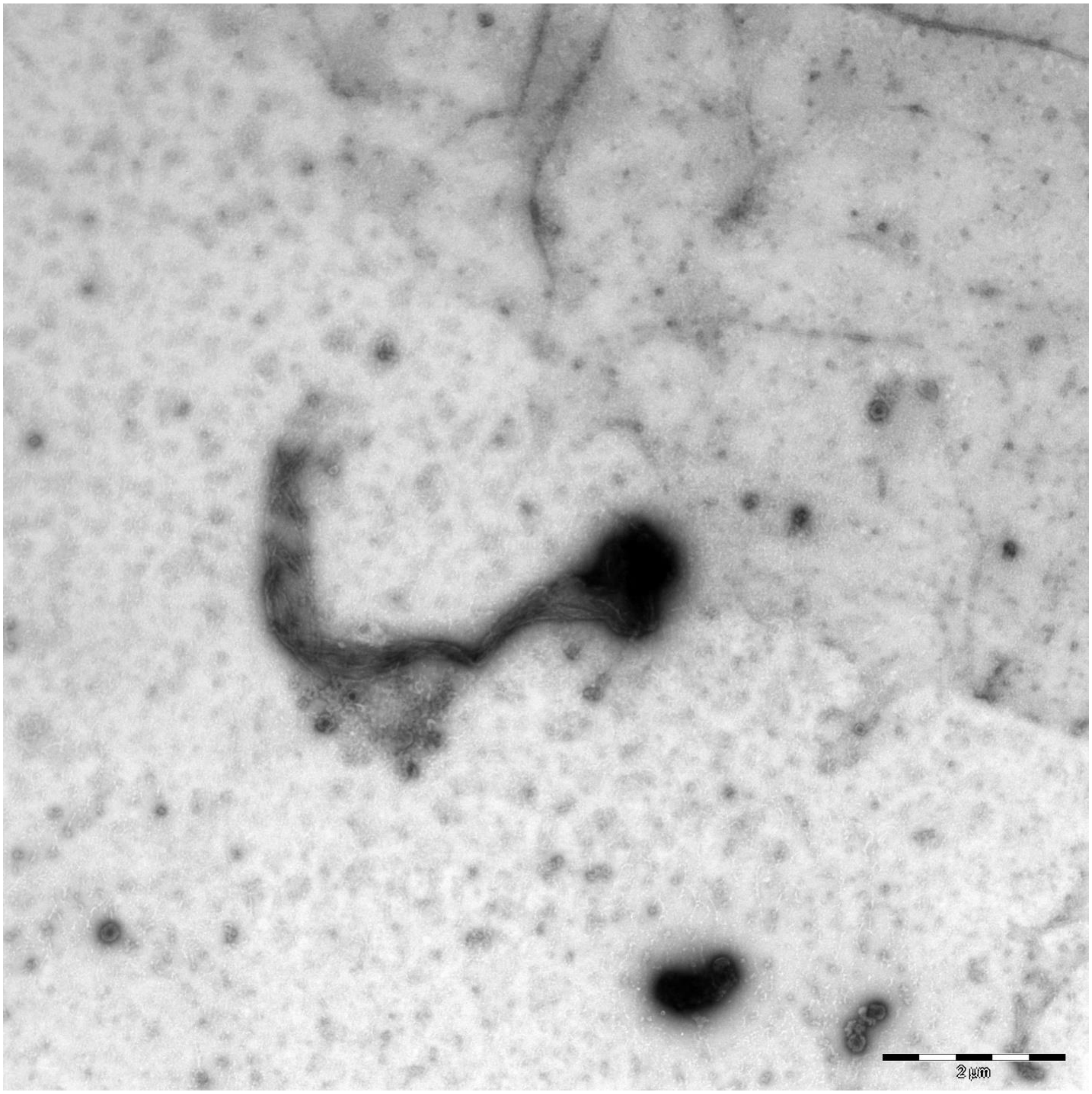
2 µm

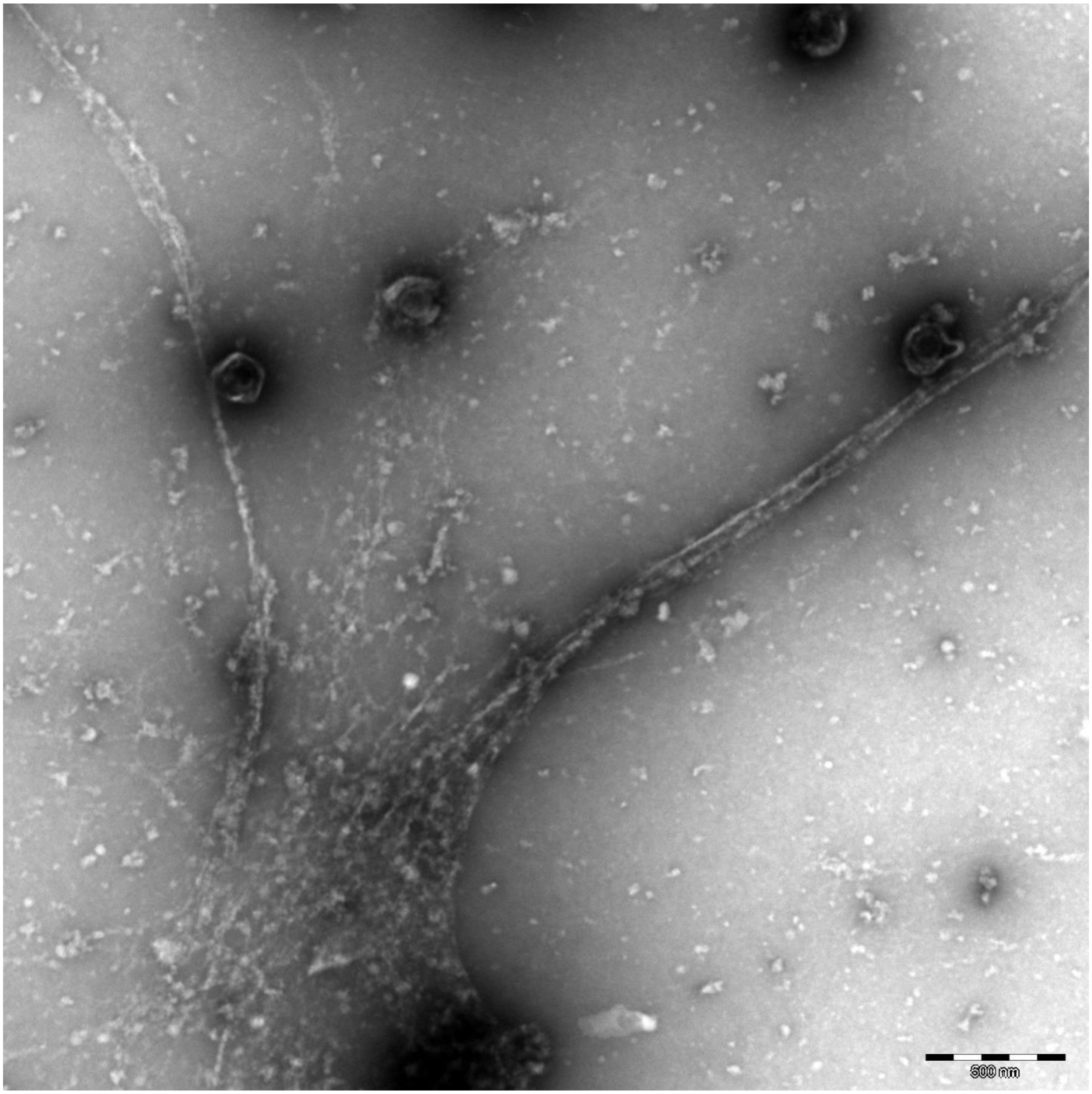
500 nm

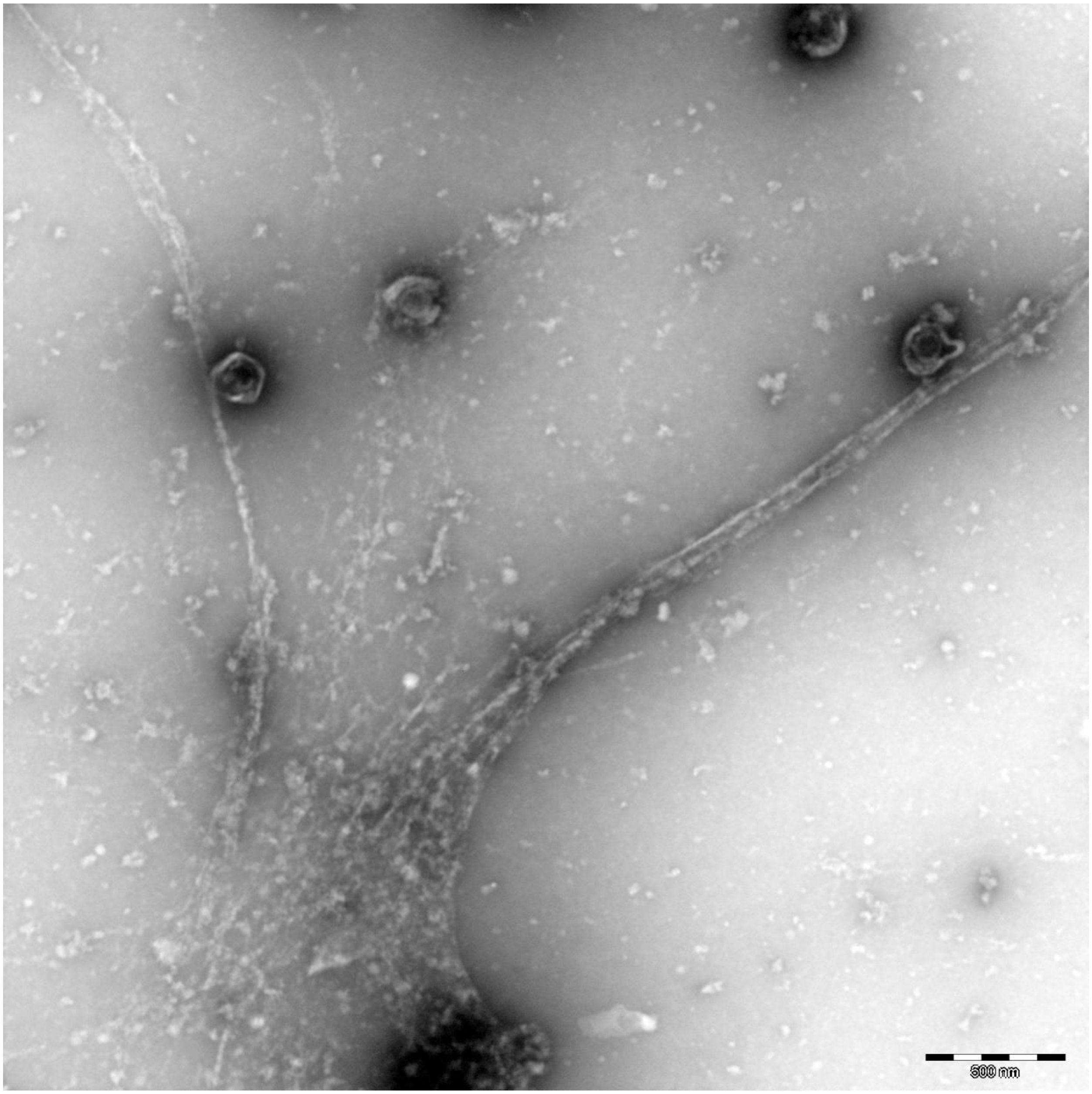
500 nm

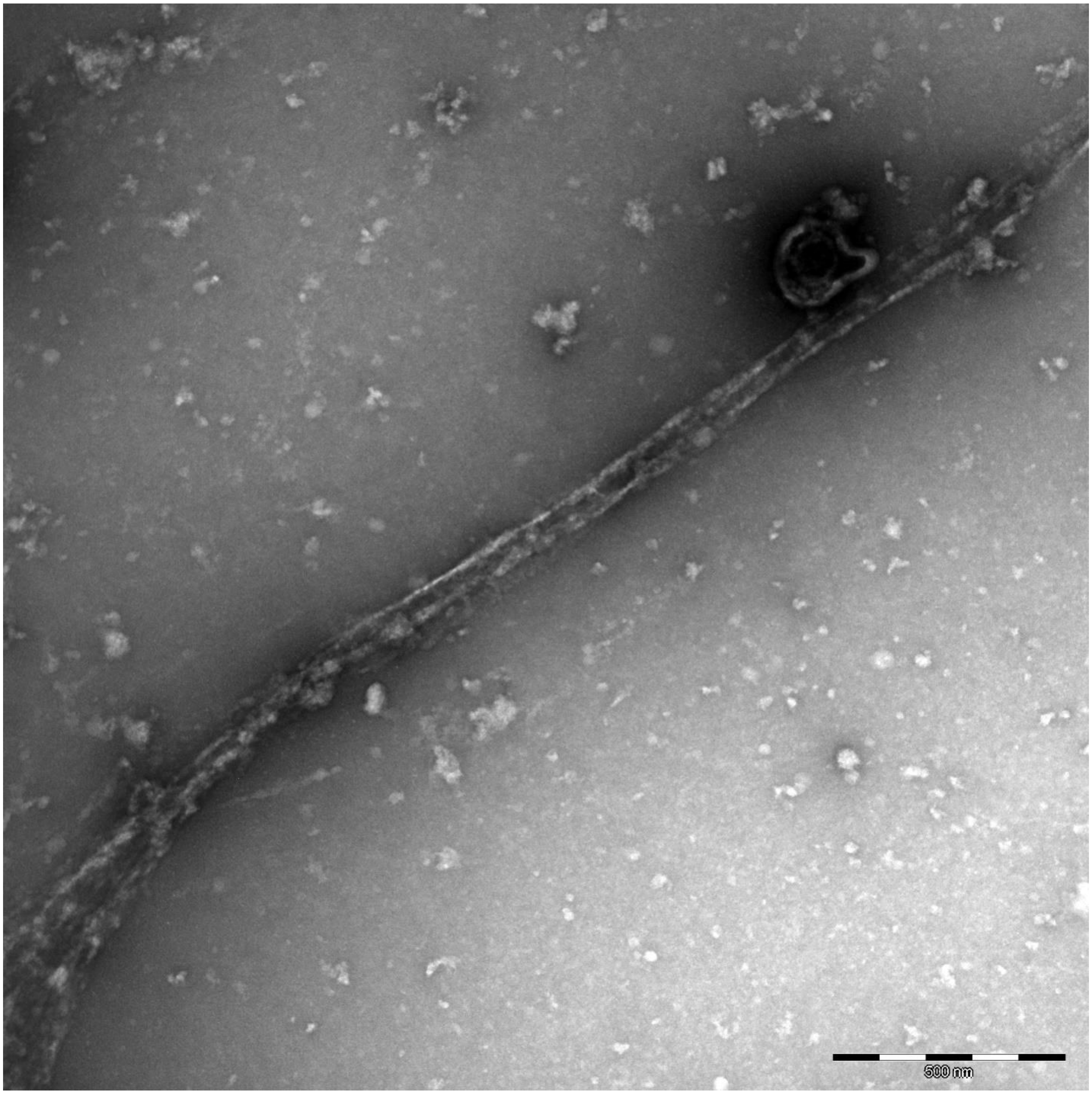
500 nm

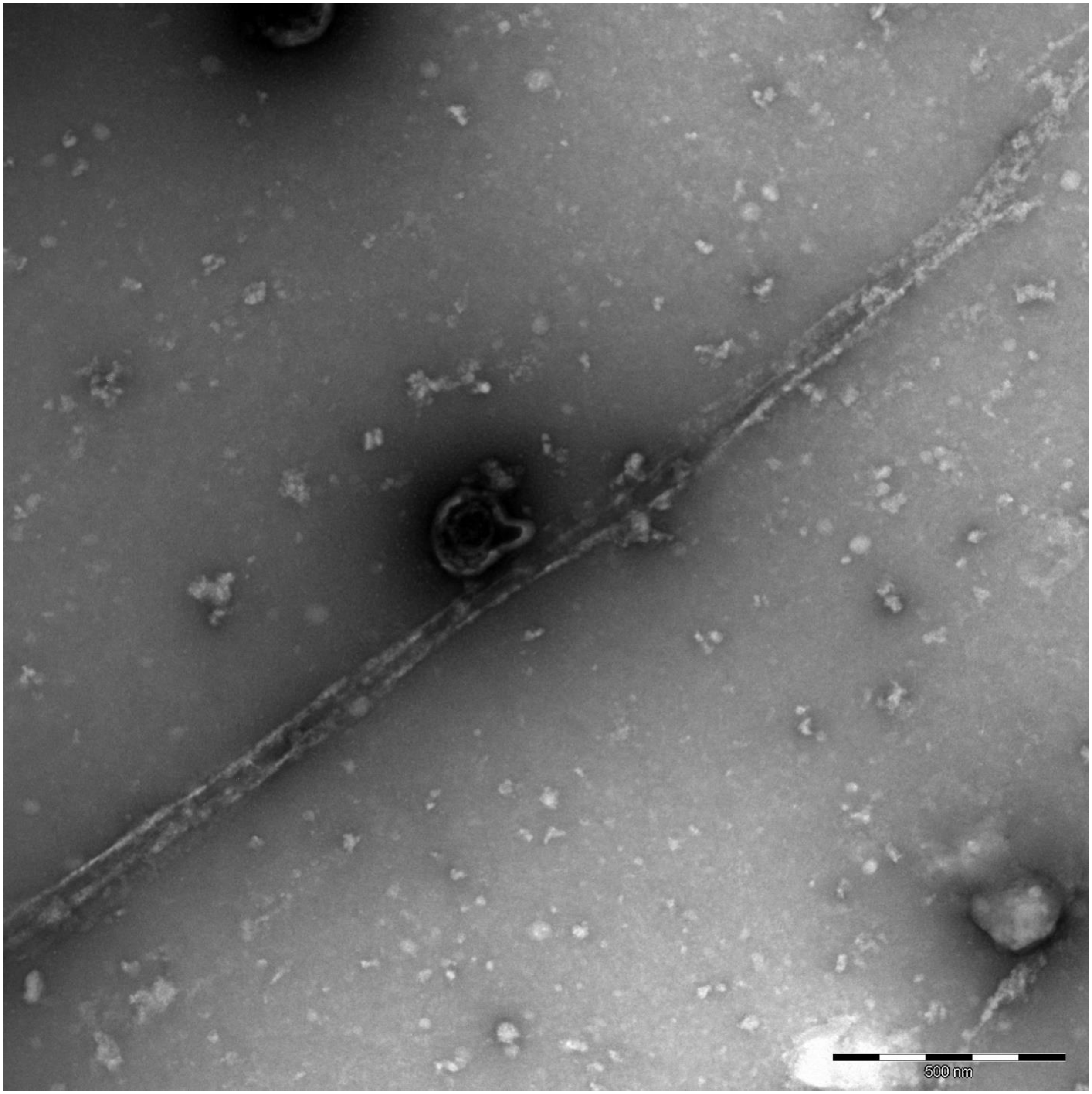
500 nm
