## Supplementary material for "SARS-CoV-2 and HSV-1 Induce Amyloid Aggregation in Human CSF Resulting in Drastic Soluble Protein Depletion": Reviewer comments & responses 1st round

### Reviewer comments and responses: 1st round

Dear Editor,

Thank you very much for reconsidering our manuscript for publication. We also thank the reviewer for their comments, which we think were all addressable and resulted in an improved paper. Here, we present a point-by-point response to the reviewer's comments. Additionally, we have modified the manuscript in accordance with the reviewer's comments and added new experimental data. The responses here and the new additions to the manuscript are highlighted in blue.

**Comment 1:**

Comments on Significance Statement: I could not find it.

**Response:**

This is the significance statement originally uploaded with the manuscript: "Viruses have generally been excluded from the etiology of amyloid pathologies based on the assumption that amyloid formation requires a proteinaceous conformational template (a prion) to form. Here we show that neuroinvasive viruses induce amyloid aggregation of a plethora of proteins in human CSF even after UV inactivation. Our work illustrates that viruses can induce amyloid aggregation of endogenous human proteins in their native environment by acting as physical catalysts of amyloid nucleation and phase transition. Demonstrating this direct mechanistic link, which is independent of templating, can help better understand the link between viruses and neurodegenerative disorders, especially in the post-COVID-19 era."

**Comment 2:**

This work reports the finding that HSV-1 and inactivated SARS-CoV-2 induce protein aggregation in the CSF. The results are interesting, but the conclusions are preliminary and the novelty debatable. It is unclear why the authors did not mention high-profile previous studies that reported that viruses can promote protein aggregation. Specifically, Tanzi, Moir and coworkers (Neuron 2018) demonstrated that HSV-1 triggers amyloid beta aggregation in the brain in a mouse model of Alzheimer's disease. The comparison with this previous study prompts the question of why amyloid beta aggregates were not found in the present work. There are also many related studies about the ability of nanoparticles to trigger protein aggregation (e.g. Linse et al PNAS 2007), which could be discussed.

**Response:**

We have now discussed the previous work mentioned by the reviewer in a revised version of the manuscript, both in the Introduction and Discussion sections. However, to our knowledge, our study is the first to report the induction of amyloid aggregation of proteins in raw human CSF. We agree with the reviewer that induction of amyloid aggregation by viruses and nanoparticles have been shown before (1, 2), including by our lab (3). However, these experiments were performed either with purified proteins at supraphysiological concentrations or utilizing protein overexpression, which make it difficult to conclude how relevant these findings are to a more natural situation in a complex human microenvironment. Using raw human CSF is as close as we can get to studying how these interactions might look like in humans, and it was surprising to us to see amyloid induction in such a natural and complex microenvironment without artificial protein enrichment or overexpression. This is why we think this study brings the field an important step closer to understanding the etiological and mechanistic link between viruses and neurodegenerative diseases in humans. In addition, to our knowledge, no studies before have demonstrated that the SARS-CoV-2 virus particles can act as a physical catalyst of amyloid formation, which we showed resulted in the aggregation of a plethora of important proteins in human CSF. This can be one of the mechanisms leading to some long-COVID symptoms, which is an enduring healthcare crisis after the pandemic.

Furthermore, we took an unbiased proteomic approach to analyze the proteins that were prone to aggregate under these conditions, again reflecting the natural repertoire of vulnerable proteins in human CSF. Moreover, we have now modified Figure 2 to illustrate that a large percentage of the proteins we found in the virus-induced amyloid fractions have also been previously found in amyloid plaques, with a large overlap among the proteins enriched in the amyloid proteome induced by both HSV-1 and SARS-CoV-2 (new Figure 2A, Supplementary Table 2). Such overlap between our findings and previous proteomic studies of amyloid plaques and between the amyloid proteomes induced by different viruses indicates that a subset of neuronal proteins is more vulnerable to aggregation triggers such as viruses, and thus consistently detected in amyloid fractions. Additionally, we added new experimental data (new Figure 2B) showing that viral-mediated amyloid induction leads to a profound depletion of a multitude of soluble proteins from the CSF (please see response to Comment 4).

**Comment 3:**

The authors should demonstrate that 4% SDS is sufficient to separate amyloid aggregates from non-amyloid ones in the CSF samples after incubation with the viral particles. Without this control, the proteins reported in Figure 2 may not be found as amyloid. It is likely that many of these proteins could form amorphous co-aggregates on the surface of the viral particles (known as corona).

**Response:**

Detergent insolubility is the classical way to separate amyloids from other non-amyloid proteins. In fact, we chose SDS and not sarkosyl (another common amyloid enriching detergent) because sarkosyl is less stringent than SDS as it can preserve less robust forms of protein aggregates that cannot withstand SDS treatment (4). Additionally, while some protocols use 2% SDS for amyloid enrichment (5, 6), we used double that amount to make sure we have the highest possible purity in our amyloid fraction. We expanded on the rationale behind our methodological choices in a revised version of the manuscript.

**Comment 4:**

The explanation of the reported findings in terms of supersaturation would require a measurement of the supersaturation concentration from the quantification of the soluble and insoluble fractions in the CSF.

**Response:**

We agree with the reviewer that the explanation in terms of protein supersaturation, while based on extensive work published previously (7–12), is speculative based on the data presented in the manuscript. Thus, we have now removed it from the Results section and discussed it only briefly in the Discussion section. Additionally, we have added an experimental demonstration of the effect of viral-induced amyloid aggregation on the concentration of soluble proteins in the CSF (new Figure 2B). We incubated human CSF with UV-inactivated viral particles for 72 hours at 37 ºC then ran the soluble fraction on a polyacrylamide gel and detected proteins using silver staining. We found that viral incubation with CSF led to a dramatic decrease in the concentration of a large percentage of soluble CSF proteins compared to CSF incubated with medium from non-infected cells. Our results clearly demonstrate that viral incubation and subsequent amyloid induction leads to a substantial decrease in the soluble protein concentration in human CSF. Such big drop is indicative of the ability of viral-induced amyloid aggregation to deplete the CSF of a multitude of functional proteins, which can be a potential pathogenic mechanism via protein loss-of-function.

Taken together, we are confident that the new version of the manuscript has been greatly improved by the reviewer's comments and the newly added experimental data, and that it is now ready for publication in PNAS.

Wish you all the best,
Kariem
