## Supplementary material for "SARS-CoV-2 and HSV-1 Induce Amyloid Aggregation in Human CSF Resulting in Drastic Soluble Protein Depletion": Reviewer comments & responses 2d round

**Reviewer comments and responses: 2nd round**

Reviewer #1:
Suitable Quality?:No
Sufficient General Interest?:No
Conclusions Justified?:No
Clearly Written?:Yes
Procedures Described?:Yes
Supplemental Material Warranted?:Yes
Sufficient data/samples?:No

Comments:
The authors have not convincingly addressed my previous comments. The claims are still overstated:

1) The novelty of studying the catalysis of aggregation induced by surfaces (including virus and nanoparticles) is not novel.

Response:

Viral-induced aggregation of human CSF proteins has never been reported before.

2) The study of aggregation in the CSF may be described as physiological, but it is still not in the brain.

Response:

Using human CSF is as close as we can get to physiological human conditions. Unfortunately, it is not possible for ethical reasons do this in human brains as the reviewer suggested.

3) There is still no direct evidence that the aggregates are amyloid.

Response:

We used standard methodology to show amyloid formation including direct visualization of the amyloid fibrils with electron microscopy. We now include all our raw electron microcopy images in new Supplementary Figures 2 and 3, which show abundant amyloid formation very clearly.

Reviewer #2:
Suitable Quality?:No
Sufficient General Interest?:No
Conclusions Justified?:No
Clearly Written?:Yes
Procedures Described?:No
Supplemental Material Warranted?:No
Sufficient data/samples?:No

Comments:
The topic of this manuscript and the questions the authors pose are very interesting. The idea to study virus induced aggregation in CSF as a representation of the natural environment of the viruses is original and may be effective. However, methodologically the presented work has serious shortcomings and the

analysis seems rather superficial.

Below I list a few of my concerns with the presented experimental data.

ThT aggregation assays:

1. According to the materials and methods the ThT aggregation assays were performed at a ThT concentration of 3mM. This is extremely high, typically the experiments are done at concentrations that are 1000x lower than this. The reasons for this is that 1) ThT affects the aggregation process itself, therefore ThT concentrations are used are typically as low as possible. 2) The solubility of ThT in pure water water is ~9mM, with the addition of salt the solvent quality decreases and the solubility will drop. I therefore wonder if the authors are not just looking at the formation of fluorescent ThT aggregates with the addition of the (very salty) CSF.

The controls were performed by substituting CSF with MQ water. The controls should have been performed in artificial CSF or a buffer with equivalent ionic strength since salt concentration is an important factor in the aggregation process.

Response:

All the samples represented contain ThT, including the samples labelled HSV-1 only, SARS-CoV-2 only and CSF only. ThT + CSF is the control for background signal in all the figures, including Figure 1. (brown line with minimal signal at the bottom). This control was also statistically compared to the viral induced signals in Supplementary Figure 1. The data in these figures clearly show that mixing ThT at the stated concentration with human CSF does not lead to a significant amyloid signal by itself. In this regard, this is a more relevant and stringent control than using artificial CSF which only contain salts mimicking CSF ionic strength but lacks any proteins. Moreover, we have an additional control which is Noninfected medium+ CSF, which also contains ThT at the same concentration. As shown in Supplementary Figure 1, the viral signal is significantly higher than this control, which in addition to having CSF with all its constituents, it also has all the constituents of a cell growth medium except for viruses. Based on these multiple controls, we are confident that the significantly higher signal in the virus-containing samples is due to viral-induced amyloid aggregation and not due to unspecific interaction between the dye and the CSF.

2. To really show that viruses physically induce aggregation a concentration dependent effect on the time to aggregation should have been included.

Response:

We have performed this experiment (new Supplementary Figure 2), where we demonstrate that concentrating that 3x concentrated HSV-1 viral stock results in a significantly higher amyloid signal with shorter time to aggregation.

TEM images:

3. The TEM images are obtained at different virus/CSF ratios than the presented ThT data

Response:

Yes, we increased the ratio to make sure we produce enough amyloid fibrils for visualization.

4. The virus controls in artificial CSF are missing.

Response:

Please see response to comment no. 1.

5. Besides the electron-dense structures that the authors assign to the viruses and the elongated fibrils, the images also contain a large portion of spherical objects (~30 nm). The authors do not mention these.

Response:

We use viruses harvested from cell supernatant, other smaller round structures could be exosomes or empty viral particles. Additionally, we now include all our raw electron microcopy images in new supplementary figures 2 and 3, which show abundant amyloid formation very clearly.

Proteomic analysis
5. Once precipitated proteins are very difficult to redissolve. The authors do not provide any convincing evidence that the precipitated proteins are amyloid, 4% SDS insoluble is not evidencing this, the precipitate could just as well (more likely) be an amorphous aggregate.

Response:

EM clearly shows non-amorphous amyloid fibrils. We now include all our raw electron microcopy images in new Supplementary Figures 2 and 3, which show abundant amyloid formation very clearly.

6. The authors only show the final result of a probably elaborate analysis, no real data is shown, nothing is mentioned on confidence or abundance.

Response:

We have a detailed description of the analysis it in the Methods section and more details are available upon request.

Gels
7. The controls showing CSF and virus in artificial CSF are missing.

Response:

Please see response to comment no. 1.

8. Specific proteins seem to disappear from the supernatant ending up in the pellet. Can the authors assign any of these based on e.g. their proteomic analysis and their known molecular weight

Response:

This is a valid point is valid and we will follow up on the missing proteins with more analysis.
